# COTree: A Statistical Framework for Deciphering Cell-Resolved Multi-Omics Trajectories

**DOI:** 10.1101/2025.10.30.685684

**Authors:** Bo Yuan, Rong Wei, Troy Brier, Enguang Fu, Zane Thornburg, Zaida Luthey-Schulten, Shulei Wang

## Abstract

Recent advances in whole-cell modeling enable the computational tracking of the temporal evolution of thousands of molecular species across genomic, transcriptomic, proteomic, and metabolomic layers. These models provide a complementary perspective for studying cellular dynamics, offering continuous, system-wide observations that are difficult to obtain from experimental technologies, which are often destructive and yield only static measurements from limited modalities. While whole-cell models generate multi-omic simulation trajectories with high temporal resolution, analyzing and interpreting such complex data remains a major challenge that limits their potential to elucidate cellular dynamics. To address this challenge, we propose COTree, a statistical framework that learns integrated multi-omic representations and constructs a trajectory principal tree to summarize cellular progression patterns. COTree enables a broad range of downstream analyses, including cell classification, fate prediction, developmental time detection, and driver species identification, that provide new insights into how cells develop and differentiate. To demonstrate its practical utility, we apply COTree to a multi-omic trajectory dataset generated from the whole-cell model of JCVI-Syn3A, revealing cell types, characterizing long-term cellular dynamics, and identifying key driver species associated with cell death and replication.

## 1. Introduction

Understanding cellular processes such as cell-cycle progression, differentiation, and migration is fundamental to elucidating how cells develop and adapt to their environments. These processes arise from complex, multimodal molecular interactions across genomic, proteomic, and metabolomic layers, which introduce variability in cell fate and give rise to distinct developmental trajectories (Bengtsson et al., 2005; Raj and van Oudenaarden, 2009). Characterizing how these interactions and their dynamics evolve is therefore essential for uncovering the principles that govern cellular behavior.

Recent advances in whole-cell modeling have enabled an integrative framework for studying cellular dynamics (Karr et al., 2012; Hutchison et al., 2016; Breuer et al., 2019; Thornburg et al., 2022; Wu et al., 2025; Fu et al., 2025; Thornburg et al., 2026). Building on decades of biochemical, single-molecule, and spectroscopic characterization of reaction kinetics, whole-cell models computationally track the temporal evolution of thousands of molecular species and intermediates across genomic, transcriptomic, proteomic, and metabolomic layers (Bremer and Dennis, 2008; Taniguchi et al., 2010; Park et al., 2016). Compared with current experimental technologies, which are often destructive and provide only snapshot measurements from limited modalities, the holistic view offered by whole-cell models reveals how the interplay among metabolism, genetic information processing, and growth drives changes in a cell’s behavior over time.

The equations and simulation rules within a whole-cell model specify cellular dynamics at the level of individual reactions or biological processes, whereas their integration gives rise to system-level behavior over an entire cell cycle. For stochastic whole-cell models such as the JCVI-syn3A model, repeated simulations can be viewed as realizations from an implicit distribution over high-dimensional, multimodal cellular trajectories (Karr et al., 2012; Thornburg et al., 2022). Although the generative mechanisms are specified, the dominant progression patterns, cell-to-cell heterogeneity, and alternative cellular outcomes induced by these mechanisms are not directly apparent from individual model components or raw simulation output. Statistical analysis is therefore needed to determine how early molecular variability propagates across omics layers, when cellular trajectories diverge, and which coordinated molecular changes are associated with different outcomes.

Most trajectory analysis methods in computational biology aim to reconstruct the progression order of gene-expression changes from single-cell snapshot measurements (Trapnell et al., 2014; Trapnell, 2015; Cannoodt, Saelens and Saeys, 2016; Street et al., 2018; Saelens et al., 2019; Hou et al., 2023) or to infer the direction of transcriptomic change using RNA-velocity or related approaches (Bergen et al., 2020; Lange et al., 2022; Li et al., 2023). While such analyses have provided valuable insights into cellular dynamics from cross-sectional transcriptomic data, they are not directly applicable to multi-omic trajectories, as they assume static measurements and overlook the explicit temporal continuity and cross-omics coupling present in whole-cell model simulations (Welch, Hartemink and Prins, 2017; Arge-laguet et al., 2020). To realize the full potential of whole-cell models in deciphering cellular dynamics, a dedicated statistical framework tailored to the analysis and interpretation of high-dimensional, multi-omic, and time-resolved data is needed.

To address this need, we propose COTree, a two-stage statistical framework for analyzing cell-resolved multi-omics trajectory data. In the first stage, COTree learns integrated low-dimensional representations using a novel dual-target multi-omics autoencoder, which combines signals across omic layers and captures both continuous growth trends and transient bursts in cellular processes. In the second stage, COTree employs a time-informed principal tree estimation algorithm to construct a trajectory tree that summarizes collective cellular dynamics in the learned representation space. Building on this representation and trajectory tree, COTree supports a wide range of downstream analyses, including cell classification, fate prediction, developmental-time estimation, and driver-species identification. We demonstrate its utility using a whole-cell model dataset (Section 2) that records per-minute molecular abundances across an entire cell cycle. The analysis delineates cell types, characterizes long-term cellular dynamics, and identifies key driver species associated with cell death and replication.

## 2. Multi-Omics Trajectories from a Whole-Cell Model Dataset

The cellular trajectory dataset used to illustrate our new framework was simulated using the fully dynamical whole-cell kinetic model of JCVI-syn3A, a minimal cell with a reduced genome comprising 493 genes (Thornburg et al., 2022). This model captures the integrated dynamics of cellular metabolism, genetic information processing, and growth using a hybrid deterministic-stochastic modeling approach. In particular, essential metabolic networks are described by ordinary differential equations, whereas genetic information processes,including DNA replication, transcription, translation, mRNA degradation, and tRNA charging, are simulated using a chemical master equation framework.

The dataset consists of 207 simulated, cell-resolved multi-omic trajectories under the well-stirred model (Figure 1). For each cell, molecular counts were recorded for 493 genes, 455 mRNA transcripts, and 454 protein products (further differentiated to 551 entries based on chemical and spatial state), together with 236 metabolites participating in 175 metabolic reactions, sampled every minute throughout a complete cell cycle (105 minutes). In the statistical analysis, we excluded metabolite species with constant values and protein species corresponding to cytoplasmic forms of membrane proteins, leaving 181 metabolites and 458 proteins for analysis. Among the 207 cells, 127 cells underwent multiple DNA replication events, 50 cells experienced a single replication event, and 30 cells exhibited glycolytic arrest due to phosphoenolpyruvate (PEP) depletion.

**FIG 1.**
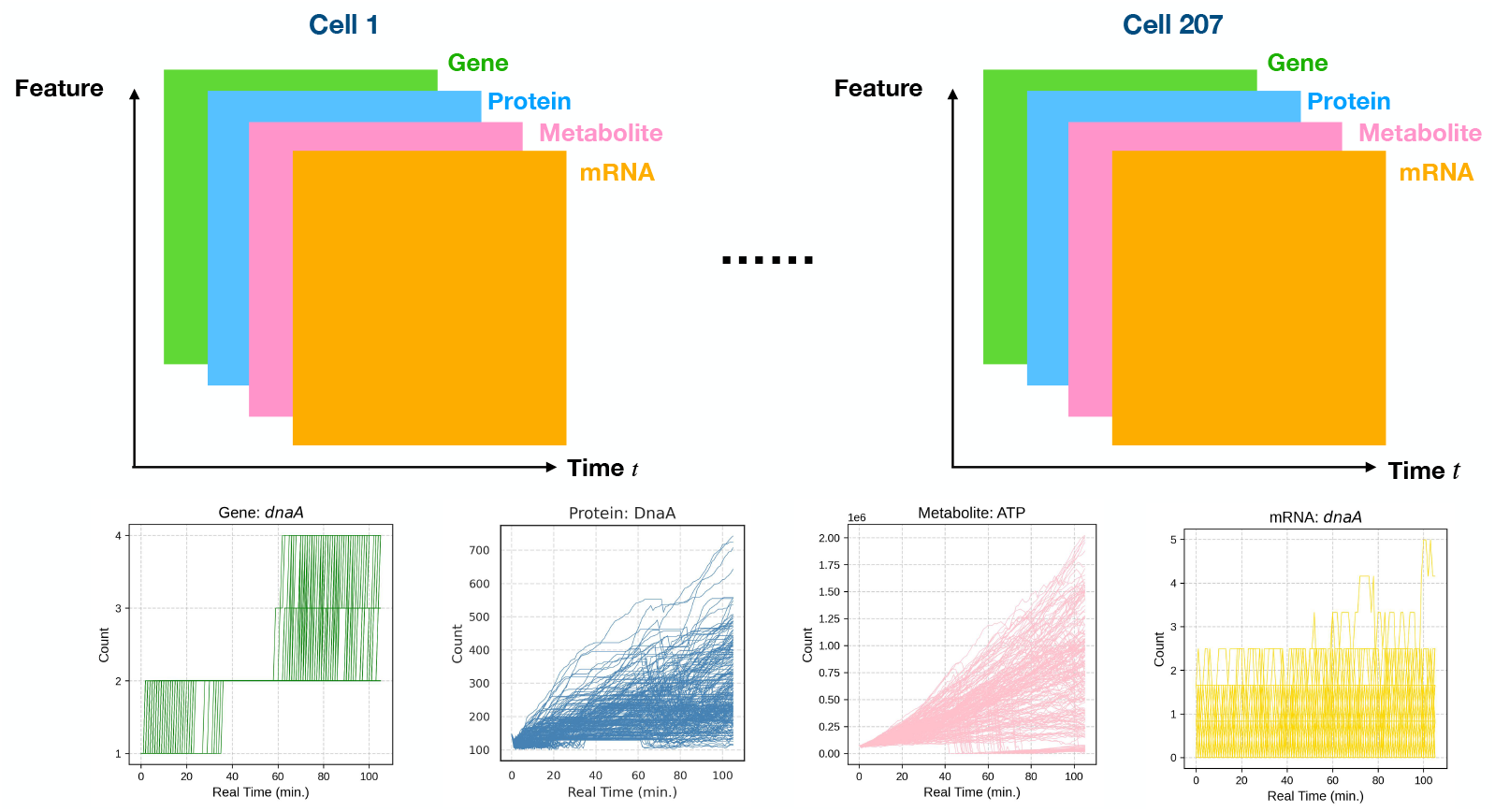
Dataset from the whole-cell model. The whole-cell model (WCM) dataset consists of 207 simulated cells. For each cell, per-minute abundances of molecular species from four omics layers, genomics, transcriptomics, proteomics, and metabolomics, are recorded over an entire cell cycle. Representative species from genes, mRNA transcripts, proteins, and metabolites are shown at the bottom of the figure, including the metabolite ATP and the protein DnaA, as well as the gene and mRNA dnaA that encode DnaA.

For each cell *i*, we denote the count matrices for DNA, mRNA, protein, and metabolite species as 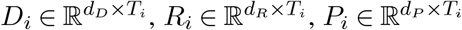, and 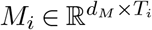, respectively, where *T*_*i*_ is the number of recorded time points in the cell cycle of cell *i*, and *d*_*D*_, *d*_*R*_, *d*_*P*_, and *d*_*M*_ denote the number of molecular species in each omic layer. Let 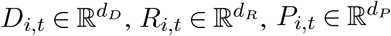, and *M*_*i*,_ 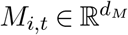represent the observed molecular counts of cell *i* at time *t*. This dataset offers a high-dimensional, multi-omics time-series structure with strong temporal dependence and cross-omic coupling, providing a realistic setting for evaluating the proposed COTree framework.

## 3. Methods

### 3.1. Overview of COTree

This paper presents a new statistical framework for analyzing cell-resolved multi-omics trajectory data, termed the Chrono-Omics Tree (COTree). As illustrated in Figure 2, COTree consists of two major stages:

**FIG 2.**
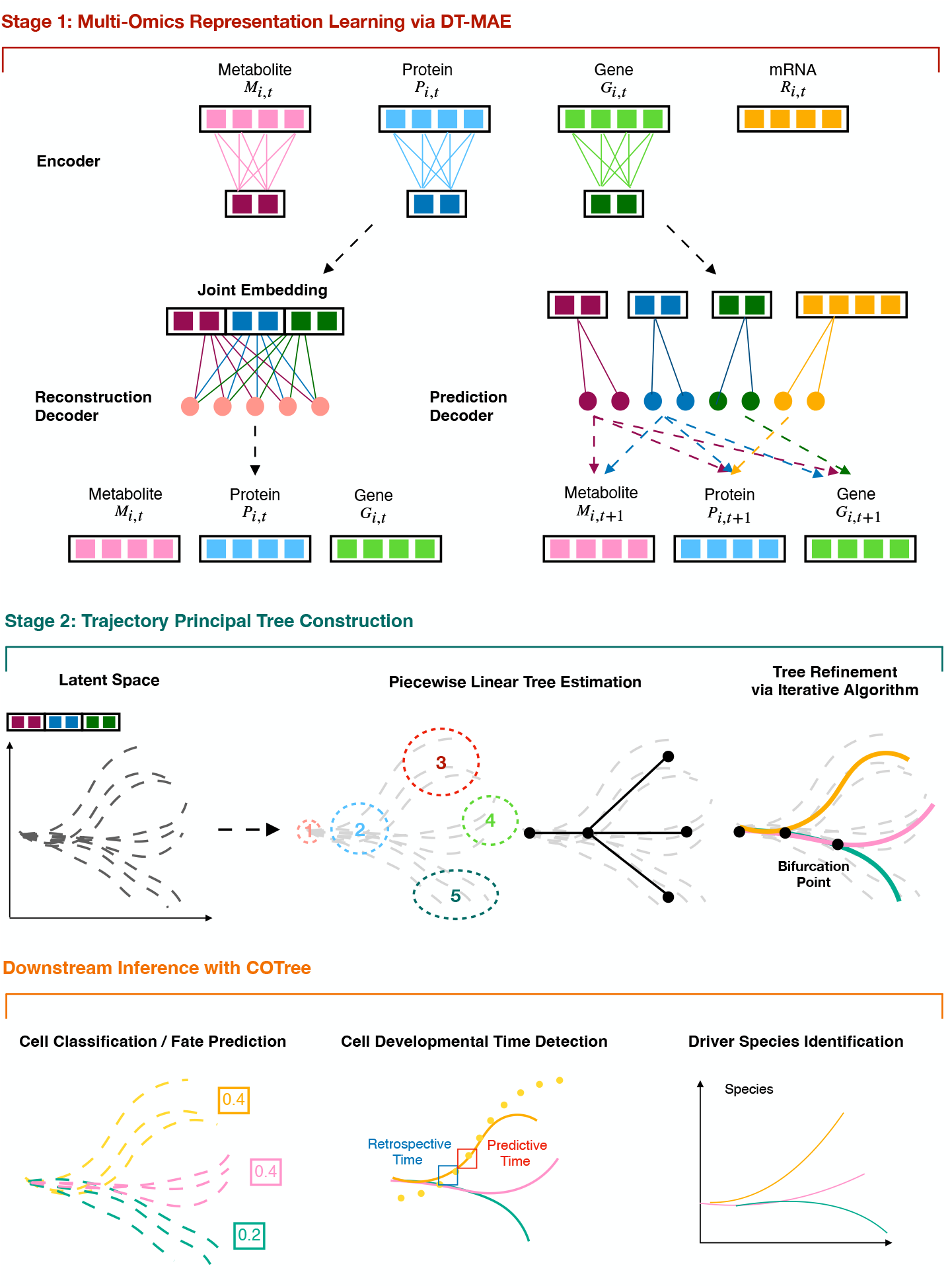
Overview of the COTree framework. COTree consists of two stages. Stage 1: A dual-target multi-omics autoencoder learns low-dimensional latent representations that integrate information across multiple omics layers. Stage 2: A trajectory principal tree is constructed in the latent space to capture cellular progression patterns. The resulting multi-omics representation and trajectory principal tree enable diverse downstream analyses, including cell classification, fate prediction, developmental time detection, and driver species identification.

- **Stage 1 – Multi-Omics Representation Learning**. In the first stage, we introduce a dualtarget multi-omics autoencoder (DT-MAE) to learn low-dimensional representations that integrate information across all omics layers. Unlike a classical autoencoder, DT-MAE employs a hybrid reconstruction–autoregressive decoder, enabling it to capture both continuous omics growth trends and transient bursts in cellular processes.
- **Stage 2 – Trajectory Principal Tree Construction**. In the second stage, we construct a trajectory principal tree that summarizes the overall cellular progression trend. The resulting tree delineates cellular lineages and identifies bifurcation nodes corresponding to key fate transitions.

The multi-omics representation and trajectory principal tree learned by COTree together enable a range of downstream inferences at both the cell and lineage levels, including cell classification, fate prediction, developmental time detection, and driver species identification.

### 3.2. Dual-Target Multi-Omics Autoencoder

The first stage of COTree aims to learn a compact representation that characterizes cellular states while preserving information relevant to cell growth. Although representation learning has been widely applied to biological data (Lopez et al., 2018; Brbić et al., 2020; Rosen et al., 2024), the high-temporal-resolution and multi-omics structure of trajectory data generated by whole-cell models presents new challenges for effective representation learning. To address these challenges, we introduce a dual-target multi-omics autoencoder (DT-MAE) specifically designed for cell-resolved multiomics trajectories.

The autoencoder is a powerful architecture for learning meaningful representations (Wang et al., 2019; Eraslan et al., 2019), as it captures complex nonlinear patterns in high-dimensional data through the reconstruction of the original signal (Hinton and Salakhutdi-nov, 2006; Ranzato et al., 2006; Bengio and LeCun, 2007). A classical autoencoder consists of two components: an encoder that compresses the input and a decoder that reconstructs the signal. However, this architecture is not directly applicable to multi-omics trajectories due to their inherent temporal and multimodal structure. Because different omics modalities exhibit distinct characteristics, we introduce a dedicated encoder for each omics layer. Specifically, we define encoders 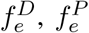, and 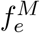 for DNA, protein, and metabolite measurements, respectively:

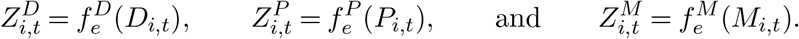

Here *D*_*i,t*_, *P*_*i,t*_, and *M*_*i,t*_ denote the observed counts of DNA, protein, and metabolites from cell *i* at time *t*, while 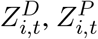, and 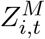represent their correpsonding latent embeddings. We do not learn representations for mRNA, as mRNA counts are highly unstable due to rapid synthesis and degradation in response to cellular demands for protein production.

To account for the temporal structure of the data, we introduce two sets of decoders: one reconstructs the current cell state, and the other predicts the future state. The reconstruction decoders ensure that the latent representations preserve information about the current cellular conditions, including transient events such as DNA replication and bursts of protein production (Ozbudak et al., 2002; Cai, Friedman and Xie, 2006). Specifically, the reconstruction module includes three decoders, 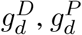, and 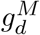, which reconstruct the counts of DNA, proteins, and metabolites at time *t*. The reconstructed values are expressed as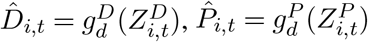, and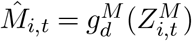. To encourage accurate reconstruction, we define the reconstruction loss as

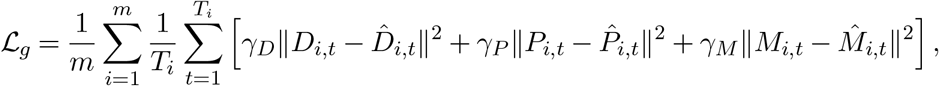

where *γ*_*D*_, *γ*_*P*_, and *γ*_*M*_ are tuning parameters that balance the contributions of different omics layers subject to the constraint *γ*_*D*_ + *γ*_*P*_ + *γ*_*M*_ = 1.

In contrast to the reconstruction decoders, the prediction decoders are designed to capture cell growth dynamics by leveraging both temporal and multimodal structures in the data. To this end, we introduce three decoders, 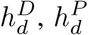, and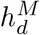, to predict the cell states at the next time point *t* + 1. During cellular growth, dependencies across different omics layers follow well-established biological principles (Berg, Tymoczko and Stryer, 2002; Alberts et al., 2014). DNA replication is regulated by initiation proteins, DNA polymerases, and accessory factors, while metabolites such as deoxynucleotides act as monomers for chromosome synthesis, and ATP provides the energy to unwind double-stranded DNA. Accordingly, the representations of DNA, protein, and metabolites are used to predict future DNA counts:

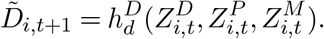

Proteins are synthesized from mRNA transcripts, and metabolites provide the energy necessary for translation and degradation processes. Thus, protein, metabolite, and mRNA features are used to predict future protein counts:

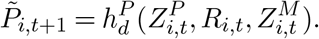

Finally, proteins act as enzymes that catalyze and regulate metabolic reactions, thereby influencing metabolite dynamics. Consequently, protein and metabolite representations are used to predict future metabolite counts:

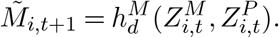

To encourage accurate prediction, we define the prediction loss as

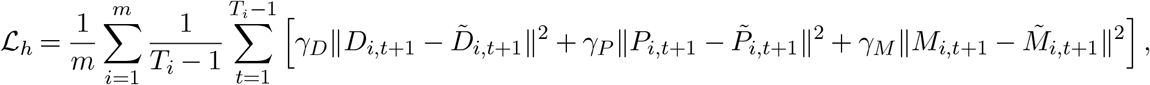

where *γ*_*D*_, *γ*_*P*_, and *γ*_*M*_ are the same tunning parameters used in ℒ_*g*_. Combining the reconstruction and prediction losses yields the dual-target objective:

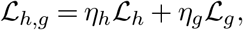

where *η*_*h*_ and *η*_*g*_ control the relative importance of the two objectives subject to the constraint *η*_*h*_ + *η*_*g*_ = 1. Through this dual-target formulation, the trained autoencoder captures information essential for both current and future cellular states. The resulting multi-omics representation is constructed by concatenating the embeddings of DNA, protein, and metabolites, i.e.,

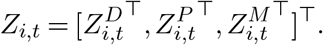

The hyperparameters of DT-MAE are selected using a two-stage procedure. In the first stage, we use Optuna (Akiba et al., 2019) to perform multi-criteria validation. We divide the cells into training and validation sets, retaining all observations from the same cell in the same set. In each trial, Optuna samples a candidate combination of hyperparameters, trains DT-MAE on the training cells, and evaluates three criteria on the validation cells: (i) prediction loss, defined as the unweighted sum of the modality-specific prediction errors; (ii) reconstruction loss, defined as the unweighted sum of the modality-specific reconstruction errors; and (iii) a manifold-smoothness criterion for the latent representation, defined as

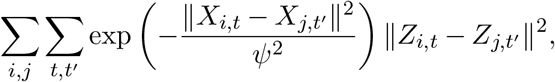

where *ψ* is the median of the pairwise distances between cell states in the preprocessed input space. The prediction and reconstruction criteria measure the predictive and reconstructive fidelity of the fitted model. We use unweighted validation losses so that their values cannot be reduced artificially by assigning small values to the modality weights *γ*_*D*_, *γ*_*P*_, and *γ*_*M*_. The manifold-smoothness criterion favors latent representations in which cell states that are similar in the input space remain locally similar, thereby informing the selection of *η*_*h*_ and *η*_*g*_.

After completing all trials, we retain the nondominated candidate models: those for which no other evaluated model performs at least as well on all three validation criteria and strictly better on at least one. In the second stage, we select from this set the model with the smallest sum of the validation prediction and reconstruction losses. The manifold-smoothness criterion is not included in this final selection criterion because it alone cannot exclude latent collapse: it equals zero when all latent embeddings are identical. Prediction and reconstruction performance are therefore used to select the final model from among the candidates that provide favorable tradeoffs among the three validation criteria. Further details on the tuning procedure and sensitivity analysis of the loss weights are provided in the Supplementary Material.

We further propose interpretability methods to examine the learned multi-omics representations from two complementary perspectives, species-level abundance and developmental progression, described in detail in the Supplemental Material.

### 3.3. Trajectory Principal Tree Construction

The second stage of COTree constructs a principal tree to summarize cellular progression trends in the latent representation space. Tree structures are commonly used in trajectory inference from snapshot measurements to characterize the progression order of cells (Street et al., 2018; Hou et al., 2023). In contrast to these approaches, our objective is to construct a cell growth tree from a collection of trajectories, 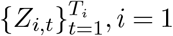, where the tree structure serves as a compact summary of all cellular trajectories. To address the alignment challenges inherent in analyzing multiple trajectories, where cells may progress at different rates of growth, we propose a trajectory principal tree construction algorithm comprising two steps. The first step estimates a piecewise linear tree, and the second step refines the structure through an iterative updating procedure. The resulting trajectory principal tree exhibits clear bifurcation points, with each branch represented as a smooth curve in the latent representation space.

The main presentation in this section focuses on trajectory principal-tree construction because a branching tree provides an appropriate summary of the JCVI-syn3A simulation trajectories analyzed in this paper. More generally, cellular state transitions need not follow a tree structure and may involve convergence, reversibility, or cycles (Tanay and Regev, 2017). We therefore extend COTree to the construction of non-tree trajectory graphs, as described in the Supplementary Material and evaluated in the controlled synthetic-data study.

#### 3.3.1. Step 1: Piecewise Linear Tree Estimation

To construct a piecewise linear tree, we first identify tree nodes by applying *k*-means clustering to snapshots of the trajectories. To better capture the terminal states of cell growth, the snapshots are divided into two sets: intermediate states (*Z*_*i,t*_, *t* ∈ [1, *T*_*i*_ − 1]) and terminal states 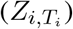. We perform *k*-means clustering separately on the two sets and then combine the resulting centroids to define the tree nodes. Next, we construct an adjacency matrix *A*, where each entry *A*_*i,j*_ records the number of trajectories transitioning from cluster *i* to cluster *j*. A minimum spanning tree is then derived from this adjacency matrix by defining the distance between nodes as 1*/A*_*i,j*_. Connecting the nodes with straight lines yields the piecewise linear tree that summarizes the overall cellular progression pattern.

To reduce the sensitivity of piecewise-linear tree construction to the user-specified number of initial clusters, we adopt an over-split-then-trim strategy. We first construct an intentionally overcomplete tree using relatively large numbers of intermediate and terminal states, following the procedure described above. This initialization may produce more candidate lineages than are needed to represent the underlying trajectory structure. Starting from this overcomplete tree, we greedily remove redundant lineages one at a time. Let *T*^(*k*)^ denote the tree at pruning iteration *k*, and let ℐ(*T* ^(*k*)^) denote its set of lineages. We define the overall projection error as

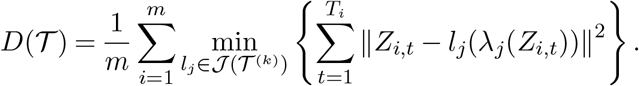

where *l*_*j*_(*λ*_*j*_(*Z*_*i,t*_)) is the projection of *Z*_*i,t*_ onto lineage *l*_*j*_ (see Step 2A for formal definitions). At each iteration, we identify the lineage whose removal produces the smallest increase in projection error:

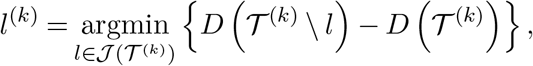

If removing *l*^(*k*)^ increases the projection error by no more than 10%, we update

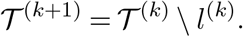

Otherwise, pruning stops. After selecting the final tree, we refit the nodes along each retained lineage using nearby cell states. Further implementation details are provided in the Supplementary Material. Figure S2 examines sensitivity to the initial numbers of intermediate and terminal states and shows that the resulting tree is stable once the initial tree is sufficiently overcomplete.

#### 3.3.2. Step 2: Iterative Algorithm for Tree Refinement

Given the initial piecewise linear tree, the second step iteratively refines the structure to better capture the underlying trajectory patterns. In each iteration, lineages are first updated independently using a time-informed principal curve algorithm (Step 2A), followed by bifurcation point detection and merging of shared branches (Step 2B). The algorithm minimizes the following loss function:

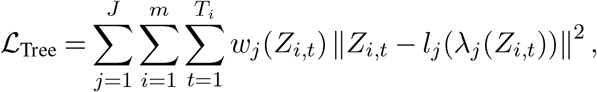

where *w*_*j*_(*Z*_*i,t*_) denotes the weight representing the contribution of point *Z*_*i,t*_ to lineage *l*_*j*_. The loss _Tree_ quantifies how well the trajectory principal tree captures the global trend of cellular progression. The iterative updates continue until the relative change in loss between successive iterations falls below a specified threshold *ϵ*. The final trajectory principal tree is denoted by *T* = (*V*, ℰ), where V represents the set of nodes and ℰ the set of branches.

#### 3.3.3. Step 2A: Lineage Update via Time-Informed Principal Curve

The principal curve algorithm is a widely used nonlinear method for learning smooth curves that summarize high-dimensional data points (Hastie and Stuetzle, 1989). It has been extensively applied to trajectory inference in single-cell RNA-sequencing analyses (Qiu et al., 2017a; Street et al., 2018). However, classical principal curve methods are designed for static data points rather than trajectories and therefore ignore temporal information. To incorporate temporal structure, we propose a time-informed principal curve algorithm that summarizes trajectories while preserving their temporal order and is used to update each lineage *l*_*j*_. Specifically, the lineage *l*_*j*_ is updated through the following steps:

1. **Estimate the growth stage**. Each point *Z*_*i,t*_ is projected onto the lineage *l*_*j*_, and the arc length *λ*_*j*_(*Z*_*i,t*_) between the root and its projection is computed. The arc length *λ*_*j*_(*Z*_*i,t*_) quantifies how far cell *i* has progressed from the initial state, referred to as the growth stage.
2. **Adjust the growth stage using temporal order**. We incorporate the observed temporal ordering directly by estimating growth stages under a monotonicity constraint. For each cell, isotonic regression projects the unconstrained growth-stage estimates onto the set of sequences that are nondecreasing across observation times (Barlow et al., 1972). The resulting estimates preserve temporal order while minimizing the squared deviation from the unconstrained estimates.
3. **Evaluate regression weights**. The weight *w*_*j*_(*Z*_*i,t*_) determines the contribution of cell state *Z*_*i,t*_ to the update of lineage *l*_*j*_. It combines point-level proximity with trajectory-level assignment:

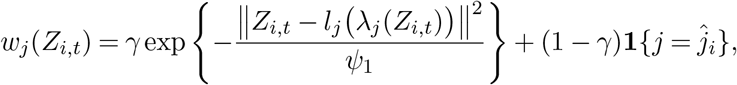

where 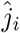 is the closest lineage of cell *i*

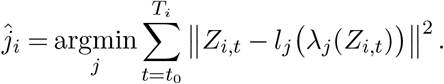

Here, *γ* ∈ [0, 1] controls the relative contributions of the two components, and *ψ*_1_ is set to the median of the point-level squared distances across all point-lineage pairs. The first component assigns greater weight to cell states located near lineage *l*_*j*_, whereas the second assigns the complete trajectory of cell *i* to its closest lineage. Thus, *γ* balances local geometric proximity against trajectory-level coherence.
4. **Update the lineage**. Each coordinate of *Z*_*i,t*_ is regressed on *λ*_*j*_(*Z*_*i,t*_) using *w*_*j*_(*Z*_*i,t*_) as observation weights. We employ locally estimated scatterplot smoothing (LOESS) (Cleve-land, 1979) for this step, though other smoothing techniques may also be applied.

The key differences from the classical principal curve algorithm are twofold: (1) the projection and growth stage are dynamically adjusted using temporal information, and (2) the weights incorporate trajectory–lineage similarity. The growth stage provides a quantitative measure of each cell’s relative progress through the cell cycle, thereby aligning biologically equivalent states and mitigating variability due to differences in growth rates.

The mixing parameter *γ* used in the lineage-update weights is selected by grid search based on the separation of the resulting lineage assignments. For each candidate value, we fit the trajectory tree and calculate the projection distance from each cell trajectory to each estimated lineage. These distances are converted into soft lineage-assignment probabilities using a Gaussian kernel whose bandwidth parameter is set to the median of the squared trajectory-to-lineage projection distances. We then calculate the entropy of the assignment probabilities for each cell and average it across cells. The value of *γ* producing the lowest average assignment entropy is selected, favoring a tree construction in which cell trajectories can be assigned more clearly to individual lineages.

#### 3.3.4. Step 2B: Bifurcation Point Detection

After updating all lineages in Step 2A, we reconstruct the overall tree structure by detecting bifurcation points and merging similar branches. Suppose there are *J* distinct lineages 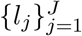 whose growth stages (i.e., arc lengths) lie within a finite interval Λ. To initialize the iterative bifurcation-detection procedure, we set *λ*_0_ = 0 and 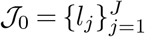. Given the *k*-th bifurcation point at *λ*_*k*_ and a lineage set ℐ_*k*_, the goal is to identify the earliest growth stage *λ*_*k*+1_ *> λ*_*k*_ at which at least one lineage diverges from the others. Formally, we select

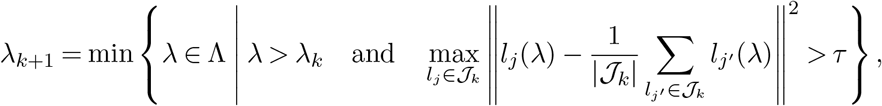

where *l*_*j*_(*λ*) denotes the point on lineage *l*_*j*_ at growth stage *λ*, and *τ >* 0 is a predefined divergence threshold. The lineage that diverges at this stage is then identified as

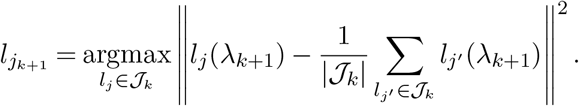

Because lineage 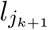 separates at *λ*_*k*+1_, all lineages in ℐ_*k*_ are merged between *λ*_*k*_ and *λ*_*k*+1_ according to

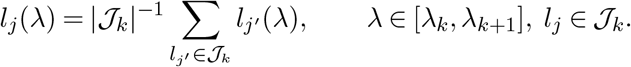

The bifurcation point is then defined as *l*_*j*_(*λ*_*k*+1_). After detecting the bifurcation and merging branches, we update the lineage set by 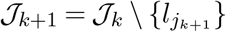. This iterative procedure continues until only one lineage remains, producing a complete trajectory principal tree with clearly defined branching topology.

We use the scale-adjusted divergence threshold *τ* = *v*^2^*R*^2^, where *R* is the maximum pair-wise distance between the unmerged lineage curves and *v* specifies the relative separation required to identify a bifurcation. We recommend *v* = 0.05 in practice. Thus, a time point is identified as a bifurcation point when the distance between a lineage and the corresponding mean trajectory exceeds 5% of the maximum pairwise lineage distance.

### 3.4. Downstream Inference with COTree

The multi-omics representations and trajectory principal tree learned by COTree provide a concise yet comprehensive summary of cell-resolved multi-omics trajectory data. A natural question that follows is how these summaries can deepen our understanding of cellular dynamics. In this section, we introduce a series of downstream analyses enabled by the learned representations and tree structure, including cell classification, fate prediction, developmental time detection, and driver species identification.

#### 3.4.1. Cell Classification

The trajectory principal tree serves not only as a summary of cellular developmental progression but also as an efficient tool for classifying cell trajectories. Because the nodes of the trajectory principal tree correspond to key milestones of cell development, a natural approach to classify cell trajectories is based on their proximity to these milestones. Given a cell trajectory *X*_1_, …, *X*_*T*_, we first apply the encoder to map it into the latent representation space, obtaining *Z*_1_, …, *Z*_*T*_ = *f*_*e*_(*X*_1_), …, *f*_*e*_(*X*_*T*_). We then project this latent trajectory onto the trajectory principal tree *T*. Let *P*_*v*_ denote the path in *T* connecting the root to a node *v* ∈ *V*. The nearest milestone is identified by

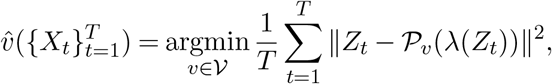

where *P*_*v*_(*λ*(*Z*_*t*_)) represents the projection of *Z*_*t*_ onto the path *P*_*v*_. The cell trajectory is then classified according to its nearest milestone 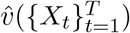, which reflects its developmental position along the trajectory principal tree.

#### 3.4.2. Cell Fate Prediction

Beyond cell classification, the nearest milestone on the trajectory principal tree can also be used to predict cell fate. Intuitively, a cell’s future state is expected to correspond to one of the descendant leaf nodes of its nearest milestone, if such descendants exist. When multiple descendant leaf nodes are present, it is natural to estimate the probability of developing into each possible future state. Suppose we observe *n* cell trajectories, denoted by 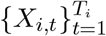. Estimating the probability of reaching a given leaf node is nontrivial because some trajectories may be truncated before reaching the terminal state. To estimate these probabilities, we first identify the nearest milestone for each trajectory, 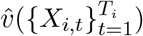, for *i* = 1, …, *n*. Given these assignments, the probability of reaching the subtree below a node *v* is estimated as

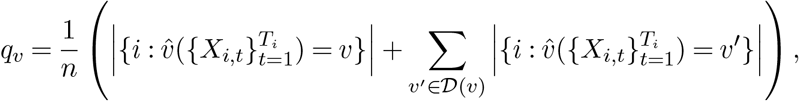

where *D*(*v*) denotes the set of descendant nodes of *v*. To obtain the probability of reaching the leaf nodes, we use a top-down iterative algorithm. Starting from the root node *ρ*, we set *p*_*ρ*_ = 1. Given a node *v* with probability *p*_*v*_, the probability for each of its children *v*^*′*^∈*C*(*v*) is updated as

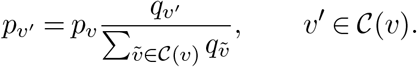

Now, suppose we observe a new cell trajectory *X*_1_, …, *X*_*T*_ and wish to predict its future state. We first determine its nearest milestone 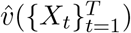 and identify all descendant leaf nodes 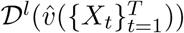. The conditional probability of the trajectory reaching a particular leaf node 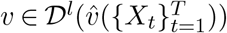 is then given by

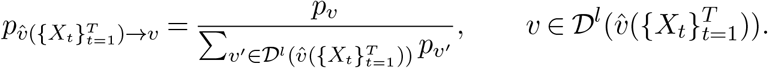

Accordingly, the trajectory *X*_1_, …, *X*_*T*_ is predicted to reach node *v* with probability 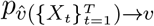 for all 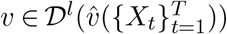. This probabilistic formulation enables quantitative prediction of cell fates, providing a principled framework for studying lineage commitment and uncertainty in cellular development.

#### 3.4.3. Cell Developmental Time Detection

To better understand cellular dynamics, it is important to identify when cells reach key developmental milestones and when their fates become determined. We focus on two types of developmental time: (1) the retrospective time 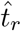, inferred by tracing the entire trajectory, and (2) the earliest predictive time 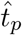, estimated when only a partial trajectory is observed. Given a lineage 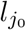, we define the retrospective time 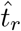 as the earliest time point at which the projection of a cell trajectory onto the principal tree crosses the final bifurcation node *v* of 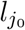. Specifically, for an observed cell trajectory *X*_1_, …, *X*_*T*_, we transform it into the latent representation trajectory *Z*_1_, …, *Z*_*T*_ and identify the closest lineage 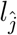such that

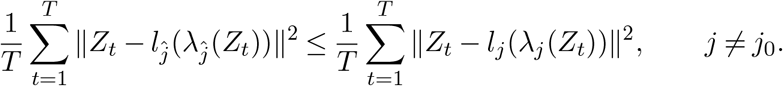

If the closest lineage 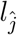differs from 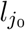, we set 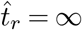 and 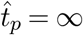. If 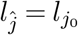, let *λ*_*v*_ denote the arc length from the root to node *v*, and define

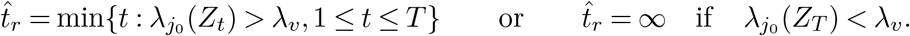

A value of 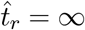 indicates that the cell trajectory has not, or will not, reach the terminal stage of lineage 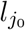. The predictive time 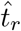 characterizes the earliest time at which the cell fate can be reliably predicted. We define it as the latest time point at which the cell fate remains ambiguous:

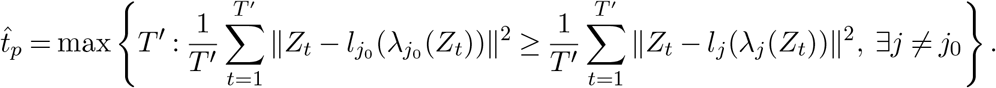

Together, 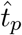 and 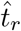 provide complementary insights: 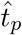marks when a cell’s fate is realized, while 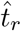 marks when it becomes predictable based on its partial trajectory. These quantities offer a quantitative framework for studying the timing of cell fate determination and heterogeneity in developmental progression.

#### 3.4.4. Driver Species Identification

Cell trajectories may exhibit abrupt directional changes within a lineage or diverge into distinct developmental paths at bifurcation points. A key question is which molecular species contribute most to these directional shifts. A major challenge in identifying such driver species lies in the fact that directional changes are characterized in the latent representation space, which is not directly linked to species-level abundances. To bridge this gap, we propose a method to infer the local direction of change in the species space that corresponds to the trajectory’s movement in the representation space.

Let 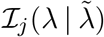 denote the collection of trajectory segments 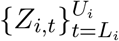 whose nearest lineage is *l*_*j*_ and whose growth stage (arc length) lies between *λ* and 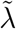. We use data points in 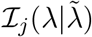 to infer the direction of change near stage *λ* along lineage *l*_*j*_. Given 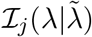, we define the average correlation measuring how well a directional perturbation in species space aligns with the direction of change in representation space as

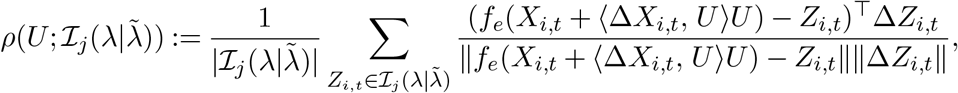

where Δ*X*_*i,t*_ = *X*_*i,t*+Δ*t*_ − *X*_*i,t*_, Δ*Z*_*i,t*_ = *Z*_*i,t*+Δ*t*_ − *Z*_*i,t*_, and 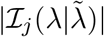 is the number of measurements in 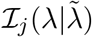. Here, Δ*t >* 0 is a small time interval. The direction *U* in species space that maximizes 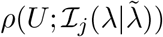 is interpreted as the direction most consistent with the trajectory’s change in the latent representation space.

With a means to infer directions in species space, we can compare directional changes across trajectories to identify driver species. Given two sets of trajectory segments, 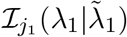 and 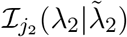, we estimate the corresponding directions by solving

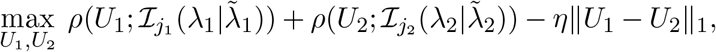

where the *L*_1_ penalty promotes sparsity in the difference between *U*_1_ and *U*_2_. The set of driver species is defined as those with large absolute differences between *U*_1_ and *U*_2_:

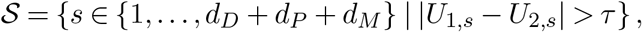

where *τ* is an automatically selected threshold (see Supplementary Materials for details). Depending on the type of directional change, different collections of trajectory segments can be compared. To identify driver species associated with a bifurcation node *v*, we select 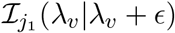 and 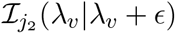, where 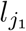 and 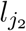are the lineages diverging at *v* and *λ*_*v*_ is the growth stage of bifurcation point *v*. To study a local directional shift along a lineage *l*_*j*_ at growth stage *λ*, we instead use ℐ_*j*_(*λ λ* + *ϵ*) and ℐ_*j*_(*λ λ ϵ*). This framework enables the systematic identification of molecular species driving local and lineage-specific transitions in cellular dynamics.

## 4. Controlled Synthetic-Data Study

To evaluate whether COTree can accurately and robustly infer cellular trajectories and recover their underlying topology, we generate synthetic two-omics datasets with known lineage structures. We construct each ground-truth trajectory in a low-dimensional latent space, with individual branches following curved, non-linear paths. Simulated cells progress along predefined lineages, and perturbations are added to their latent states to represent cell-to-cell biological variability. These latent states are subsequently mapped to the observed feature spaces through modality-specific Poisson emission models. Additional details of the simulation design are provided in the Supplementary Materials. Because the lineage assignments, bifurcation times, and trajectory topologies are known by construction, the inferred trajectories can be evaluated directly against the ground truth.

We evaluate COTree from two complementary perspectives: tree-topology recovery and cell-lineage inference. For topology recovery, we determine whether COTree identifies the correct number of lineages, measured by lineage-number recovery accuracy, and reconstructs the correct sequence of bifurcation events, measured by bifurcation-order accuracy. For cell-lineage inference, we use the adjusted Rand index (ARI) to quantify agreement between the inferred and ground-truth lineage assignments.

We first isolate and evaluate the tree-construction algorithm by applying it directly to the ground-truth latent representations. As shown in the second column of Figure 3(a), COTree accurately reconstructs the tree topology and the order of the branching events, demonstrating the effectiveness of its tree-construction procedure. We next evaluate the complete COTree pipeline. In this setting, DT-MAE first learns low-dimensional representations from the observed count data, and COTree then uses these representations to construct the trajectory tree.

**FIG 3.**
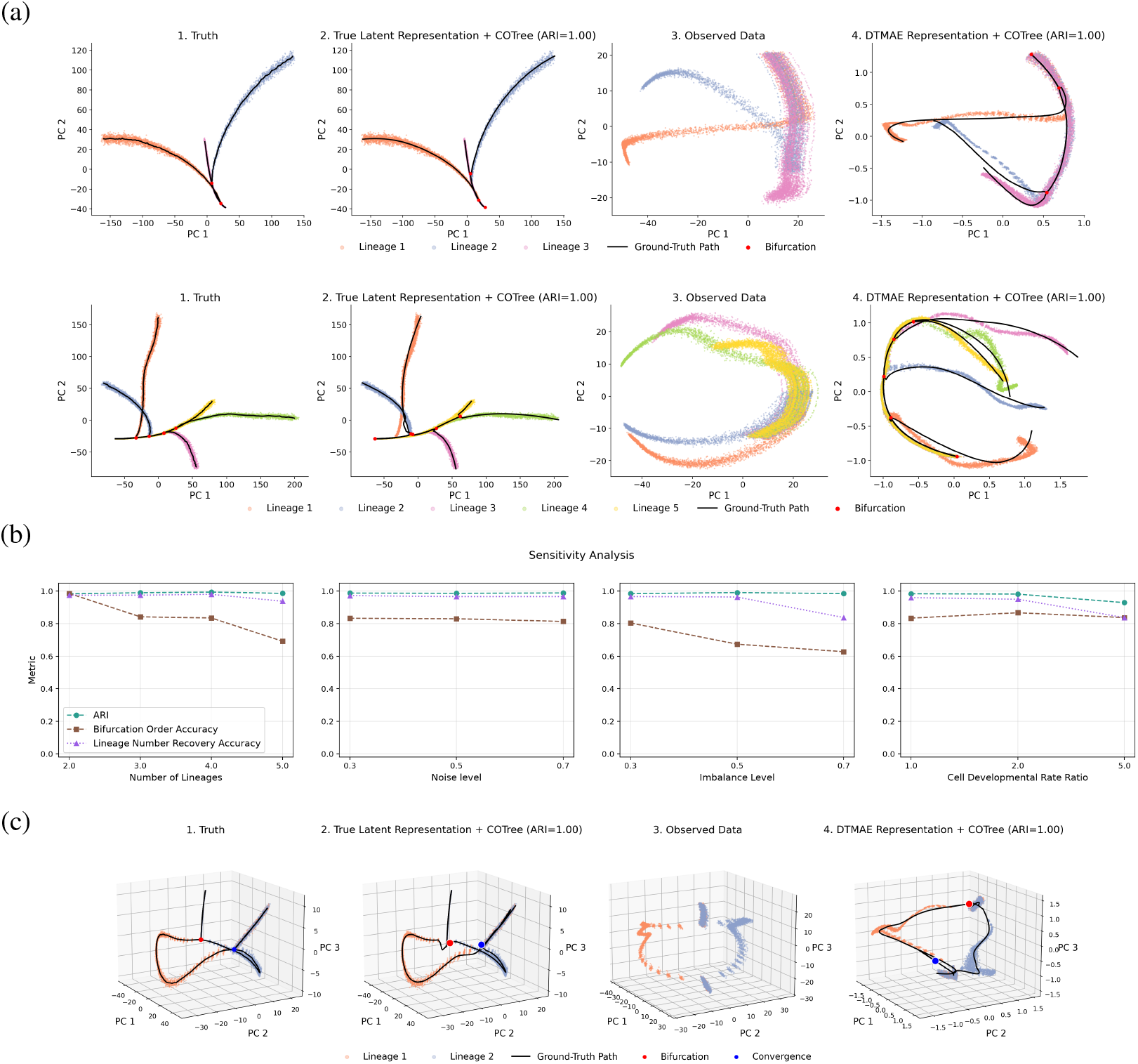
COTree robustly recovers tree topology and cell lineages. (a) compares trees constructed from the true latent representation and DT-MAE representations with the ground-truth tree in simulation. (b) presents sensitivity analysis under varying numbers of lineages, different noise levels, imbalanced cell numbers across lineages, and varying ratios of developmental rates between lineages. Cell classification performance is evaluated using the average adjusted Rand index between inferred cell lineages and the ground truth across 300 iterations, while tree topology recovery is evaluated by the rates of recovering the correct number of lineages and the correct ordering of bifurcations across simulation replicates. (c) presents the topology-reconstruction results obtained by applying COTree to data with a non-tree structure.

Although sampling noise obscures the underlying structure in the observed feature spaces, the complete pipeline recovers the correct topology and accurately assigns cells to their respective lineages, as shown in Figure 3(a). Together, these experiments demonstrate the effectiveness of both the representation-learning and tree-construction components of COTree. We further examine the robustness of COTree under a range of simulation conditions, including different numbers of lineages, varying noise levels, imbalanced lineage sizes, and lineage-specific developmental rates. Performance is evaluated using the same three criteria: ARI, bifurcation-order accuracy, and lineage-number recovery accuracy. The results, summarized in Figure 3(b), show that COTree consistently recovers the underlying cellular dynamics across diverse simulation settings. Finally, we investigate whether COTree can accommodate trajectory structures that are more complex than a conventional branching tree. As illustrated in Figure 3(c), COTree successfully reconstructs the simulated structure, indicating its potential to recover more general, non-tree trajectory topologies.

## 5. Whole-Cell Model Application

We apply COTree to the dataset described in Section 2. The cells in this dataset can be broadly categorized into three types: healthy cells with multiple replication events, healthy cells with a single replication event, and unhealthy cells that experience phosphoenolpyruvate (PEP) depletion and die. Given its well-characterized biological background and clearly defined cell types, this dataset serves as an effective benchmark for evaluating the performance of COTree in multi-omics trajectory analysis. Implementation details of COTree applied to this dataset are provided in the Supplemental Materials.

### 5.1. Multi-Omics Representation Preserves Biological Signals

This section evaluates how well the multi-omics representations learned by the dual-target multi-omics autoen-coder (DT-MAE) preserve biologically meaningful variation in cell-resolved trajectories. We compare DT-MAE with several commonly used non-temporal dimension-reduction and representation-learning methods, including principal component analysis (PCA), t-SNE (Maaten and Hinton, 2008), UMAP (McInnes, Healy and Melville, 2018), and MOFA+ (Argelaguet et al., 2020). Because these methods have objectives different from that of DT-MAE, we use them as reference methods rather than as direct trajectory-aware competitors. The purpose of this comparison is to illustrate how the learned representation differs when temporal information is explicitly incorporated. In DT-MAE, the latent representations for genes, metabolites, and proteins have dimensions 10, 50, and 50, respectively. As shown in Figure 4(a), DT-MAE separates trajectories associated with different cell types, whereas these groups are less clearly distinguished by the non-temporal reference methods. The DT-MAE representation also exhibits a clearer organization of cellular progression over time, consistent with the intended role of its autoregressive prediction objective. To quantify the biological information retained in each representation, we apply linear probing, in which a linear classifier is trained on the representations at each time point (Alain and Bengio, 2016). As summarized in Figures S3(a) and S3(b), cell types become distinguishable at earlier developmental stages using the DT-MAE representation than using the reference representations. Within the present dataset, these results indicate that DT-MAE preserves biological variation associated with cellular progression and heterogeneous cell outcomes.

**FIG 4.**
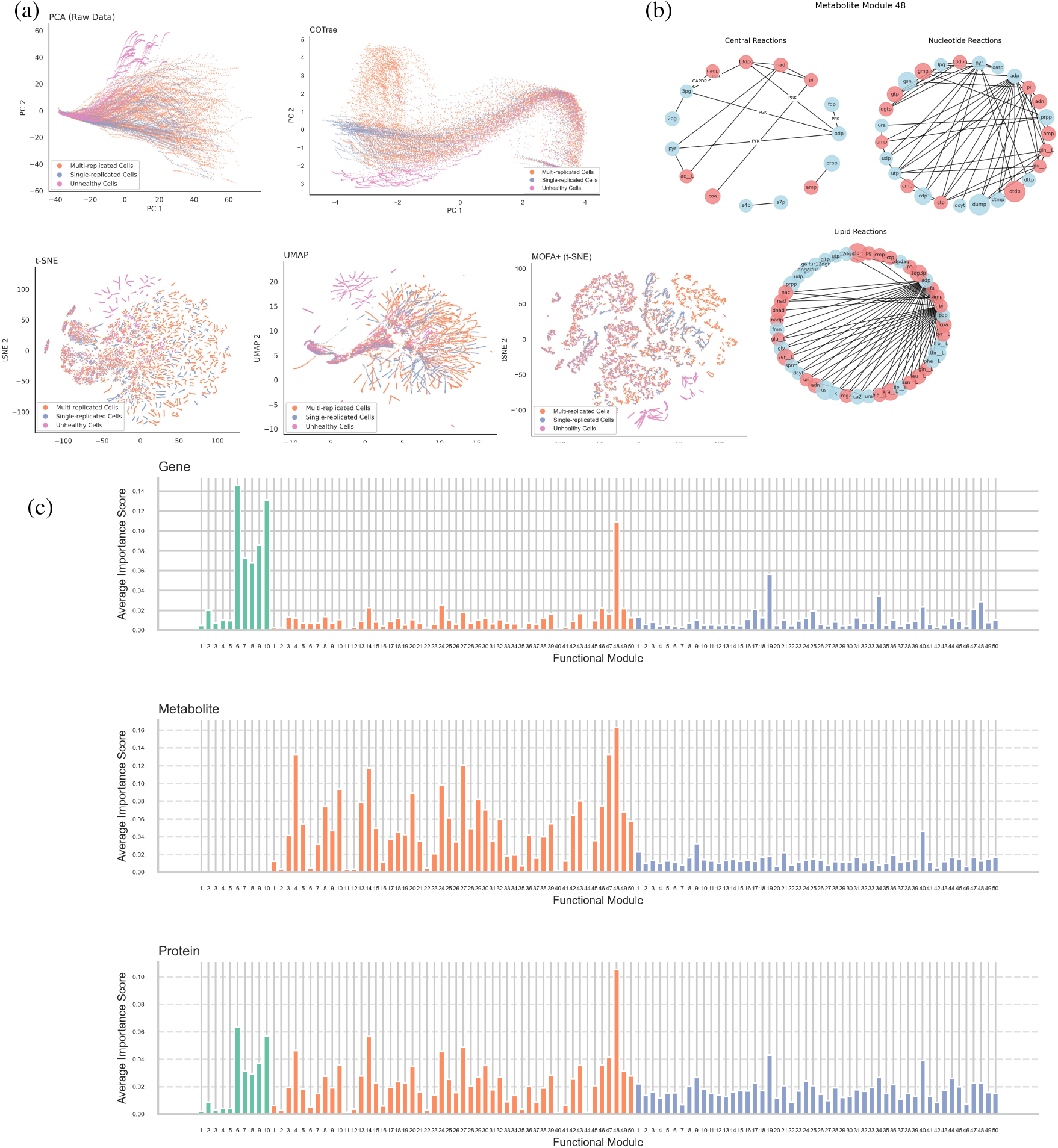
COTree effectively preserves biological signals through representation learning. (a) presents latent representations of multi-omics trajectories learned by DT-MAE, PCA, t-SNE, UMAP, and MOFA+. Each point represents a cell, colored by cell type. (b) shows metabolite functional module 48, where colors indicate species with positive (blue) or negative (red) contributions within the module, and marker size reflects the magnitude of their contribution. Species participating in the same reactions are connected by edges. (c) summarizes average importance scores of functional modules in predicting gene, metabolite, and protein abundances. Green, orange, and blue denote gene, metabolite, and protein functional modules, respectively.

Beyond separating cell types, the representations learned by DT-MAE also capture the coordinated behavior of groups of molecular species involved in related biological processes. As detailed in Section S1 of the Supplemental Material, each coordinate of the multi-omics representation can be interpreted as corresponding to a functional module, a group of molecular features exhibiting change patterns similar to that coordinate. In this dataset, DT-MAE uncovers 10 gene, 50 metabolite, and 50 protein functional modules, each reflecting shared progression patterns among species. In the gene functional modules, most genes in modules 8 and 9 finish replication after 40 minutes, whereas most genes in modules 6 and 10 finish replication before 40 minutes (Figure S4). The metabolite functional modules typically consist of species associated with one or two key metabolites that play central roles in cell growth. For example, metabolite module 48 contains metabolites participating in the central glycolytic reactions GAPDP, PFK, and PGK, together with metabolites involved in ADP-associated reactions, linking this module to glycolysis and cellular energy metabolism (Figure 4(b)).

The protein modules also exhibit functional organization (Table S2). Protein module 40 is enriched in proteins associated with genome maintenance and nucleotide metabolism, including DNA replication and repair, and also contains proteins involved in ribosome- and tRNA-related processes, redox regulation, and cellular growth. Protein module 19 contains enzymes associated with ATP production. The grouping of functionally related species into coherent modules demonstrates that the representations learned by DT-MAE effectively preserve the underlying biological structure. To assess the relative influence of these functional modules on cellular progression, we evaluate their average importance scores in predicting gene, metabolite, and protein abundances (Figure 4(c)). Notably, metabolite functional module 48 exhibits the highest average importance score, consistent with the important role of energy synthesis in cell growth. Collectively, these results show that the multi-omics representations learned by DT-MAE capture biologically meaningful and interpretable patterns across molecular layers.

### 5.2. Trajectory Principal Tree Captures Cellular Progression Trend and Reveals Cell Types

The second stage of COTree constructs a trajectory principal tree atop the learned multi-omics representations, providing a global summary of cellular growth dynamics. The trajectory principal tree identifies three distinct lineages (Figure 5(b)), corresponding to three major cell types in the dataset. When cell trajectories are classified according to this tree, the resulting lineages exhibit clear biological distinctions: all unhealthy cells are assigned to lineage 1; most cells undergoing a multiple replication event belong to lineage 2; and cells in lineage 3 predominantly display single replication events (Figure 5(d)). This observation further validates the ability of the trajectory principal tree to uncover biologically meaningful structure within the data. Beyond cell-type identification, the trajectory principal tree also characterizes the timing of key developmental transitions, such as when trajectories diverge at bifurcation nodes. As shown in Figure 5(e), unhealthy cells typically cross the bifurcation node and enter lineage 1 at approximately 15 minutes, indicating that early signs of cell death emerge soon after the onset of the cell cycle.

**FIG 5.**
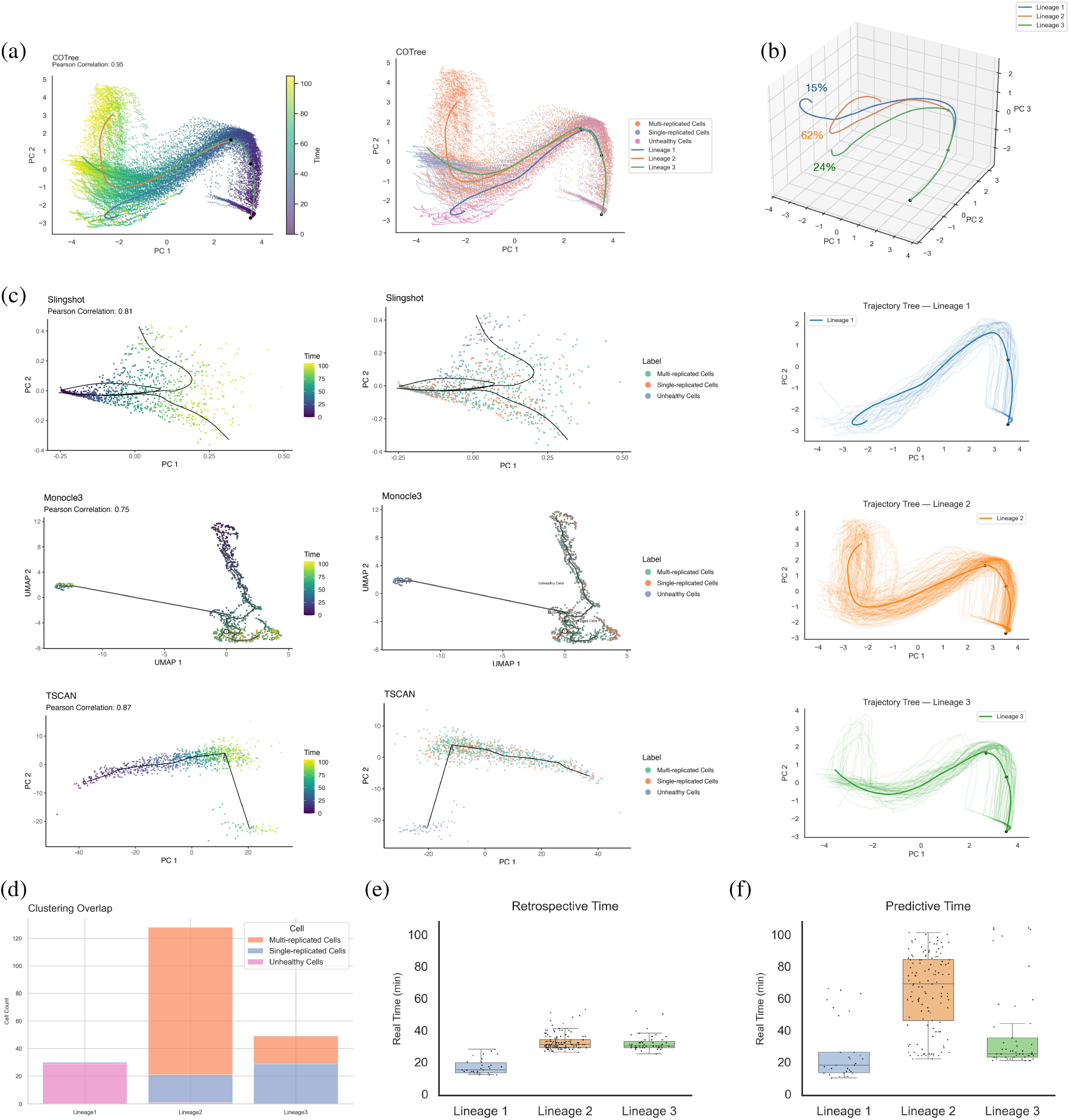
COTree reveals three major cell types in the dataset. (a) shows trajectory principal tree constructed by COTree. Each point represents a cell measurement at a given time, colored by time and known biological labels. COTree identifies three distinct cellular lineages. (b) presents estimated probabilities of cell development toward each node and trajectories of cells classified into the corresponding lineages. (c) shows trajectory trees constructed by Slingshot, Monocle3, and TSCAN for comparison. (d) presents numbers of cells classified into each lineage; colors indicate multi-replicated, single-replicated, and unhealthy cells. (e) and (f) summarize boxplots of retrospective and predictive times in cell development time detection.

The trajectory principal tree also serves as a valuable tool for early prediction of cell fate. Given a new cell trajectory, it can be projected onto the principal tree to determine the cur-rent growth stage, and the predicted cell fate is represented by the probability of reaching each leaf node. Figure 5(f) summarizes the earliest time at which cell fate can be accurately predicted. For instance, the fate of unhealthy cells can be predicted at 26 minutes on average, whereas their actual mean death time, defined as the point when PEP concentration reaches zero, is approximately 57.5 minutes. These results demonstrate that the trajectory principal tree enables early and accurate fate prediction while capturing heterogeneity in cellular developmental outcomes.

Having demonstrated the performance of the trajectory principal tree, we next compare COTree with existing trajectory inference methods. Most existing tree construction approaches are designed for snapshot measurements, making direct comparisons somewhat unequal, as COTree is specifically developed for trajectory data. We compare COTree with three state-of-the-art methods: TSCAN (Ji and Ji, 2016), Monocle3 (Qiu et al., 2017b), and Slingshot (Street et al., 2018). Because these existing methods require snapshot data as input, we randomly sampled five time points from each trajectory to create a comparable dataset. As shown in Figures 5(a) and 5(c), the trees constructed by existing methods fail to capture the continuous progression of cellular development and tend to mix cells from different types. Furthermore, their performance remains similar even when all time points in the trajectories are used (Figure S5). In contrast, by leveraging the temporal information inherent in full trajectories, COTree’s trajectory principal tree accurately recovers both the global cell growth trend and the distinct lineage structure corresponding to different cell types.

In addition to constructing a trajectory principal tree from the integrated multi-omics representation, we can also build trees based on each individual omics layer. These single-omics trajectory principal trees reveal omics-specific progression dynamics and provide additional resolution for refining cell types. Similar to the multi-omics tree, the trees constructed from metabolite and protein representations both exhibit three lineages (Figures 6(a) and 6(b)). Cells classified into lineages defined by metabolite or protein representations show substantial overlap with those defined by the multi-omics representation (Figure 6(c)), while subtle differences highlight previously unrecognized subpopulations. For example, eleven cells classified into lineage 3 under the multi-omics representation are reassigned to lineage 2 under the metabolite-based representation. These cells, similar to most with active gene replication events, exhibit elevated phosphoenolpyruvate (PEP) levels and reduced dATP levels compared to other lineage 2 cells in the multi-omics tree (Figure 6(d)), suggesting sufficient energy availability and high consumption of deoxynucleotides. This newly identified cell sub-type implies that favorable metabolite conditions may promote multiple replication events but are not sufficient to initiate them without supportive signals from other omics layers.

**FIG 6.**
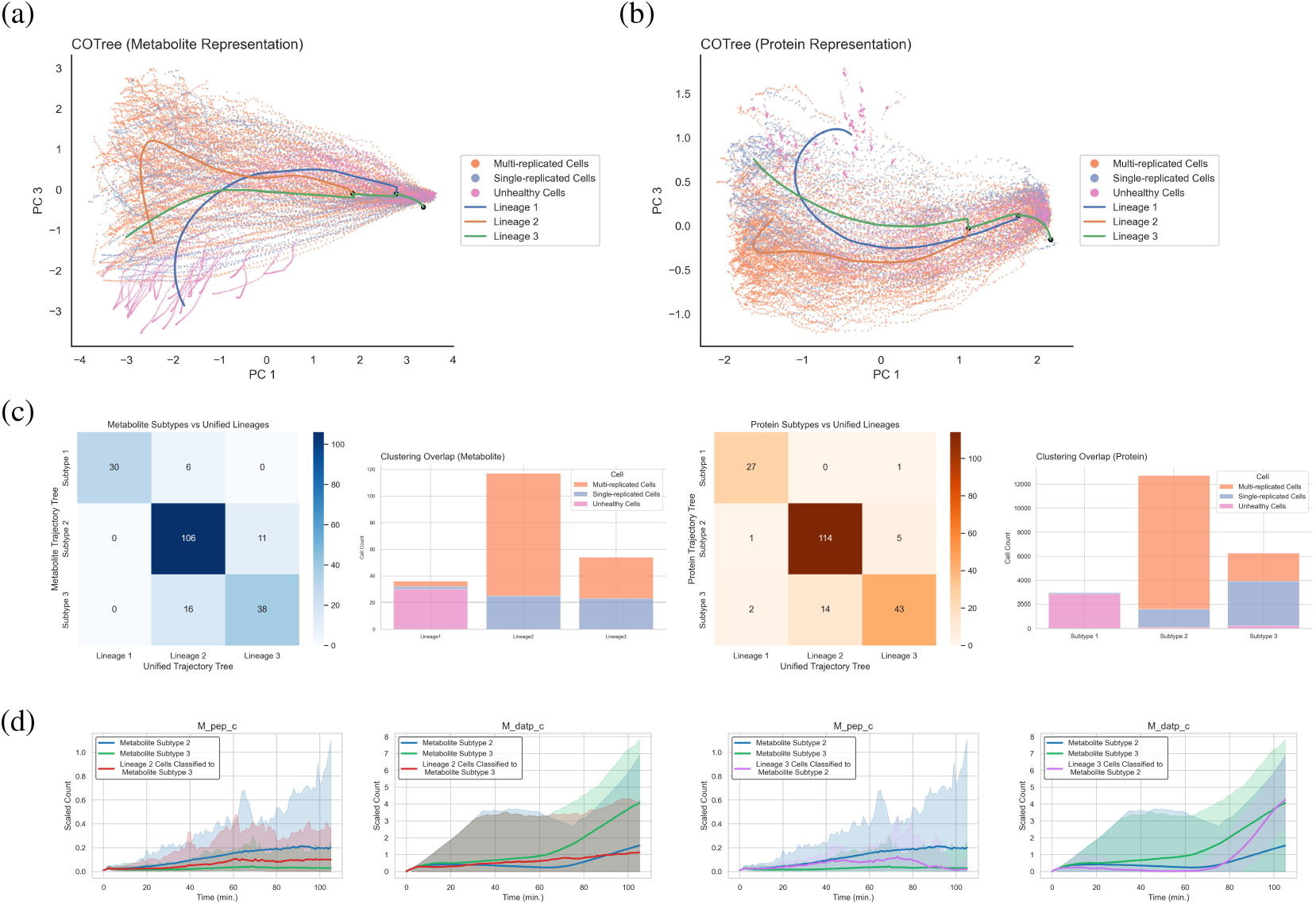
Trajectory principal trees built on single-omics representations reveal cell subtypes with modality-specific characteristics. (a) and (b) show trajectory trees constructed from metabolite and protein representations, respectively. (c) presents overlap between modality-specific subtypes and multi-omics lineages, together with the composition of each subtype in terms of multi-replicated, single-replicated, and unhealthy cells. (d) illustrates temporal trends of the metabolite species phosphoenolpyruvate (PEP) and deoxyadenosine triphosphate (dATP) across subtypes.

### 5.3. Downstream Inference Identifies Driver Species

Beyond identifying cell types and characterizing cellular dynamics, the multi-omics representation and trajectory principal tree also facilitate the identification of driver species. Because the trajectory principal tree contains two bifurcation nodes, we first examine which species contribute most to these divergence events (Figure 7(a)). At the first bifurcation node, the lineage of unhealthy cells separates from the other two, suggesting that the significant species are closely associated with early signs of cell death. Specifically, cells in lineage 1 exhibit reduced levels of key upstream metabolites, fructose 1,6-bisphosphate (FDP), glucose-6-phosphate (G6P) and sedoheptulose 7-phosphate (S7P), which are essential for converting extracellular glucose into energy (Figure 7(b)). As these metabolites are central intermediates in the glycolytic pathway (Figure 7(d)), the results indicate that impaired glucose import and upstream glycolytic processing may cause energy depletion in later stages of the cell cycle. Analysis of the second bifurcation node identifies key species associated with DNA replication, where the single-replication and multiple-replication lineages diverge. In particular, cells in lineage 2 show elevated levels of phosphate (Pi), pyrophosphate (PPi), and 1,3-bisphosphoglycerate (13DPG), along with an increased abundance of the replication initiator protein DnaA/0001. These findings suggest that elevated energy supply and activation of replication initiation protein may together promote additional replication events during the cell cycle.

**FIG 7.**
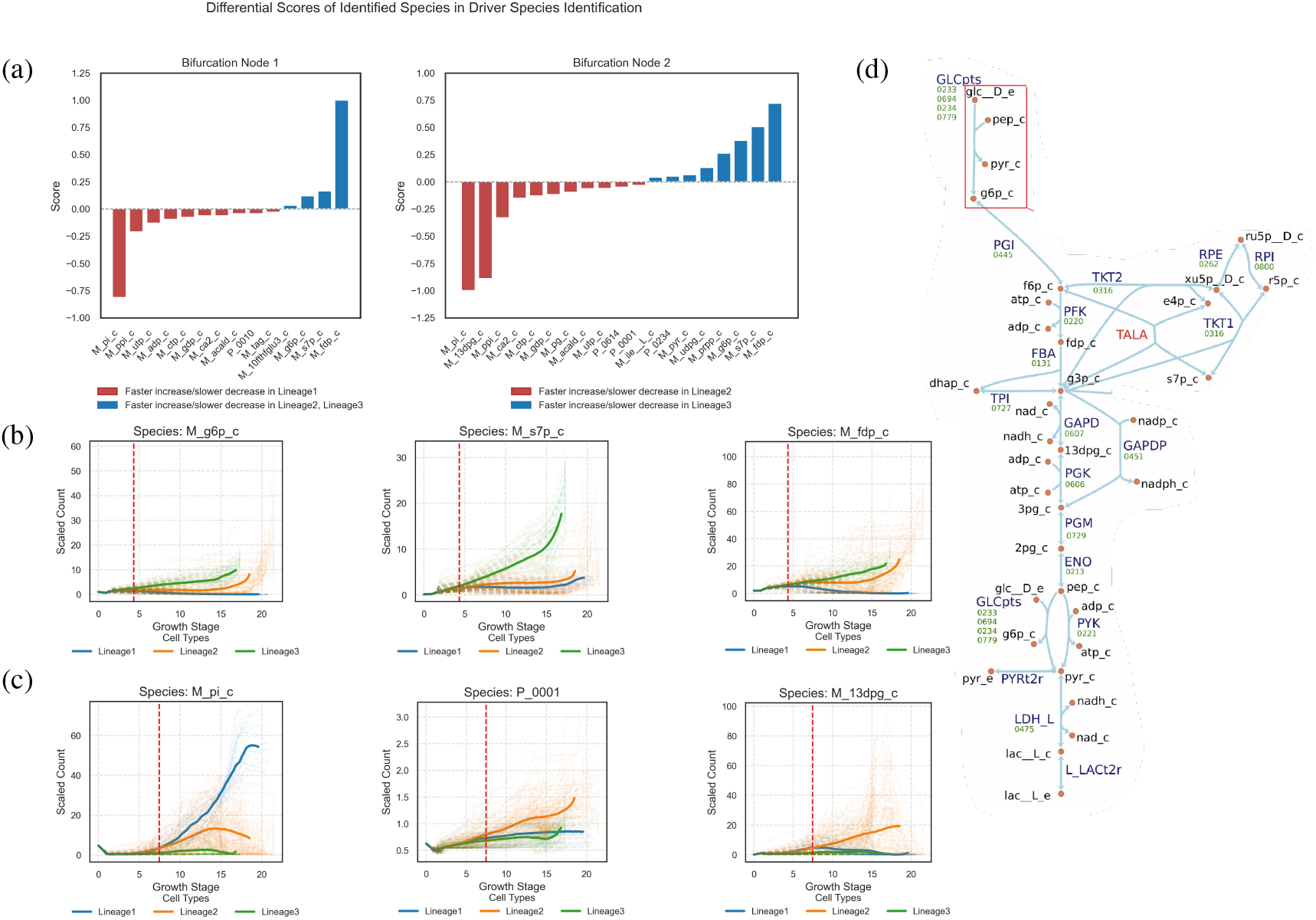
COTree identifies potential driver species underlying cell divergence. (a) summarizes species exhibiting distinct change directions at bifurcation nodes between lineages. (b) presents trajectories of glucose-6-phosphate (G6P), sedoheptulose 7-phosphate (S7P), and fructose-1,6-bisphosphate (FDP) identified at bifurcation node 1, where the dashed line indicates the bifurcation time. (c) shows trajectories of inorganic phosphate (Pi), the replication initiator protein DnaA/0001 (P_0001), and 1,3-bisphosphoglycerate (13DPG) identified at bifurcation node 2. (d) presents the major pathway in the glycolysis pathway of JCVI-syn3A.

## 6. Discussion

This paper introduces COTree, a statistical framework for analyzing and interpreting cell-resolved multi-omics trajectories. Whereas most existing trajectoryinference tools are designed for snapshot measurements from single-omics data, particularly transcriptomics, COTree integrates information across multiple omics layers, including genomics, transcriptomics, proteomics, and metabolomics, to characterize overall trends in cellular progression. It can also accommodate user-defined modalities and inter-modality relationships, supporting flexible modeling across diverse biological systems. Leveraging the temporal and cross-omics structure of the data, COTree combines a dual-target multi-omics autoencoder for learning low-dimensional representations with a trajectory principal tree that summarizes cellular dynamics. The resulting representations and tree structure support down-stream analyses including cell classification, fate prediction, developmental-time detection, and candidate driver-species identification. Applying COTree to a whole-cell model of the minimal cell JCVI-syn3A, with multi-omics measurements recorded at one-minute intervals, demonstrates its ability to characterize cellular progression, distinguish alternative cellular outcomes, and identify candidate molecular species associated with cell death and DNA replication.

Whole-cell models have already produced important biological insights, including previously unobserved DNA-replication dynamics in Mycoplasma genitalium and emergent patterns of mRNA turnover and cellular energy use in JCVI-syn3A (Karr et al., 2012; Thornburg et al., 2022). These studies illustrate the value of targeted investigation of particular biological processes and simulation outputs. COTree complements such process-focused analyses by jointly analyzing complete multi-omics trajectories and identifying their population-level progression structure without requiring a particular pathway or cellular outcome to be specified in advance.

In the JCVI-syn3A simulations, COTree identifies three major cell types, determines when their trajectories diverge, and prioritizes coordinated molecular changes associated with each outcome. Specifically, it detects early divergence toward glycolytic arrest, predicts this out-come before PEP depletion, and associates single- and multiple-replication outcomes with changes involving glycolytic intermediates, ATP, phosphate species, and DnaA. These results demonstrate how system-wide molecular dynamics organize into heterogeneous, long-term cellular outcomes. The model equations and COTree therefore answer complementary questions: the equations and simulation rules encode reaction- and process-level dynamics, whereas COTree summarizes the collective behaviors that emerge from their interactions across stochastic cellular trajectories. The molecular species and dependencies identified by COTree should be interpreted as hypothesis-generating associations that prioritize particular processes and time intervals for mechanistic inspection and targeted in silico perturbation, rather than as independent evidence of biochemical causality.

In its current implementation, the dual-target multi-omics autoencoder employs a fully connected neural network to preserve information from multi-omics trajectories. Future extensions could incorporate more advanced architectures, such as transformers, to enhance representation learning. Beyond the reconstruction and autoregressive losses, additional objectives, such as denoising or masking, may further improve robustness and interpretability. In constructing the trajectory principal tree, the current two-step algorithm updates lineages and detects bifurcation points separately; a unified framework may yield more accurate structural estimation. Although we illustrate four downstream analyses, additional applications are possible, including differential trajectory analyses across lineages, biological conditions, or perturbations. Exploring these extensions represents an important direction for future research.

Comprehensive whole-cell-model datasets are currently available for only a limited number of biological systems, with most mechanistic models focusing on relatively simple organisms. However, recent advances in whole-cell and broader virtual-cell modeling reflect growing interest in modeling more complex biological systems and predicting cellular responses to genetic, environmental, and therapeutic perturbations (Bunne et al., 2024). As these efforts generate increasingly diverse ensembles of cell-resolved trajectories, statistical methods such as COTree will become increasingly valuable for their analysis and interpretation. COTree is not intrinsically restricted to bacterial systems and can be applied whenever repeated, temporally resolved, multimodal cellular trajectories are available. Future advances in longitudinal multi-omics measurement may further broaden its applicability to experimental data, although adaptations may be required to accommodate measurement noise, missing modalities, and incomplete trajectories.

## SUPPLEMENTARY MATERIAL

### Supplement Materials

Implementation details and more results.

### Data and Code

The raw cell trajectory data from Thornburg et al. (2022) is available at https://github.com/BoYuan07/COTree_manuscript_code. The python package is available at https://github.com/BoYuan07/COTree. All analyses can be found under https://github.com/BoYuan07/COTree_manuscript_code.

