## Supplemental Materials for "COTree: A Statistical Framework for Deciphering Cell-Resolved Multi-Omics Trajectories"

#### S1 Multi-Omics Representation Interpretation

During the representation learning step, COTree compresses the input species space to a low-dimensional latent space. Although these latent representations capture current cell states and developmental progression, the embedded information can still be difficult to interpret. To overcome this challenge, we propose a new method to interpret the multi-omics representations from two different perspectives: one is from current cell species abundance and the other is from cell developmental progression. Specifically, we aim to interpret each coordinate of multi-omics representations by detecting the relevant species and estimating a prediction score.

**Functional Module Detection** To interpret an individual coordinate of the multi-omics representation, we seek to identify a group of molecular features that exhibit change patterns similar to that coordinate. For simplicity, we refer to such a group as a functional module corresponding to the coordinate. To detect a functional module, we search for a group of species whose encoder activations are highly correlated with the coordinate in the latent representation space. Specifically, we can consider the following optimization problem

$$\min_{F_k} -\frac{1}{m}\sum_{i=1}^{m}\frac{1}{T_i}\sum_{t=1}^{T_i}\frac{\left[f_e\left(X_{i,t}+F_k\right)-Z_{i,t}\right]^\top e_k}{\left\|f_e\left(X_{i,t}+F_k\right)-Z_{i,t}\right\|}+\eta_F\|F_k\|_1.$$

Here, $X_{i,t} \in \mathbb{R}^{d_D+d_P+d_M}$ is the concatenated vector of count vectors in DNA, $D_{i,t}$, protein, $P_{i,t}$, and metabolite, $M_{i,t}$. $f_e$ is the concatenated encoder of DT-MAE of different omics layers

$$f_e(D_{i,t}, P_{i,t}, M_{i,t}) = [f_e^D(D_{i,t})^\top, f_e^P(P_{i,t})^\top, f_e^M(M_{i,t})^\top]^\top = Z_{i,t}.$$

$e_k$ is the standard basis vector in the latent representation space with a 1 in the $k$-th coordinate and 0 in all the other coordinates. $F_k \in \mathbb{R}^{d_D+d_P+d_M}$ represents the change pattern of the functional module corresponding to the $k$-th coordinate. We add an $L_1$ penalty to promote the sparsity of $F_k$. The detected species in the functional module corresponding to the $k$-th coordinate are defined as the entries of $F_k$ whose absolute values exceed a threshold $\tau$:

$$\mathcal{S}_k = \left\{s \in \{1,\ldots,d_D+d_P+d_M\} \mid |F_{k,s}| > \tau\right\}.$$

Because the representation is a concatenation of representations from different omics layers, we can expect the species detected in each functional module to come from the same omics layer.

**Functional Module Prediction Score Estimation** Since the representations aim to capture the cell growth trend, we can also assess how important each coordinate of representation, corresponding to each functional module, is in predicting future species abundance. The importance in predicting individual species can be a direct measure of how well each functional module can capture the developmental progression of these species. To evaluate the importance in predicting future species abundance, we define the prediction score of functional module $k$ on species $s$ as the average relative perturbation when we mask the $k$-th coordinate of the representation

$$S(k|s) = \frac{1}{m}\sum_{i=1}^{m}\frac{1}{T_i}\sum_{t=1}^{T_i}\frac{\left\|h_d(Z_{i,t}^{\text{masked}(k)}, R_{i,t})_s - h_d(Z_{i,t}, R_{i,t})_s\right\|}{\|h_d(Z_{i,t}, R_{i,t})_s\|}$$

Here, $h_d$ is the concatenated decoder of prediction decoders in DT-MAE

$$h_d(Z_{i,t}, R_{i,t}) = [h_d^D(Z_{i,t}^D, Z_{i,t}^P, Z_{i,t}^M)^\top, h_d^P(Z_{i,t}^P, R_{i,t}, Z_{i,t}^M)^\top, h_d^M(Z_{i,t}^M, Z_{i,t}^P)^\top]^\top,$$

and $h_d(\cdot)_s$ denotes the predicted abundance of species $s$. $Z_{i,t}^{\text{masked}(k)}$ is the latent representation $Z_{i,t}$ with the $k$-th coordinate set to zero. The definition of $S(k|s)$ suggests that a high prediction score of functional module $k$ on species $s$ implies that functional module $k$ can well capture the growth trend of species $s$.

### S2 Implementation Details

#### S2.1 Input Preprocessing for DT-MAE

To stabilize the training of the neural network, we apply feature scaling to the raw count vectors for DNA ($D_{i,t}^{\text{raw}}$), protein ($P_{i,t}^{\text{raw}}$), metabolite ($M_{i,t}^{\text{raw}}$), and mRNA ($R_{i,t}^{\text{raw}}$). For protein, metabolite, and mRNA vectors, we scale the raw counts by their mean values:

$$P_{i,t} = \frac{P_{i,t}^{\text{raw}}}{\frac{1}{m}\sum_{i=1}^{m}\frac{1}{T_i}\sum_{t=1}^{T_i}P_{i,t}^{\text{raw}}}, \quad M_{i,t} = \frac{M_{i,t}^{\text{raw}}}{\frac{1}{m}\sum_{i=1}^{m}\frac{1}{T_i}\sum_{t=1}^{T_i}M_{i,t}^{\text{raw}}}, \quad R_{i,t} = \frac{R_{i,t}^{\text{raw}}}{\frac{1}{m}\sum_{i=1}^{m}\frac{1}{T_i}\sum_{t=1}^{T_i}R_{i,t}^{\text{raw}}}.$$

The DNA vectors are scaled by the mean value multiplied by 0.5:

$$D_{i,t} = \frac{D_{i,t}^{\text{raw}}}{\frac{1}{2m}\sum_{i=1}^{m}\frac{1}{T_i}\sum_{t=1}^{T_i}D_{i,t}^{\text{raw}}}.$$

#### S2.2 Implementation Details for DT-MAE

Our representation model DT-MAE consists of three components: a multi-omics encoder module, a reconstruction decoder, and a prediction decoder. Before describing the architectures of the encoders and decoders, we first define a basic layer function that is used throughout the model:

$$\sigma(X) = \text{LeakyReLU}(\text{LayerNorm}(W^T X + b)), \tag{S2.1}$$

where $W$ and $b$ are learnable parameters. Layer normalization (Ba et al., 2016) is applied to stabilize the learning process and is defined as:

$$\text{LayerNorm}(X) = \frac{X - \mathbb{E}(X)}{\sqrt{\text{Var}(X) + \epsilon}} \cdot \gamma + \beta,$$

where $\gamma$ and $\beta$ are learnable scaling and shifting parameters. The Leaky ReLU activation function is defined as:

$$\text{LeakyReLU}(x) = \begin{cases} x, & \text{if } x \geq 0, \\ \alpha x, & \text{if } x < 0, \end{cases}$$

with $\alpha = 0.05$ in our implementation. Compared with the ReLU activation function, the Leaky ReLU function allows a small gradient for negative inputs to mitigate the dying ReLU problem (Xu et al., 2015).

The encoder module consists of separate encoders for each omics layer, where the scaled count vectors $D_{i,t}$, $P_{i,t}$, and $M_{i,t}$ are used as input:

$$Z_{i,t}^D = f_e^D(D_{i,t}), \qquad Z_{i,t}^P = f_e^P(\text{LayerNorm}(P_{i,t})), \qquad Z_{i,t}^M = f_e^M(\text{LayerNorm}(M_{i,t})),$$

The encoders $f_e^D$, $f_e^P$, and $f_e^M$ are each implemented as two-layer networks, where each layer follows the formulation in Eq. S2.1.

The reconstruction decoder reconstructs the counts $D_{i,t}$, $P_{i,t}$, and $M_{i,t}$ using representations $Z_{i,t}^D$, $Z_{i,t}^P$ and $Z_{i,t}^M$

$$\hat{D}_{i,t} = g_d^D(Z_{i,t}^D), \qquad \hat{P}_{i,t} = g_d^P(Z_{i,t}^P), \qquad \hat{M}_{i,t} = g_d^M(Z_{i,t}^M),$$

where $g_d^D$, $g_d^P$ and $g_d^M$ mirror the structure of $f_e^D$, $f_e^P$ and $f_e^M$.

The prediction decoder takes the concatenated vector of representations as input and predicts the counts at the next time point $t+1$

$$\tilde{D}_{i,t+1} = h_d^D(Z_{i,t}^D, Z_{i,t}^P, Z_{i,t}^M), \qquad \tilde{P}_{i,t+1} = h_d^P(Z_{i,t}^P, R_{i,t}, Z_{i,t}^M), \qquad \tilde{M}_{i,t+1} = h_d^M(Z_{i,t}^M, Z_{i,t}^P).$$

The functions $h_d^D$, $h_d^P$, and $h_d^M$ are each implemented as two-layer networks, where each layer follows the formulation in Eq. S2.1.

The model is trained with 200 epochs using the Adam optimizer with learning rate 0.005 (Kingma and Ba, 2014), and the size of each batch is 200. Given a small batch of cells $\mathcal{S}$, the batch-wise loss is defined as

$$\mathcal{L}_{\text{RL}}^{\text{batch}} = \frac{\eta_g}{|\mathcal{S}|} \sum_{(i,t)\in\mathcal{S}} \left[ \gamma_D \|D_{i,t} - \hat{D}_{i,t}\|^2 + \gamma_P \|P_{i,t} - \hat{P}_{i,t}\|^2 + \gamma_M \|M_{i,t} - \hat{M}_{i,t}\|^2 \right]$$
$$+ \frac{\eta_h}{|\mathcal{S}|} \sum_{(i,t)\in\mathcal{S}} \left[ \gamma_D \|D_{i,t+1} - \tilde{D}_{i,t+1}\|^2 + \gamma_P \|P_{i,t+1} - \tilde{P}_{i,t+1}\|^2 + \gamma_M \|M_{i,t+1} - \tilde{M}_{i,t+1}\|^2 \right].$$

where

$$\gamma_D + \gamma_P + \gamma_M = 1, \qquad \eta_g + \eta_h = 1.$$

Hyperparameters are optimized with Optuna using a multi-objective formulation. We divide the cells into training and validation sets. In each trial, Optuna samples a candidate hyperparameter configuration, trains the DT-MAE model on the training set, and evaluates three objectives on the validation set:

1. the prediction loss, which is defined as an unweighted sum of per-modality prediction losses on held-out cells;

2. the reconstruction loss, which is defined as an unweighted sum of per-modality reconstruction losses on held-out cells;

3. a manifold-smoothness penalty evaluated over all cells, which is defined as

$$\sum_{i,j} \sum_{t,t'} \exp\left( -\frac{\|X_{i,t} - X_{j,t'}\|^2}{\psi^2} \right) \|Z_{i,t} - Z_{j,t'}\|^2,$$

   where $\psi$ is the median of all pairwise distances in the input space. This penalty encourages cell states that are similar in the input space to have locally smooth latent embeddings and is essential for the selection of $\eta_h$ and $\eta_g$.

After all 100 trials are completed, Optuna returns the Pareto front, defined as the set of trials for which no other trial performs at least as well on all three objectives and strictly better on at least one. Among the trials on the Pareto front, we select the model with the lowest sum of the validation prediction loss and validation reconstruction loss. Table S1 lists the searched hyperparameters. Loss weights are sampled log-uniformly. A single hidden-layer dimension per modality is shared across that modality's encoder, decoder and predictor head. Learning rate and batch size are selected as $5 \times 10^{-3}$ and 200 respectively.

Table S1: Hyperparameter search space.

| Hyperparameter | Distribution | Range / choices |
|---|---|---|
| $\gamma_D,\ \gamma_M,\ \gamma_P$ (raw) | log-uniform | [0.1, 1.0], normalized to sum 1 |
| $\eta_g$ | log-uniform | [0.1, 1.0] |
| hidden layer dimension (per modality) | categorical | {128, 256} |
| latent dim $M$ | categorical | {10, 30, 50} |
| latent dim $P$ | categorical | {20, 50, 80} |
| latent dim $D$ | categorical | {5, 10, 15} |
| learning rate | fixed | $5 \times 10^{-3}$ (Adam) |
| batch size | fixed | 200 |

#### S2.3 Implementation Details for Functional Module Detection

In the functional module detection, we consider the optimization problem

$$\min_{F_k} -\frac{1}{m}\sum_{i=1}^{m}\frac{1}{T_i}\sum_{t=1}^{T_i}\frac{\left[f_e\left(X_{i,t}+F_k\right)-Z_{i,t}\right]^\top e_k}{\|f_e\left(X_{i,t}+F_k\right)-Z_{i,t}\|}+\eta_F\|F_k\|_1$$

where the first term measures the negative cosine similarity between the latent direction induced by adding $F_k$ in the input space and the target direction $e_k$. To simplify the notation, we denote the target direction in the input space as $u_k$ and let

$$\rho(F_k;u_k)=\frac{1}{m}\sum_{i=1}^{m}\frac{1}{T_i}\sum_{t=1}^{T_i}\frac{\left[f_e\left(X_{i,t}+F_k\right)-Z_{i,t}\right]^\top e_k}{\|f_e\left(X_{i,t}+F_k\right)-Z_{i,t}\|}.$$

Additionally, an $L_1$ penalty is applied to enforce sparsity in the feature selection and we select $\eta_F = 0.001$ in the implementation. It has frequently been observed that the $L_1$ penalty can shrink the coefficients of irrelevant features exactly to zero under many settings (Tibshirani, 1996). However, when applying the $L_1$ penalty for parameter selection in neural networks, the coefficients typically approach values very close to zero rather than becoming exactly zero due to the complex loss landscape and numerical fluctuations. Under this scenario, pruning is usually performed using a global threshold $\tau$ (Han et al., 2015; Wang et al., 2020). Similarly, we define the detected species in the functional module corresponding to the $k$-th coordinate as the entries of $F_k$ whose absolute values exceed a threshold $\tau$:

$$\mathcal{S}_k = \{s \in \{1, \ldots, d_D + d_P + d_M\} \mid |F_{k,s}| > \tau\}.$$

To determine the optimal value of $\tau$, we adopt a grid search procedure, which aims to find a value of $\tau$ such that filtering features according to this threshold has a negligible effect on the optimization objective. We define the set of candidates as $\mathcal{G}_\tau = \{\tau_n\}_{n=1}^N$ where $\tau_0 = \min_s |F_{k,s}|$ and $\tau_N = \max_s |F_{k,s}|$. For each $\tau_n$, we denote the set of species that are not identified in functional module $k$ as

$$\mathcal{N}_k^{\tau_n} = \{s \in \{1, \ldots, d_D + d_P + d_M\} \mid |F_{k,s}| \leq \tau_n\}.$$

and the vector obtained by masking the entries of $F_k$ corresponding to $\mathcal{N}_k^{\tau_n}$ as

$$F_{k,s}^{\text{masked}(\tau_n)} = \begin{cases} 0, & s \in \mathcal{N}_k^{\tau_n}, \\ F_{k,s}, & s \in \mathcal{S}_k^{\tau_n}. \end{cases}$$

To assess the importance of the species in $\mathcal{N}_k^{\tau_n}$ for maximizing the correlation, we measure the change in correlation resulting from masking the species in $\mathcal{N}_k^{\tau_n}$

$$\Delta\rho^{\tau_n} = \rho(F_k; u_k) - \rho(F_k^{\text{masked}(\tau_n)}; u_k).$$

A small value of $\Delta\rho^{\tau_n}$ suggests that species in $\mathcal{N}_k^{\tau_n}$ contribute negligibly to the estimation of $F_k$. When $\tau_n$ is larger than the optimal threshold, the first rapid increase in $\Delta\rho^{\tau_n}$ can be expected due to the masking of informative species. To identify the appropriate value of $\tau$, we plot $\Delta\rho^{\tau_n}$ as a function of $\tau_n$ and select the elbow point within the interval $[\tau_0, \tau_{\max}]$, where $\tau_{\max}$ is chosen to avoid over-pruning

$$\tau_{\max} = \max\left\{\tau_n \in \mathcal{G}_\tau : \frac{\Delta\rho^{\tau_n}}{\rho(F_k; u_k)} < 0.1\right\}.$$

The optimization problem is solved with 100 epochs using the Adam optimizer with learning rate 0.01, and the size of each batch is 1000.

#### S2.4 Implementation Details for Trajectory Tree Construction

**Estimation of Tree Topology** The trajectory tree construction consists of two steps. In the first step of piecewise linear tree estimation, COTree identifies tree nodes by applying

$k$-means clustering to cell states and constructs a minimum spanning tree. We set the number of clusters for intermediate states to 15 and for final states to 6. The minimum spanning tree is estimated using the `minimum_spanning_tree` function in the Python package `scipy`.

Then we adopt an over-split-then-trim strategy on the constructed tree. Starting from this over-split tree, we greedily remove redundant lineages one at a time. Let $\mathcal{T}^{(k)}$ denote the tree at iteration $k$, and let $\mathcal{J}^{(k)}$ denote its set of lineages on $\mathcal{T}^{(k)}$. We define the validation reconstruction distance as

$$D(\mathcal{T}) = \frac{1}{m} \sum_{i=1}^{m} \min_{l_j \in \mathcal{J}^{(k)}} \left\{ \sum_{t=1}^{T_i} \| Z_{i,t} - l_j(\lambda_j(Z_{i,t})) \|^2 \right\}.$$

where $l_j(\lambda_j(Z_{i,t}))$ is the projection of $Z_{i,t}$ onto lineage $l_j$. At each iteration, we remove the lineage whose exclusion produces the smallest increase in validation reconstruction distance:

$$l^{(k)} = \arg\min_{l \in \mathcal{J}^{(k)}} \left\{ D\left(\mathcal{T}^{(k)} \setminus l\right) - D\left(\mathcal{T}^{(k)}\right) \right\},$$

and update the tree as

$$\mathcal{T}^{(k+1)} = \mathcal{T}^{(k)} \setminus l^{(k)}.$$

The pruning procedure stops when removing any remaining lineage would substantially increase the reconstruction distance, defined as an increase of more than 10%.

After pruning, the nodes along each surviving lineage are refitted using nearby cells. For each lineage $j$, we first identify the set of cells $\mathcal{I}_j$ assigned to that lineage based on their projection distances. For $i \in \mathcal{I}_j$, we then assign each point $Z_{i,t}$ to its nearest node on lineage $j$. The position of node $q$ is updated as the mean of the points assigned to it:

$$v_{j,q} = \frac{1}{|\mathcal{A}_{j,q}|} \sum_{(i,t) \in \mathcal{A}_{j,q}} Z_{i,t},$$

where

$$\mathcal{A}_{j,q} = \left\{ (i,t) : i \in \mathcal{I}_j,\ q = \arg\min_{q'} \| Z_{i,t} - v_{j,q'} \|_2^2 \right\}.$$

Here, $|\mathcal{A}_{j,q}|$ is the number of points assigned to node $q$.

In the second step, an iterative algorithm is applied to refine the tree. We select the maximum number of iterations as 20, and the algorithm stops if the relative loss falls below 0.1. In each iteration, COTree updates each lineage via a time-informed principal curve algorithm (Step 2A), and reconstructs the shared branches (Step 2B). In Step 2A, COTree first estimates the growth stage by projecting each point $Z_{i,t}$ onto the lineage, which is implemented using the Python package `pcurvepy2`. The growth stage is then adjusted using isotonic regression to ensure consistency with the ordering of real time. After obtaining the growth stage estimation, COTree applies the LOESS algorithm to refit the curve, where the weight is calculated as

$$w_j(Z_{i,t}) = \gamma \exp\left(-\frac{\|Z_{i,t} - l_j(\lambda_j(Z_{i,t}))\|^2}{\psi_1}\right) + (1-\gamma)\mathbf{1}\left\{j = \arg\min_k \sum_{t=t_0}^{T_i} \|Z_{i,t} - l_k(\lambda_k(Z_{i,t}))\|^2\right\},$$

$\psi_1$ is set to the median of the point-level squared distances across all point-lineage pairs. The LOESS algorithm is implemented using the Python package `skmisc`. In iteration step 2B, COTree detects the bifurcation point and reconstructs the shared branch. We initialize the first point $\lambda_0 = 0$. Given the $k$-th bifurcation point with $\lambda_k$ and the set of lineages $\mathcal{J}_k$ that do not diverge at $\lambda_k$, the next $\lambda_{k+1}$ is selected based on the distance of each lineage to the average

$$\lambda_{k+1} = \min\left\{\lambda \in \Lambda \;\middle|\; \lambda > \lambda_k \quad \text{and} \quad \max_{l_j \in \mathcal{J}_k} \left\| l_j(\lambda) - \frac{1}{|\mathcal{J}_k|} \sum_{l_{j'} \in \mathcal{J}_k} l_{j'}(\lambda) \right\|^2 > \tau \right\},$$

The divergence threshold is chosen as

$$\tau = \nu^2 R^2,$$

with $\nu = 0.05$, where $R^2$ is the maximum pairwise squared distance between unmerged lineage curves.

The parameter $\gamma$ is selected based on assignment entropy. For each candidate value of $\gamma \in [0, 1]$, the projection distances from each cell trajectory to each lineage are converted into soft lineage-assignment probabilities using a Gaussian kernel with a median-heuristic bandwidth. We then compute the entropy of the resulting lineage-assignment distribution for each cell. Finally, we select the value of $\gamma$ that yields the lowest mean assignment entropy across cell trajectories.

**Estimation of Non-Tree Topology** Cell trajectories may also follow non-tree topologies, since biological state transitions can involve cycles, convergence, and reversibility (Tanay and

Regev, 2017). COTree can estimate non-tree structures for complex cellular dynamics with the following steps.

In the first step, COTree constructs a minimum spanning tree from the adjacency matrix using the same procedure described in Section 3.3.1 and then gradually adds edges back to the graph. Let $A_{u,v}$ denote the adjacency value between nodes $u$ and $v$. We define the direct edge weight between $u$ and $v$ as

$$w(u, v) = \frac{1}{A_{u,v}}.$$

Starting with the pair of nodes $(u, v)$ having the largest adjacency value but no direct edge in the tree, COTree identifies the shortest path connecting these two nodes and calculates the shortest-path distance $d(u, v)$, defined as the sum of the edge weights along this path. If $d(u, v)$ is substantially greater than the direct edge weight $w(u, v)$ and satisfies

$$d(u, v) \geq \kappa w(u, v), \qquad \kappa > 1,$$

COTree adds the edge $(u, v)$ to the graph. We use $\kappa = 1.5$ in practice. After each edge insertion, COTree recomputes the shortest-path distances using the updated graph. The procedure stops when no remaining candidate edge satisfies the criterion.

After the edge-insertion process, COTree identifies all simple paths originating from the root, where a simple path is defined as a path that does not visit any node more than once. COTree first identifies all paths from the root to nodes of degree 1. For each remaining region containing a cycle, COTree identifies the longest simple path from the root through the cycle. It then extracts an additional simple path that traverses the remaining portion of the cycle not covered by the first path. These paths are treated as separate lineages.

In the second step, each collected lineage is refined using the same iterative principal-curve algorithm described in Steps 2A–2B, with an additional backward convergence-detection step. Starting from the largest shared growth stage $\lambda_{\max}$ and moving toward the root, COTree calculates the distance between the position $l_j(\lambda)$ of each lineage $j$ and the average position across all currently included lineages. If the maximum distance remains below the divergence threshold $\tau$, the lineage segments over $[\lambda, \lambda_{\max}]$ are averaged to form a shared branch. If one lineage has a distance greater than $\tau$, it is removed from the group, and the lineage-to-average distances are recomputed for the remaining lineages. The remaining lineages are merged if their maximum distance is below $\tau$. This process continues until no pair of lineages remains sufficiently close to be merged into a shared branch. The forward bifurcation-detection and backward convergence-detection procedures allow COTree to construct trajectory structures that include loops.

#### S2.5 Implementation Details for Driver Species Identification

To identify the species that drive directional shifts between two trajectory segments $\mathcal{I}_{j_1}(\lambda_1|\tilde{\lambda}_1)$ and $\mathcal{I}_{j_2}(\lambda_2|\tilde{\lambda}_2)$, we infer the direction of change in species space associated with each segment by solving the objective function

$$\max_{U_1,U_2} \; \rho(U_1;\mathcal{I}_{j_1}(\lambda_1|\tilde{\lambda}_1)) + \rho(U_2;\mathcal{I}_{j_2}(\lambda_2|\tilde{\lambda}_2)) - \eta\|U_1 - U_2\|_1,$$

where

$$\rho(U;\mathcal{I}_j(\lambda|\tilde{\lambda})) := \frac{1}{|\mathcal{I}_j(\lambda|\tilde{\lambda})|} \sum_{Z_{i,t}\in\mathcal{I}_j(\lambda|\tilde{\lambda})} \frac{(f_e(X_{i,t} + \langle \Delta X_{i,t},\, U\rangle U) - Z_{i,t})^\top \Delta Z_{i,t}}{\|f_e(X_{i,t} + \langle \Delta X_{i,t},\, U\rangle U) - Z_{i,t}\|\|\Delta Z_{i,t}\|},$$

$$\Delta X_{i,t} = X_{i,t+\Delta t} - X_{i,t},$$

$$\Delta Z_{i,t} = Z_{i,t+\Delta t} - Z_{i,t}.$$

In the implementation, we reparametrize the problem in terms of the shared direction $a$ and the difference $b$ between the two directions,

$$a = U_1, \qquad b = U_2 - U_1,$$

so that $U_1 = a$ and $U_2 = a + b$. This leads to the following optimization problem

$$\max_{a,b} \; \rho(a;\mathcal{I}_{j_1}(\lambda_1|\tilde{\lambda}_1)) + \rho(a+b;\mathcal{I}_{j_2}(\lambda_2|\tilde{\lambda}_2)) - \eta\|b\|_1,$$

where the $L_1$ penalty on $b = U_2 - U_1$ promotes sparsity in the difference between the two directions. We initialize $a^{(0)}$ and $b^{(0)}$ as

$$a^{(0)} = \frac{1}{|\mathcal{I}_{j_1}(\lambda_1|\tilde{\lambda}_1)|} \sum_{Z_{i,t}\in\mathcal{I}_{j_1}(\lambda_1|\tilde{\lambda}_1)} \Delta X_{i,t},$$

$$b^{(0)} = \frac{1}{|\mathcal{I}_{j_2}(\lambda_2|\tilde{\lambda}_2)|} \sum_{Z_{i,t}\in\mathcal{I}_{j_2}(\lambda_2|\tilde{\lambda}_2)} \Delta X_{i,t} - a^{(0)}.$$

We select $\eta = 0.001$ in the analysis, and the optimization problem is solved using the Adam optimizer with learning rate 0.01 and 1000 epochs. In the experiment, $\Delta t$ is selected to be 20, $\tilde{\lambda} = \lambda + 0.5$.

The set of driver species is identified as the species whose entry of $b$ has absolute value exceeding the threshold $\tau$

$$\mathcal{S} = \{s \in \{1, \ldots, d_D + d_P + d_M\} \mid |b_s| > \tau\}.$$

Since $b = U_2 - U_1$, this is equivalent to selecting species with a large absolute difference $|U_{1,s} - U_{2,s}|$ between the two estimated directions, as defined in the main text. We determine $\tau$ by identifying how many species are needed to recover the directional shift encoded by $b$, and then setting $\tau$ to retain exactly those species. We first rank the species in decreasing order of coefficient magnitude, so that $|b_{(1)}| \geq |b_{(2)}| \geq \cdots \geq |b_{(K)}|$, where $K = d_D + d_P + d_M$. For $k \in \{0, 1, \ldots, K\}$, let $b^{(k)}$ denote the vector that retains the $k$ largest-magnitude entries of $b$ and sets the remaining entries to zero. We assess how well the top-$k$ species reproduce the direction $U_2 = a + b$ by the recovered-gain curve

$$R(k) = \frac{\rho\Big(a + b^{(k)}; \mathcal{I}_{j_2}(\lambda_2|\tilde{\lambda}_2)\Big) - \rho\Big(a; \mathcal{I}_{j_2}(\lambda_2|\tilde{\lambda}_2)\Big)}{\rho\Big(a + b; \mathcal{I}_{j_2}(\lambda_2|\tilde{\lambda}_2)\Big) - \rho\Big(a; \mathcal{I}_{j_2}(\lambda_2|\tilde{\lambda}_2)\Big)},$$

which measures the fraction of the achievable alignment gain. By construction $R(0) = 0$ and $R(K) = 1$, and $R(k)$ typically rises with diminishing returns as informative species are added.

The number of driver species $k^\star$ is chosen as the knee of $R(k)$, detected with the Kneedle algorithm (Satopaa et al., 2011) under a concave, increasing profile. To prevent a long, weakly contributing tail from being selected, $k^\star$ is capped at the smallest size that already recovers a target fraction $p$ of the achievable gain,

$$k_{\text{target}} = \min\{k : R_{\text{mono}}(k) \geq p\}, \qquad k_{\text{sel}} = \min\{k^\star, k_{\text{target}}\},$$

with $p = 0.95$ in our analysis; if no knee is detected, we set $k_{\text{sel}} = k_{\text{target}}$, and $\tau$ is set as the smallest coefficient value in the selected set.

#### S2.6 Simulation Settings

To benchmark trajectory recovery against a known ground truth, we generate synthetic multi-omics trajectory datasets with known latent tree topologies. In each simulation, cells progress along predefined lineages in a low-dimensional latent space. Observed omics features are then generated from the latent states through modality-specific emission models. Because the true latent tree topology and lineage assignments are known by construction, the inferred trajectories can be quantitatively evaluated against the ground truth.

##### Latent Tree Construction

We simulate a latent trajectory tree in a 10-dimensional latent space. Here we use two lineages as an example, while trees with more lineages follow a similar procedure. For lineage $j = 1, 2$, the latent path is defined as

$$l_j(t) = \sum_{\ell=1}^{N_j} d_{j,\ell} r_{j,\ell} \left[\min\left(t, \tau_{j,\ell}^{\text{end}}\right) - \tau_{j,\ell}^{\text{start}}\right]_+ ,$$

where

$$[x]_+ = \max(x, 0),$$

$N_j$ is the number of linear segments of lineage $j$, $d_{j,\ell} \in \mathbb{R}^{10}$ is the unit direction vector of segment $\ell$ for lineage $j$, $r_{j,\ell} > 0$ is its progression rate, and $\tau_{j,\ell}^{\text{start}}$ and $\tau_{j,\ell}^{\text{end}}$ define the active time window of the segment. We set $\tau_{j,1}^{\text{start}} = 0$ and $\tau_{j,\ell}^{\text{end}} = \tau_{j,\ell+1}^{\text{start}}$ for $\ell = 1, \ldots, N_j - 1$, so that $l_j$ is continuous and all lineages share the common root $l_j(0) = 0$. The directions of the linear segments change smoothly along the trajectory, producing a curved path in latent space.

One lineage is selected to be the reference lineage, which we index by $j = 1$. Its path is constructed from a sequence of segment directions that gradually rotate from one latent direction $f_0$ toward a second latent direction $f_1$, where $f_0$ and $f_1$ are orthonormal vectors in $\mathbb{R}^{10}$. Specifically, for the linear segments $\ell \in \{1, \ldots, N_1\}$ that belong to lineage 1, we define

$$u_{1,\ell} = \frac{\ell}{N_1},$$

and compute the unnormalized segment direction as

$$\widetilde{d}_{1,\ell} = \cos\left(\frac{\pi}{2} u_{1,\ell}\right) f_0 + \sin\left(\frac{\pi}{2} u_{1,\ell}\right) f_1 + a_1 \sin\left(2\pi u_{1,\ell}\right) w_1 + a_2 \sin\left(5\pi u_{1,\ell}\right) w_2.$$

Here, $w_1$ and $w_2$ are unit directions orthogonal to $f_0$ and $f_1$, which introduce additional curvature, and $a_1$ and $a_2$ control the strengths of the two nonlinear bends. The final segment direction is obtained by normalization:

$$d_{1,\ell} = \frac{\widetilde{d}_{1,\ell}}{\left\|\widetilde{d}_{1,\ell}\right\|_2}.$$

Next, a bifurcation time $t_b$ is selected as the time at which the other lineage separates from the reference lineage. We take $t_b$ to be the end time of segment $\ell = N_b$

$$t_b = \tau^{\text{end}}_{1,N_b}, \quad N_b \in \{1, \ldots, N_1 - 1\}.$$

For $t \leq t_b$, lineage 2 shares the path of lineage 1. We have

$$d_{2,\ell} = d_{1,\ell}, \qquad r_{2,\ell} = r_{1,\ell}, \qquad \tau^{\text{start}}_{2,\ell} = \tau^{\text{start}}_{1,\ell}, \qquad \tau^{\text{end}}_{2,\ell} = \tau^{\text{end}}_{1,\ell}, \qquad \ell = 1, \ldots, N_b.$$

After bifurcation, lineage 2 peels away from lineage 1. The direction $d_{1,N_b}$, is the branch-continuation direction of previous path. We select a fan-out direction $f_2$ for the peeling lineage, chosen to be orthogonal to $d_{1,N_b}$. The unnormalized branch direction is then generated by rotating from the branch-continuation direction toward the fan-out direction:

$$\widetilde{d}_{2,\ell} = \cos\left(\phi\, u^r_{2,\ell}\right) d_{1,N_b} + \sin\left(\phi\, u^r_{2,\ell}\right) f_2 + c_1 \sin\left(2\pi u_{2,\ell}\right) g_1 + c_2 \sin\left(5\pi u_{2,\ell}\right) g_2, \quad \ell = N_b + 1, \ldots, N_2,$$

where

$$u_{2,\ell} = \frac{\ell - N_b - 1}{N_2 - N_b - 1} \in [0, 1], \qquad N_b \leq N_2 - 2.$$

Here, $\phi \in (0, \pi/2]$ is the fan-out angle, which controls how far the lineage deviates from the branch, and $r > 0$ controls the sharpness of early divergence: because $u_{2,\ell} \leq 1$, smaller values of $r$ make $u^r_{2,\ell}$ grow faster right after the bifurcation and therefore cause the lineage to separate more rapidly. The vectors $g_1$ and $g_2$ are unit directions orthogonal to both $d_{1,N_b}$ and $f_2$, and $c_1$ and $c_2$ control the strengths of the two nonlinear bends. The final branch direction is normalized as

$$d_{2,\ell} = \frac{\widetilde{d}_{2,\ell}}{\left\|\widetilde{d}_{2,\ell}\right\|_2}.$$

In the case with $J$ lineages, $J - 1$ bifurcation times are selected and one lineage peels away at each of them, each time using the construction above with the currently shared branch playing the role of lineage 1. This construction creates a ground-truth nonlinear latent trajectory tree with a known topology.

**Cell Latent Representation Construction**

Each cell is assumed to follow the path of one lineage on the tree, up to cell-specific variability. Suppose cell $i$ is assigned to lineage $j$. The $q$-th coordinate of its latent state at time $t$ is generated according to

$$Z_{i,t,q} \sim \mathcal{N}\big([l_j(t)]_q,\, [l_j(t)]_q^2 \sigma_{z,q}^2\big)\,, \qquad q = 1, \ldots, 10.$$

Here $l_j(t)$ denotes the latent path of lineage $j$, $[\,\cdot\,]_q$ extracts its $q$-th coordinate, and $\sigma_{z,q}$ is the coefficient of variation, so that it controls the level of cell variability in latent dimension $q$. The latent representation of cell $i$ is $Z_{i,t} = (Z_{i,t,1}, \ldots, Z_{i,t,10})^\top$.

**Observed Data Simulation**

Given the latent trajectory, modality-specific emission models map each latent state $Z_{i,t}$ to observed species values. To simulate data resembling protein and metabolite measurements, we use a log-link compositional count model with Poisson sampling. For modality $k$, each species $s$ has a baseline parameter $\alpha_s^k$, and the latent state is mapped to species-level linear predictors through a modality-specific loading matrix $W^k$. Specifically, we define

$$\theta_{i,s}^k = \alpha_s^k + \left(W^k Z_{i,t}\right)_s.$$

Species relative abundances are then obtained by applying a softmax transformation:

$$p_{i,\cdot}^k(t) = \mathrm{softmax}\left(\theta_{i,\cdot}^k(t)\right).$$

For each cell $i$, a modality-specific library size is sampled as

$$N_i^k \sim \mathrm{LogNormal}\left(\mu_{\mathrm{lib}}^k, \sigma_{\mathrm{lib}}^k\right),$$

and the expected count vector for cell $i$ at time $t$ in modality $k$ is given by

$$\mu_{i,\cdot}^k(t) = N_i^k\, p_{i,\cdot}^k(t).$$

The observed counts are then sampled from a Poisson distribution:

$$X^k_{i,t,s} \sim \text{Poisson}\left(\mu^k_{i,s}(t)\right).$$

In the experiment, we simulate two modalities with 100 and 200 species, respectively.

##### Evaluation Metrics

Each simulation returns the ground-truth terminal lineage labels, bifurcation times, latent positions, and modality-specific feature matrices. Lineage assignment accuracy is measured using the adjusted Rand index (ARI) (Hubert and Arabie, 1985). Topology recovery is assessed by comparing the inferred number of lineages with the true number of lineages, as well as by comparing the inferred bifurcation ordering with the ground truth.

To evaluate the robustness of COTree, we apply the full COTree pipeline, including hyperparameter tuning of DT-MAE, to the simulated data. We perform simulation sweeps across several scenarios, including varying numbers of initial clusters, varying numbers of lineages, observation noise levels by changing $\sigma_z$, lineage imbalance, and developmental rate differences by changing $r_\ell$. Metrics are aggregated across 300 simulation replicates. In the simulation, the total number of cells is set to 40 times the number of lineages.

### S3 Supplemental Figures

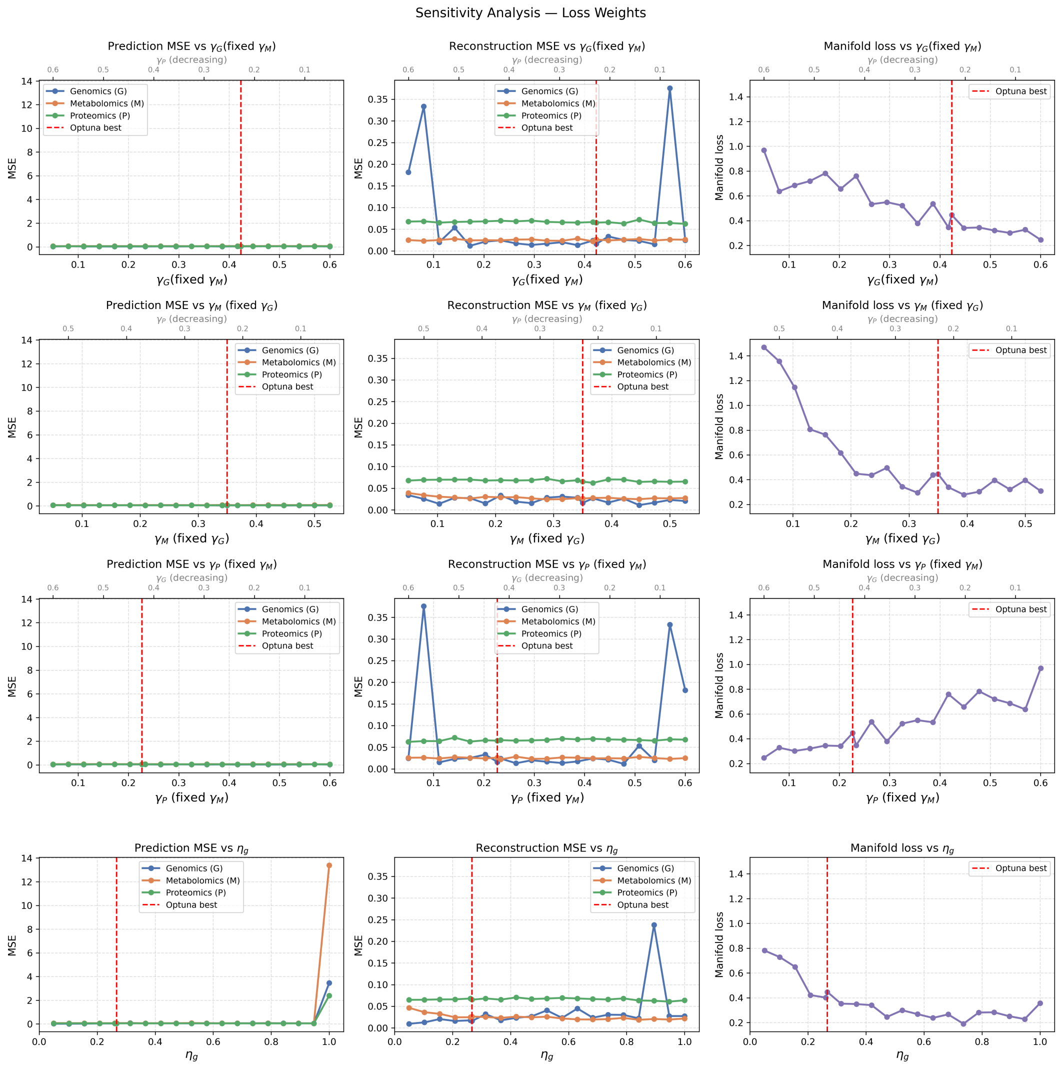


Figure S1: **Sensitivity analysis of loss weights $\gamma_D$, $\gamma_P$, $\gamma_M$, $\eta_h$, and $\eta_g$.** We vary one parameter at a time while fixing the remaining parameters at their optimal values. Prediction loss, reconstruction loss, and manifold smoothness loss are evaluated under each setting. The weights are constrained to satisfy $\gamma_D + \gamma_P + \gamma_M = 1, \eta_h + \eta_g = 1$.

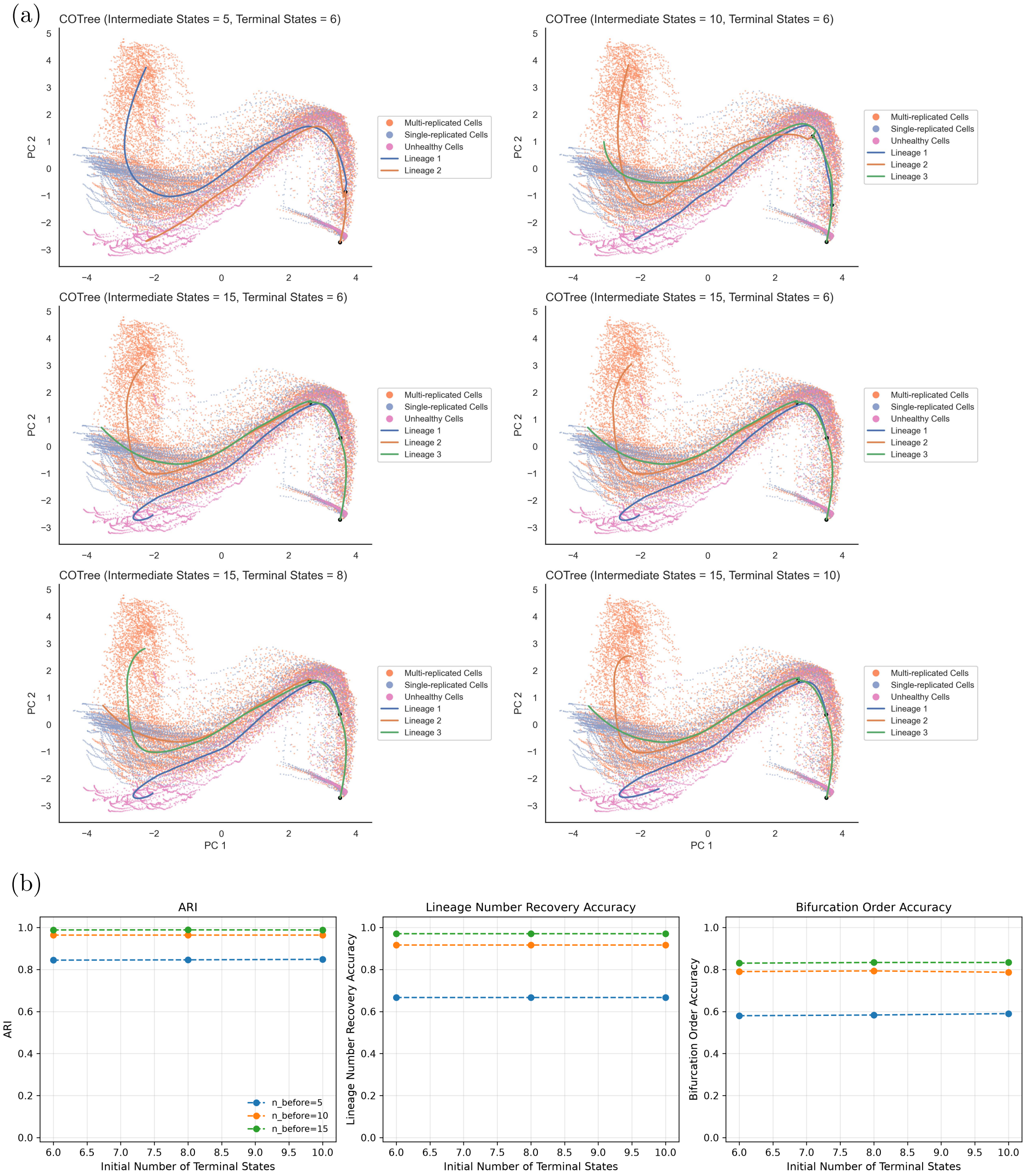


Figure S2: **Reconstructed trajectory trees with different numbers of initial clusters.** Figure (a) compares reconstructed trajectory trees initialized with different numbers of intermediate and terminal nodes. Figure (b) compares reconstruction performance with varied numbers of intermediate and terminal nodes in the simulation, including lineage classification accuracy, recovery of the correct number of lineages, and recovery of the correct bifurcation ordering.

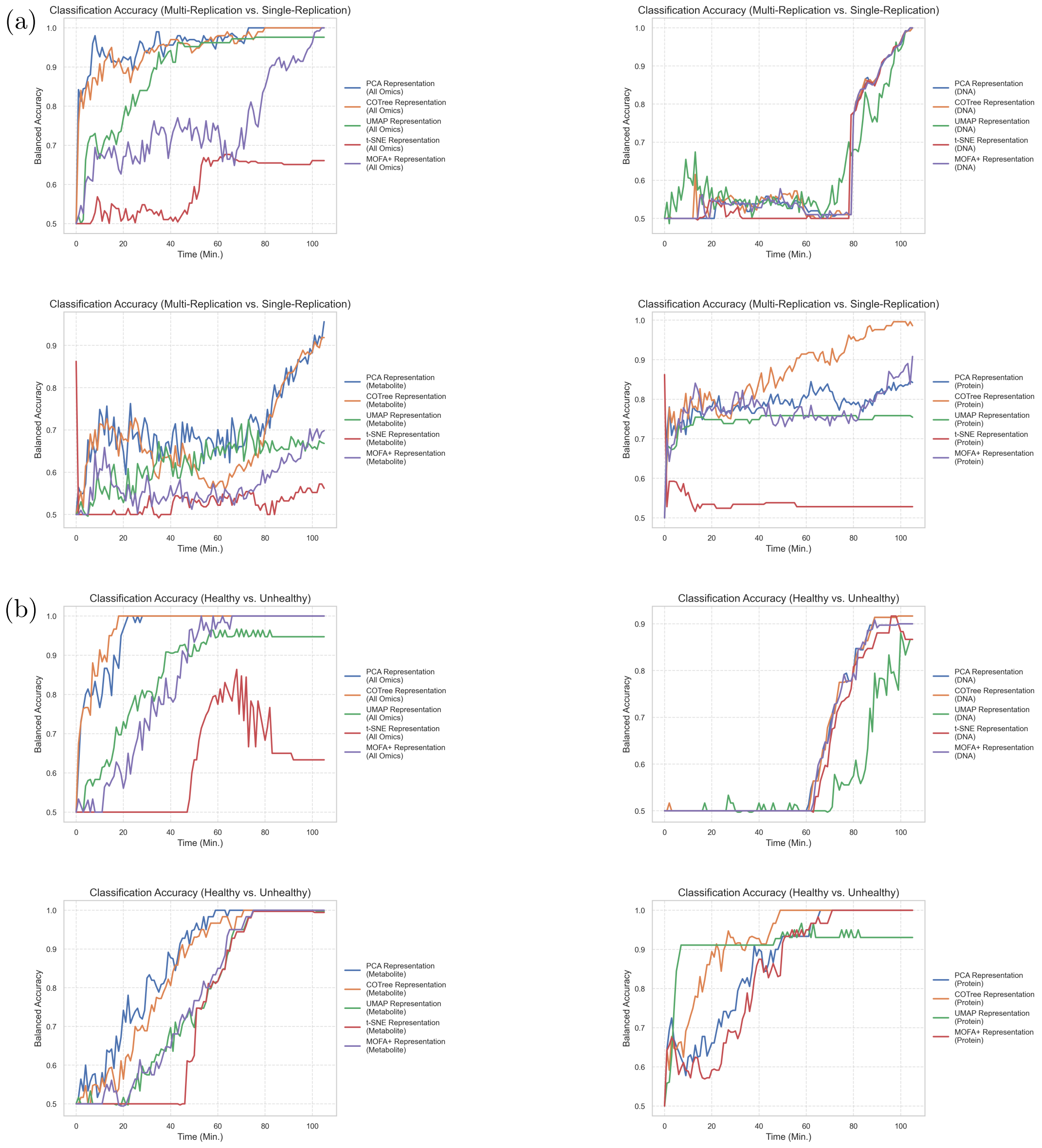


Figure S3: **Balanced accuracy of logistic regression model in classifying cells with different biological behaviors.** A logistic regression model is trained at each time point $t$ to distinguish between two classification tasks: (a) healthy cells undergoing a single DNA replication event versus those with multiple replication events, and (b) healthy versus unhealthy cells. The $y$-axis indicates balanced accuracy, while the $x$-axis represents time. Model performance using representations learned by COTree is compared against PCA, t-SNE, UMAP, and MOFA+ representations.

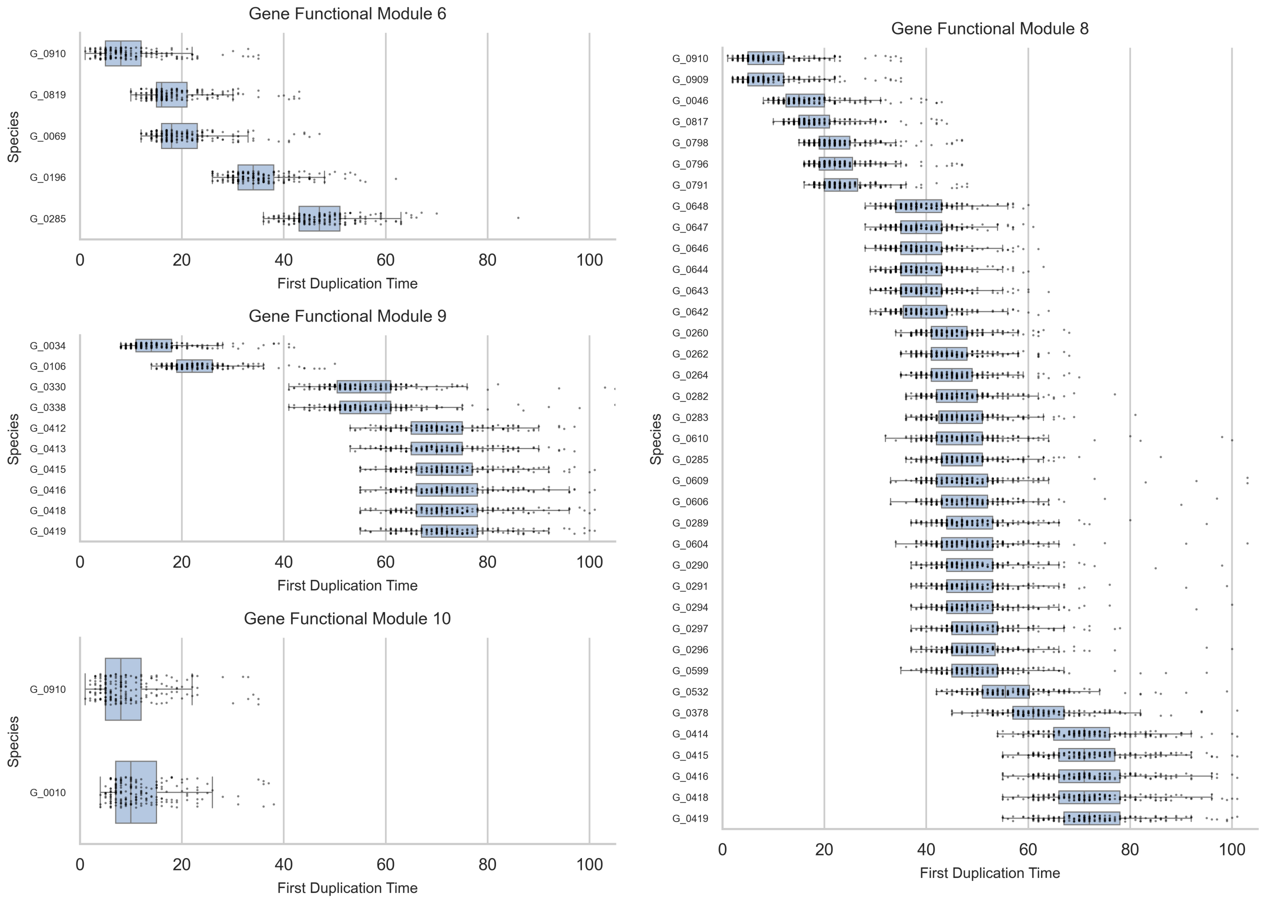


Figure S4: **First duplication time for genes in gene functional modules 6, 8, 9, and 10.** Boxplots present the first duplication time across cells for genes in four selected functional modules.

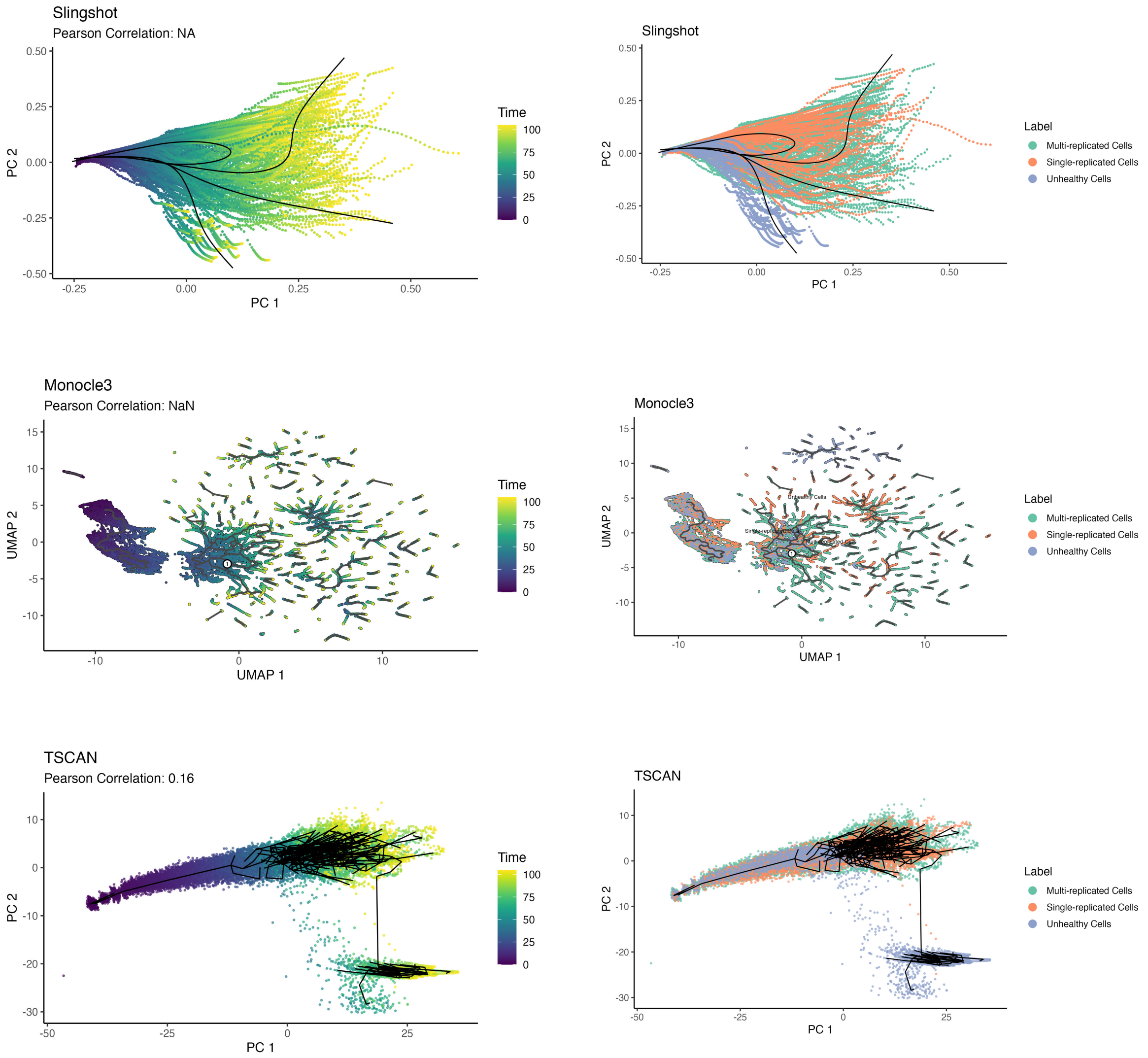


Figure S5: **Trajectory tree constructed using existing methods.** Trajectory tree constructed by Slingshot, Monocle3, and TSCAN with full cell trajectories as input.

### S4 Supplemental Tables

Table S2: Protein functional modules.

| Module | Protein ID | Function |
|---|---|---|
| Module 1 | P_0426 | Phosphate ABC transporter permease |
| | P_0005 | Uncharacterized protein |
| | P_0814 | UDP-galactopyranose mutase |
| | P_0202 | 16S rRNA (guanine(966)-N(2))-methyltransferase |
| | P_0692 | Uncharacterized pseudouridine synthase |
| | P_0726 | Glucosamine-6-phosphate deaminase |
| | P_0441 | Cysteine desulfurase |
| | P_0195 | Spermidine/putrescine ABC transporter permease |
| | P_0523 | Cell division protein |
| | P_0639 | Uncharacterized efflux ABC transporter permease |
| | P_0708 | Thiamine ABC transporter substrate-binding protein |
| | P_0010 | Nucleoside ABC transporter ATP-binding protein |
| | P_0608 | Primosomal protein |
| | P_0526 | 50S ribosomal protein L32 |
| | P_0137 | 50S ribosomal protein L31 |
| | P_0644 | 50S ribosomal protein L17 |
| | P_0819 | Thioredoxin-disulfide reductase |
| | P_0375 | Uncharacterized protein |
| | P_0302 | Flavin reductase |

*continued on next page*

Table S2 – *continued from previous page*

| **Module** | **Protein ID** | **Function** |
|---|---|---|
| | P_0793 | F0F1 ATP synthase subunit delta |
| | P_0524 | 16S rRNA (cytosine(1402)-N(4))-methyltransferase |
| | P_0650 | Type I methionyl aminopeptidase |
| | P_0348 | Ribosome biogenesis GTPase |
| | P_0097 | Flap endonuclease and 5' exonuclease |
| | P_0391 | Elongation factor P |
| | P_0840 | Antitermination protein |
| | P_0039 | Membrane anchored AAA+ protease |
| | P_0645 | DNA-directed RNA polymerase subunit alpha |
| | P_0431 | Uncharacterized metallophosphoesterase |
| | P_0411 | Uncharacterized protein |
| | P_0452 | DNA topoisomerase IV subunit B |
| | P_0113 | Glycolipid synthase A |
| | P_0881 | Uncharacterized MFS transporter |
| | P_0002 | DNA polymerase III subunit beta |
| | P_0691 | Uncharacterized protein |
| | P_0289 | Ribosome-binding factor A |
| | P_0642 | ECF transporter ATPase |
| | P_0338 | Uncharacterized lipoprotein |
| | P_0428 | Phosphate transport system regulatory protein |
| | P_0250 | Uncharacterized protein |
| | P_0299 | Uncharacterized protein |
| | P_0145 | Uncharacterized acetyltransferase |
| | P_0285 | Elongation factor 4 |

*continued on next page*

Table S2 – *continued from previous page*

| Module | Protein ID | Function |
|---|---|---|
| | P_0885 | tRNA uridine(34) 5-carboxymethylaminomethyl synthesis enzyme |
| | P_0045 | dTMP kinase |
| | P_0834 | Replicative DNA helicase |
| | P_0132 | Toxin-antitoxin AAA ATPase |
| | P_0537 | UMP kinase |
| | P_0689 | Glutamyl-tRNA amidotransferase subunit C |
| | P_0330 | Deoxyguanosine kinase |
| | P_0361 | 23S rRNA (pseudouridine(1915)-N3)-methyltransferase |
| | P_0515 | Cytidine deaminase |
| | P_0346 | Uncharacterized protein |
| | P_0605 | Uncharacterized protein |
| | P_0294 | 30S ribosomal protein S15 |
| | P_0614 | Nicotinate phosphoribosyltransferase |
| | P_0832 | Aminoacyl-tRNA hydrolase |
| | P_0445 | Glucose-6-phosphate isomerase |
| Module 2 | P_0666 | 50S ribosomal protein L22 |
| | P_0257 | RNase J family beta-CASP ribonuclease |
| | P_0360 | Signal recognition particle protein |
| | P_0372 | Flippase B |
| | P_0107 | Transcription antitermination factor |
| | P_0030 | Uncharacterized ABC transporter ATP-binding protein |
| | P_0640 | tRNA pseudouridine(38-40) synthase |
| | P_0796 | F0F1 ATP synthase subunit A |

*continued on next page*

Table S2 – *continued from previous page*

| **Module** | **Protein ID** | **Function** |
|---|---|---|
| | P_0617 | Fatty acid binding protein |
| | P_0661 | 50S ribosomal protein L14 |
| | P_0646 | 30S ribosomal protein S11 |
| | P_0814 | UDP-galactopyranose mutase |
| | P_0930 | 50S ribosomal protein L33 |
| | P_0852 | Uncharacterized protein |
| | P_0138 | Uncharacterized protein |
| | P_0524 | 16S rRNA (cytosine(1402)-N(4))-methyltransferase |
| | P_0851 | Uncharacterized lipoprotein |
| | P_0380 | Nicotinate (nicotinamide) nucleotide adenylyltransferase |
| | P_0776 | SsrA-binding protein |
| | P_0657 | 30S ribosomal protein S8 |
| | P_0406 | DNA primase |
| | P_0918 | Imidazoleglycerol-phosphate dehydratase |
| | P_0168 | Oligopeptide ABC transporter ATP-binding protein |
| | P_0353 | Cell division DivIVA/GpsB protein |
| | P_0166 | Oligopeptide ABC transporter permease |
| | P_0040 | tRNA lysidine(34) synthetase |
| | P_0693 | Uncharacterized protease |
| | P_0611 | DNA polymerase I |
| | P_0328 | Chromosome segregation protein B |
| | P_0167 | Oligopeptide ABC transporter ATP-binding protein |

*continued on next page*

Table S2 – *continued from previous page*

| Module | Protein ID | Function |
|---|---|---|
| | P_0375 | Uncharacterized protein |
| | P_0046 | Recombination protein |
| | P_0240 | tRNA 4-thiouridine(8) synthase |
| | P_0285 | Elongation factor 4 |
| | P_0043 | Uncharacterized methyltransferase |
| | P_0906 | Uncharacterized protein |
| | P_0610 | DNA-formamidopyrimidine glycosylase |
| | P_0515 | Cytidine deaminase |
| | P_0437 | Putative 3'-5' exoribonuclease |
| | P_0438 | Uncharacterized protein |
| | P_0250 | Uncharacterized protein |
| | P_0359 | Ribonuclease Y |
| | P_0291 | FAD synthetase |
| | P_0818 | Diacylglyceryl transferase |
| | P_0402 | rRNA maturation RNase |
| | P_0516 | Uncharacterized protein |
| | P_0254 | Excinuclease ABC subunit C |
| | P_0200 | Translation initiation factor IF-3 |
| | P_0137 | 50S ribosomal protein L31 |
| | P_0195 | Spermidine/putrescine ABC transporter permease |
| | P_0428 | Phosphate transport system regulatory protein |
| | P_0872 | Uncharacterized ATPase |
| | P_0876 | Uncharacterized amino acid permease |

*continued on next page*

Table S2 – *continued from previous page*

| Module | Protein ID | Function |
|---|---|---|
| | P_0144 | tRNA (adenosine(37)-N6)-threonylcarbamoyltransferase complex transferase subunit |
| | P_0126 | Glutamate–tRNA ligase |
| | P_0154 | Uncharacterized peptidase |
| | P_0392 | Uncharacterized protein |
| | P_0418 | Ribonuclease III |
| | P_0541 | Molecular chaperone |
| | P_0444 | Oligopeptidase |
| | P_0433 | Uncharacterized protein |
| | P_0525 | Cell division/cell wall cluster transcriptional repressor |
| | P_0452 | DNA topoisomerase IV subunit B |
| | P_0033 | Uncharacterized protein |
| | P_0800 | Ribose 5-phosphate isomerase B |
| | P_0887 | Coenzyme A Disulfide Reductase |
| | P_0026 | Single-stranded DNA-binding protein |
| | P_0081 | tRNA uridine(34) 5-carboxymethylaminomethyl synthesis GTPase |
| | P_0411 | Uncharacterized protein |
| | P_0479 | Uncharacterized peptidase |
| | P_0202 | 16S rRNA (guanine(966)-N(2))-methyltransferase |
| | P_0684 | Bifunctional 5,10-methylene-tetrahydrofolate dehydrogenase/5,10-methylene-tetrahydrofolate cyclohydrolase |

*continued on next page*

Table S2 – *continued from previous page*

| **Module** | **Protein ID** | **Function** |
|---|---|---|
| | P_0045 | dTMP kinase |
| | P_0639 | Uncharacterized efflux ABC transporter permease |
| | P_0214 | Phosphatidylglycerophosphatase |
| | P_0641 | ECF transporter T component |
| | P_0127 | Uncharacterized phosphohydrolase |
| | P_0419 | Phosphate acyltransferase |
| | P_0301 | Ribosome assembly cofactor |
| | P_0687 | Glutamyl-tRNA amidotransferase subunit B |
| | P_0453 | DNA topoisomerase IV subunit A |
| | P_0344 | Inorganic diphosphatase |
| | P_0427 | Phosphate ABC transporter ATP-binding protein |
| | P_0133 | Toxin-antitoxin serine protease |
| | P_0142 | Protein-(glutamine-N5) methyltransferase, release factor-specific |
| | P_0215 | Pre-16S rRNA nuclease |
| | P_0114 | Glycolipid synthase B |
| | P_0840 | Antitermination protein |
| | P_0732 | Deoxyribose-phosphate aldolase |
| | P_0670 | 50S ribosomal protein L4 |
| | P_0707 | Thiamine ABC transporter ATP-binding protein |
| | P_0390 | Methionyl-tRNA formyltransferase |
| | P_0006 | DNA gyrase subunit B |
| Module 3 | P_0147 | Cardiolipin synthase |

*continued on next page*

Table S2 – *continued from previous page*

| **Module** | **Protein ID** | **Function** |
|---|---|---|
| | P_0360 | Signal recognition particle protein |
| | P_0835 | Uncharacterized lipoprotein |
| | P_0375 | Uncharacterized protein |
| | P_0429 | Signal recognition particle-docking protein |
| | P_0876 | Uncharacterized amino acid permease |
| | P_0026 | Single-stranded DNA-binding protein |
| | P_0029 | FMN reductase |
| | P_0148 | 30S ribosomal protein S12 |
| | P_0066 | dUMP phosphatase |
| | P_0435 | Mannose-6-phosphate isomerase |
| | P_0127 | Uncharacterized phosphohydrolase |
| | P_0505 | Uncharacterized lipoprotein |
| | P_0771 | Ribonucleotide-diphosphate reductase subunit alpha |
| | P_0146 | Uncharacterized protein |
| | P_0289 | Ribosome-binding factor A |
| | P_0652 | Preprotein translocase subunit |
| | P_0831 | Phosphoribosylpyrophosphate synthetase |
| | P_0879 | magnesium-importing ATPase |
| | P_0144 | tRNA (adenosine(37)-N6)-threonylcarbamoyltransferase complex transferase subunit |
| | P_0668 | 50S ribosomal protein L2 |
| | P_0257 | RNase J family beta-CASP ribonuclease |
| | P_0642 | ECF transporter ATPase |

*continued on next page*

Table S2 – *continued from previous page*

| Module | Protein ID | Function |
|---|---|---|
| | P_0366 | L16-binding dependent 50S subunit-maturation GTPase |
| | P_0908 | Membrane protein insertase |
| | P_0428 | Phosphate transport system regulatory protein |
| | P_0886 | Proton-glutamate symporter |
| | P_0636 | Uncharacterized lipoprotein |
| | P_0859 | DNA topoisomerase I |
| | P_0218 | Glycerol kinase |
| | P_0881 | Uncharacterized MFS transporter |
| | P_0108 | Uncharacterized lipoprotein |
| | P_0353 | Cell division DivIVA/GpsB protein |
| | P_0887 | Coenzyme A Disulfide Reductase |
| | P_0639 | Uncharacterized efflux ABC transporter permease |
| | P_0228 | Dihydrolipoyl dehydrogenase |
| | P_0116 | Uncharacterized protein |
| | P_0438 | Uncharacterized protein |
| | P_0814 | UDP-galactopyranose mutase |
| | P_0789 | F0F1 ATP synthase subunit delta/epsilon |
| | P_0651 | Adenylate kinase |
| | P_0729 | Phosphoglycerate mutase (2,3-diphosphoglycerate-independent) |
| | P_0168 | Oligopeptide ABC transporter ATP-binding protein |
| | P_0412 | Uncharacterized protein |
| | P_0330 | Deoxyguanosine kinase |

*continued on next page*

Table S2 – *continued from previous page*

| Module | Protein ID | Function |
| --- | --- | --- |
| | P_0549 | Non-canonical purine NTP pyrophosphatase |
| | P_0610 | DNA-formamidopyrimidine glycosylase |
| | P_0453 | DNA topoisomerase IV subunit A |
| | P_0689 | Glutamyl-tRNA amidotransferase subunit C |
| | P_0821 | HPr(Ser) kinase/phosphatase |
| | P_0431 | Uncharacterized metallophosphoesterase |
| | P_0391 | Elongation factor P |
| | P_0167 | Oligopeptide ABC transporter ATP-binding protein |
| | P_0166 | Oligopeptide ABC transporter permease |
| | P_0838 | 23S rRNA (guanosine(2251)-2'-O)-methyltransferase |
| | P_0592 | Uncharacterized protein |
| | P_0733 | Phosphopentomutase |
| | P_0362 | 30S ribosomal protein S16 |
| | P_0614 | Nicotinate phosphoribosyltransferase |
| | P_0710 | Low specificity hydrolase |
| | P_0328 | Chromosome segregation protein B |
| | P_0686 | Potassium transporter |
| | P_0427 | Phosphate ABC transporter ATP-binding protein |
| | P_0140 | Thymidine kinase |
| | P_0605 | Uncharacterized protein |
| | P_0250 | Uncharacterized protein |
| Module 4 | P_0229 | Phosphate acetyltransferase |
| | P_0006 | DNA gyrase subunit B |

*continued on next page*

Table S2 – *continued from previous page*

| Module | Protein ID | Function |
|---|---|---|
| | P_0410 | Degradosome RNA helicase |
| | P_0138 | Uncharacterized protein |
| | P_0361 | 23S rRNA (pseudouridine(1915)-N3)-methyltransferase |
| | P_0366 | L16-binding dependent 50S subunit-maturation GTPase |
| | P_0733 | Phosphopentomutase |
| | P_0403 | Ribosome GTPase |
| | P_0879 | magnesium-importing ATPase |
| | P_0728 | Low specificity hydrolase |
| | P_0651 | Adenylate kinase |
| | P_0380 | Nicotinate (nicotinamide) nucleotide adenylyl-transferase |
| | P_0511 | Uncharacterized protein |
| | P_0794 | F0F1 ATP synthase subunit B |
| | P_0429 | Signal recognition particle-docking protein |
| | P_0301 | Ribosome assembly cofactor |
| | P_0303 | PolC-type DNA polymerase III |
| | P_0645 | DNA-directed RNA polymerase subunit alpha |
| | P_0126 | Glutamate–tRNA ligase |
| | P_0872 | Uncharacterized ATPase |
| | P_0108 | Uncharacterized lipoprotein |
| | P_0691 | Uncharacterized protein |
| | P_0834 | Replicative DNA helicase |
| | P_0779 | PTS sugar transporter |
| | P_0165 | Oligopeptide ABC transporter permease |

*continued on next page*

Table S2 – *continued from previous page*

| Module | Protein ID | Function |
|---|---|---|
| | P_0601 | Uncharacterized protein |
| | P_0378 | NAD(+) synthase |
| | P_0697 | Uncharacterized glycosyl transferase |
| | P_0005 | Uncharacterized protein |
| | P_0391 | Elongation factor P |
| | P_0859 | DNA topoisomerase I |
| | P_0042 | Uncharacterized transcriptional regulator |
| | P_0044 | DNA polymerase III subunit delta |
| | P_0512 | Acyl-phosphate glycerol 3-phosphate acyltransferase |
| | P_0302 | Flavin reductase |
| | P_0832 | Aminoacyl-tRNA hydrolase |
| | P_0548 | tRNA (cytidine(34)-2'-O)-methyltransferase |
| | P_0202 | 16S rRNA (guanine(966)-N(2))-methyltransferase |
| | P_0218 | Glycerol kinase |
| | P_0419 | Phosphate acyltransferase |
| | P_0887 | Coenzyme A Disulfide Reductase |
| | P_0726 | Glucosamine-6-phosphate deaminase |
| | P_0729 | Phosphoglycerate mutase (2,3-diphosphoglycerate-independent) |
| | P_0885 | tRNA uridine(34) 5-carboxymethylaminomethyl synthesis enzyme |
| | P_0200 | Translation initiation factor IF-3 |
| | P_0881 | Uncharacterized MFS transporter |
| | P_0359 | Ribonuclease Y |

*continued on next page*

Table S2 – *continued from previous page*

| **Module** | **Protein ID** | **Function** |
|---|---|---|
| | P_0517 | Uncharacterized RNA pseudouridine synthase |
| | P_0250 | Uncharacterized protein |
| | P_0611 | DNA polymerase I |
| | P_0799 | Serine hydroxymethyltransferase |
| Module 5 | P_0479 | Uncharacterized peptidase |
| | P_0516 | Uncharacterized protein |
| | P_0407 | RNA polymerase subunit sigma |
| | P_0430 | Uncharacterized DNA-binding protein |
| | P_0114 | Glycolipid synthase B |
| | P_0792 | F0F1 ATP synthase subunit alpha |
| | P_0451 | Glyceraldehyde-3-phosphate dehydrogenase |
| | P_0168 | Oligopeptide ABC transporter ATP-binding protein |
| | P_0299 | Uncharacterized protein |
| | P_0378 | NAD(+) synthase |
| | P_0291 | FAD synthetase |
| | P_0389 | Uncharacterized protein |
| | P_0007 | DNA gyrase subunit A |
| | P_0108 | Uncharacterized lipoprotein |
| | P_0870 | Uncharacterized C4-dicarboxylate ABC transporter |
| | P_0608 | Primosomal protein |
| | P_0694 | ptsH: Phosphocarrier protein |
| | P_0592 | Uncharacterized protein |
| | P_0652 | Preprotein translocase subunit |
| | P_0687 | Glutamyl-tRNA amidotransferase subunit B |

*continued on next page*

Table S2 – *continued from previous page*

| Module | Protein ID | Function |
|---|---|---|
| | P_0435 | Mannose-6-phosphate isomerase |
| | P_0655 | 50S ribosomal protein L18 |
| | P_0141 | Peptide chain release factor 1 |
| | P_0139 | NanoRNAse |
| | P_0148 | 30S ribosomal protein S12 |
| | P_0887 | Coenzyme A Disulfide Reductase |
| | P_0773 | Ribonucleoside-diphosphate reductase subunit beta |
| | P_0538 | Uncharacterized protein |
| | P_0330 | Deoxyguanosine kinase |
| | P_0614 | Nicotinate phosphoribosyltransferase |
| | P_0359 | Ribonuclease Y |
| | P_0434 | 23S rRNA (uridine(1939)-m5)-methyltransferase |
| | P_0411 | Uncharacterized protein |
| | P_0366 | L16-binding dependent 50S subunit-maturation GTPase |
| | P_0040 | tRNA lysidine(34) synthetase |
| | P_0165 | Oligopeptide ABC transporter permease |
| | P_0045 | dTMP kinase |
| | P_0544 | Heat-inducible transcription repressor |
| | P_0605 | Uncharacterized protein |
| | P_0439 | Uncharacterized lipoprotein |
| | P_0199 | 50S ribosomal protein L35 |
| | P_0611 | DNA polymerase I |

*continued on next page*

Table S2 – *continued from previous page*

| **Module** | **Protein ID** | **Function** |
|---|---|---|
| | P_0707 | Thiamine ABC transporter ATP-binding protein |
| | P_0441 | Cysteine desulfurase |
| | P_0772 | Ribonucleotide reductase assembly protein |
| | P_0127 | Uncharacterized phosphohydrolase |
| | P_0200 | Translation initiation factor IF-3 |
| | P_0549 | Non-canonical purine NTP pyrophosphatase |
| | P_0651 | Adenylate kinase |
| | P_0238 | 30S ribosomal protein S4 |
| | P_0872 | Uncharacterized ATPase |
| | P_0444 | Oligopeptidase |
| | P_0202 | 16S rRNA (guanine(966)-N(2))-methyltransferase |
| | P_0885 | tRNA uridine(34) 5-carboxymethylaminomethyl synthesis enzyme |
| | P_0668 | 50S ribosomal protein L2 |
| | P_0732 | Deoxyribose-phosphate aldolase |
| | P_0642 | ECF transporter ATPase |
| | P_0771 | Ribonucleotide-diphosphate reductase subunit alpha |
| | P_0249 | Uncharacterized protein |
| | P_0203 | Guanylate kinase |
| | P_0727 | Triose-phosphate isomerase |
| | P_0138 | Uncharacterized protein |
| | P_0697 | Uncharacterized glycosyl transferase |
| | P_0371 | Flippase A |

*continued on next page*

Table S2 – *continued from previous page*

| Module | Protein ID | Function |
|---|---|---|
| | P_0042 | Uncharacterized transcriptional regulator |
| | P_0289 | Ribosome-binding factor A |
| | P_0418 | Ribonuclease III |
| | P_0195 | Spermidine/putrescine ABC transporter permease |
| | P_0838 | 23S rRNA (guanosine(2251)-2'-O)-methyltransferase |
| | P_0821 | HPr(Ser) kinase/phosphatase |
| | P_0500 | Maturation protease for ribosomal protein L27 |
| Module 6 | P_0143 | Uncharacterized protein |
| | P_0445 | Glucose-6-phosphate isomerase |
| | P_0771 | Ribonucleotide-diphosphate reductase subunit alpha |
| | P_0107 | Transcription antitermination factor |
| | P_0601 | Uncharacterized protein |
| | P_0615 | Uncharacterized protein |
| | P_0046 | Recombination protein |
| | P_0080 | Uncharacterized protein |
| | P_0697 | Uncharacterized glycosyl transferase |
| | P_0108 | Uncharacterized lipoprotein |
| | P_0729 | Phosphoglycerate mutase (2,3-diphosphoglycerate-independent) |
| | P_0169 | Oligopeptide ABC transporter substrate-binding protein |
| | P_0790 | F0F1 ATP synthase subunit beta |
| | P_0138 | Uncharacterized protein |

*continued on next page*

Table S2 – *continued from previous page*

| Module | Protein ID | Function |
|---|---|---|
| | P_0500 | Maturation protease for ribosomal protein L27 |
| | P_0060 | Uncharacterized protein |
| | P_0326 | Uncharacterized protein |
| | P_0165 | Oligopeptide ABC transporter permease |
| | P_0387 | tRNA 2-thiouridine(34) synthase |
| | P_0639 | Uncharacterized efflux ABC transporter permease |
| | P_0851 | Uncharacterized lipoprotein |
| | P_0526 | 50S ribosomal protein L32 |
| | P_0859 | DNA topoisomerase I |
| | P_0610 | DNA-formamidopyrimidine glycosylase |
| | P_0353 | Cell division DivIVA/GpsB protein |
| | P_0419 | Phosphate acyltransferase |
| | P_0007 | DNA gyrase subunit A |
| | P_0906 | Uncharacterized protein |
| | P_0726 | Glucosamine-6-phosphate deaminase |
| | P_0616 | Fatty acid binding protein |
| | P_0794 | F0F1 ATP synthase subunit B |
| | P_0239 | Uncharacterized protein |
| | P_0411 | Uncharacterized protein |
| | P_0366 | L16-binding dependent 50S subunit-maturation GTPase |
| | P_0129 | CTP synthase |
| | P_0392 | Uncharacterized protein |
| | P_0042 | Uncharacterized transcriptional regulator |
| | P_0834 | Replicative DNA helicase |

*continued on next page*

Table S2 – *continued from previous page*

| **Module** | **Protein ID** | **Function** |
|---|---|---|
| | P_0202 | 16S rRNA (guanine(966)-N(2))-methyltransferase |
| | P_0773 | Ribonucleoside-diphosphate reductase subunit beta |
| | P_0294 | 30S ribosomal protein S15 |
| | P_0197 | Spermidine/putrescine ABC transporter ATP-binding protein |
| | P_0420 | Fatty acid kinase subunit A |
| | P_0316 | Transketolase |
| | P_0824 | Excinuclease ABC subunit A |
| | P_0097 | Flap endonuclease and 5' exonuclease |
| | P_0285 | Elongation factor 4 |
| | P_0229 | Phosphate acetyltransferase |
| | P_0201 | Peptide deformylase |
| | P_0840 | Antitermination protein |
| | P_0537 | UMP kinase |
| | P_0872 | Uncharacterized ATPase |
| | P_0530 | Uncharacterized protein |
| | P_0819 | Thioredoxin-disulfide reductase |
| | P_0505 | Uncharacterized lipoprotein |
| | P_0733 | Phosphopentomutase |
| | P_0139 | NanoRNAse |
| | P_0415 | Chromosome segregation protein |
| | P_0512 | Acyl-phosphate glycerol 3-phosphate acyltransferase |
| | P_0549 | Non-canonical purine NTP pyrophosphatase |

*continued on next page*

Table S2 – *continued from previous page*

| Module | Protein ID | Function |
|---|---|---|
| | P_0200 | Translation initiation factor IF-3 |
| | P_0689 | Glutamyl-tRNA amidotransferase subunit C |
| | P_0651 | Adenylate kinase |
| | P_0441 | Cysteine desulfurase |
| | P_0727 | Triose-phosphate isomerase |
| | P_0382 | Deoxynucleoside kinase |
| | P_0422 | 50S ribosomal protein L28 |
| | P_0026 | Single-stranded DNA-binding protein |
| | P_0444 | Oligopeptidase |
| | P_0410 | Degradosome RNA helicase |
| | P_0427 | Phosphate ABC transporter ATP-binding protein |
| | P_0240 | tRNA 4-thiouridine(8) synthase |
| | P_0887 | Coenzyme A Disulfide Reductase |
| | P_0609 | Chromosome replication initiation protein |
| | P_0203 | Guanylate kinase |
| | P_0063 | Uncharacterized tRNA dihydrouridine synthase |
| Module 7 | P_0254 | Excinuclease ABC subunit C |
| | P_0047 | DNA polymerase III subunit gamma and tau |
| | P_0605 | Uncharacterized protein |
| | P_0113 | Glycolipid synthase A |
| | P_0887 | Coenzyme A Disulfide Reductase |
| | P_0645 | DNA-directed RNA polymerase subunit alpha |
| | P_0203 | Guanylate kinase |
| | P_0249 | Uncharacterized protein |
| | P_0733 | Phosphopentomutase |

*continued on next page*

Table S2 – *continued from previous page*

| Module | Protein ID | Function |
|---|---|---|
| | P_0512 | Acyl-phosphate glycerol 3-phosphate acyltransferase |
| | P_0794 | F0F1 ATP synthase subunit B |
| | P_0271 | tRNA N6-adenosine(37)-N6-threonylcarbamoyltransferase complex dimerization subunit |
| | P_0302 | Flavin reductase |
| | P_0908 | Membrane protein insertase |
| | P_0006 | DNA gyrase subunit B |
| | P_0197 | Spermidine/putrescine ABC transporter ATP-binding protein |
| | P_0453 | DNA topoisomerase IV subunit A |
| | P_0859 | DNA topoisomerase I |
| | P_0872 | Uncharacterized ATPase |
| | P_0832 | Aminoacyl-tRNA hydrolase |
| | P_0042 | Uncharacterized transcriptional regulator |
| | P_0129 | CTP synthase |
| | P_0600 | Ribonuclease J |
| | P_0791 | F0F1 ATP synthase subunit gamma |
| | P_0138 | Uncharacterized protein |
| | P_0617 | Fatty acid binding protein |
| | P_0344 | Inorganic diphosphatase |
| | P_0407 | RNA polymerase subunit sigma |
| | P_0299 | Uncharacterized protein |
| | P_0010 | Nucleoside ABC transporter ATP-binding protein |

*continued on next page*

Table S2 – *continued from previous page*

| Module | Protein ID | Function |
|---|---|---|
| | P_0799 | Serine hydroxymethyltransferase |
| | P_0116 | Uncharacterized protein |
| | P_0592 | Uncharacterized protein |
| | P_0009 | Nucleoside ABC transporter permease |
| | P_0377 | Ribosome GTPase |
| | P_0330 | Deoxyguanosine kinase |
| | P_0874 | 16S rRNA (guanine(527)-N(7))-methyltransferase |
| | P_0614 | Nicotinate phosphoribosyltransferase |
| | P_0438 | Uncharacterized protein |
| | P_0228 | Dihydrolipoyl dehydrogenase |
| | P_0823 | Dihydrofolate synthase |
| | P_0779 | PTS sugar transporter |
| | P_0132 | Toxin-antitoxin AAA ATPase |
| | P_0371 | Flippase A |
| | P_0824 | Excinuclease ABC subunit A |
| | P_0500 | Maturation protease for ribosomal protein L27 |
| | P_0201 | Peptide deformylase |
| | P_0834 | Replicative DNA helicase |
| | P_0819 | Thioredoxin-disulfide reductase |
| | P_0081 | tRNA uridine(34) 5-carboxymethylaminomethyl synthesis GTPase |
| | P_0688 | Glutamyl-tRNA amidotransferase subunit A |
| | P_0695 | ATP-dependent DNA helicase |
| | P_0548 | tRNA (cytidine(34)-2'-O)-methyltransferase |

*continued on next page*

Table S2 – *continued from previous page*

| **Module** | **Protein ID** | **Function** |
|---|---|---|
| | P_0427 | Phosphate ABC transporter ATP-binding protein |
| | P_0039 | Membrane anchored AAA+ protease |
| | P_0285 | Elongation factor 4 |
| | P_0689 | Glutamyl-tRNA amidotransferase subunit C |
| Module 8 | P_0440 | Uncharacterized lipoprotein |
| | P_0115 | UTP–glucose-1-phosphate uridylyltransferase |
| | P_0154 | Uncharacterized peptidase |
| | P_0030 | Uncharacterized ABC transporter ATP-binding protein |
| | P_0126 | Glutamate–tRNA ligase |
| | P_0168 | Oligopeptide ABC transporter ATP-binding protein |
| | P_0730 | Uncharacterized protein |
| | P_0544 | Heat-inducible transcription repressor |
| | P_0642 | ECF transporter ATPase |
| | P_0415 | Chromosome segregation protein |
| | P_0203 | Guanylate kinase |
| | P_0434 | 23S rRNA (uridine(1939)-m5)-methyltransferase |
| | P_0600 | Ribonuclease J |
| | P_0690 | DNA ligase (NAD(+)) |
| | P_0113 | Glycolipid synthase A |
| | P_0708 | Thiamine ABC transporter substrate-binding protein |
| | P_0789 | F0F1 ATP synthase subunit delta/epsilon |

*continued on next page*

Table S2 – *continued from previous page*

| Module | Protein ID | Function |
|---|---|---|
| | P_0840 | Antitermination protein |
| | P_0239 | Uncharacterized protein |
| | P_0435 | Mannose-6-phosphate isomerase |
| | P_0373 | Uncharacterized protein |
| | P_0729 | Phosphoglycerate mutase (2,3-diphosphoglycerate-independent) |
| | P_0645 | DNA-directed RNA polymerase subunit alpha |
| | P_0686 | Potassium transporter |
| | P_0221 | Pyruvate kinase |
| | P_0143 | Uncharacterized protein |
| | P_0524 | 16S rRNA (cytosine(1402)-N(4))-methyltransferase |
| | P_0610 | DNA-formamidopyrimidine glycosylase |
| | P_0346 | Uncharacterized protein |
| | P_0400 | Uncharacterized protease |
| | P_0361 | 23S rRNA (pseudouridine(1915)-N3)-methyltransferase |
| | P_0451 | Glyceraldehyde-3-phosphate dehydrogenase |
| | P_0238 | 30S ribosomal protein S4 |
| | P_0511 | Uncharacterized protein |
| | P_0146 | Uncharacterized protein |
| | P_0364 | tRNA (guanosine(37)-N1)-methyltransferase |
| | P_0289 | Ribosome-binding factor A |
| | P_0541 | Molecular chaperone |
| | P_0445 | Glucose-6-phosphate isomerase |
| | P_0517 | Uncharacterized RNA pseudouridine synthase |

*continued on next page*

Table S2 – *continued from previous page*

| **Module** | **Protein ID** | **Function** |
|---|---|---|
| | P_0412 | Uncharacterized protein |
| | P_0065 | Thioredoxin |
| | P_0636 | Uncharacterized lipoprotein |
| | P_0870 | Uncharacterized C4-dicarboxylate ABC transporter |
| | P_0516 | Uncharacterized protein |
| | P_0148 | 30S ribosomal protein S12 |
| | P_0641 | ECF transporter T component |
| | P_0366 | L16-binding dependent 50S subunit-maturation GTPase |
| | P_0834 | Replicative DNA helicase |
| | P_0330 | Deoxyguanosine kinase |
| | P_0710 | Low specificity hydrolase |
| | P_0831 | Phosphoribosylpyrophosphate synthetase |
| | P_0294 | 30S ribosomal protein S15 |
| | P_0229 | Phosphate acetyltransferase |
| | P_0835 | Uncharacterized lipoprotein |
| | P_0145 | Uncharacterized acetyltransferase |
| | P_0504 | 16S rRNA (cytidine(1402)-2'-O)-methyltransferase |
| | P_0303 | PolC-type DNA polymerase III |
| | P_0697 | Uncharacterized glycosyl transferase |
| | P_0428 | Phosphate transport system regulatory protein |
| | P_0523 | Cell division protein |
| | P_0733 | Phosphopentomutase |
| | P_0791 | F0F1 ATP synthase subunit gamma |

*continued on next page*

Table S2 – *continued from previous page*

| **Module** | **Protein ID** | **Function** |
|---|---|---|
| | P_0872 | Uncharacterized ATPase |
| | P_0503 | Uncharacterized protein |
| | P_0549 | Non-canonical purine NTP pyrophosphatase |
| | P_0140 | Thymidine kinase |
| | P_0427 | Phosphate ABC transporter ATP-binding protein |
| | P_0688 | Glutamyl-tRNA amidotransferase subunit A |
| | P_0650 | Type I methionyl aminopeptidase |
| | P_0732 | Deoxyribose-phosphate aldolase |
| | P_0526 | 50S ribosomal protein L32 |
| | P_0291 | FAD synthetase |
| | P_0906 | Uncharacterized protein |
| | P_0494 | N-acetylmannosamine-6-phosphate 2-epimerase |
| | P_0301 | Ribosome assembly cofactor |
| | P_0001 | Chromosomal replication initiator protein |
| | P_0299 | Uncharacterized protein |
| | P_0372 | Flippase B |
| | P_0391 | Elongation factor P |
| | P_0874 | 16S rRNA (guanine(527)-N(7))-methyltransferase |
| | P_0002 | DNA polymerase III subunit beta |
| | P_0479 | Uncharacterized peptidase |
| | P_0838 | 23S rRNA (guanosine(2251)-2'-O)-methyltransferase |
| | P_0859 | DNA topoisomerase I |

*continued on next page*

Table S2 – *continued from previous page*

| Module | Protein ID | Function |
|---|---|---|
| | P_0614 | Nicotinate phosphoribosyltransferase |
| | P_0805 | Uncharacterized protein |
| | P_0080 | Uncharacterized protein |
| | P_0649 | Translation initiation factor IF-1 |
| | P_0344 | Inorganic diphosphatase |
| | P_0684 | Bifunctional 5,10-methylene-tetrahydrofolate dehydrogenase/5,10-methylene-tetrahydrofolate cyclohydrolase |
| | P_0257 | RNase J family beta-CASP ribonuclease |
| | P_0350 | DNA-binding protein |
| | P_0144 | tRNA (adenosine(37)-N6)-threonylcarbamoyltransferase complex transferase subunit |
| | P_0247 | Ribosome biogenesis GTP-binding protein |
| | P_0377 | Ribosome GTPase |
| | P_0371 | Flippase A |
| | P_0444 | Oligopeptidase |
| | P_0794 | F0F1 ATP synthase subunit B |
| Module 9 | P_0779 | PTS sugar transporter |
| | P_0500 | Maturation protease for ribosomal protein L27 |
| | P_0652 | Preprotein translocase subunit |
| | P_0494 | N-acetylmannosamine-6-phosphate 2-epimerase |
| | P_0353 | Cell division DivIVA/GpsB protein |
| | P_0365 | 50S ribosomal protein L19 |
| | P_0304 | Phosphatidate cytidylyltransferase |

*continued on next page*

Table S2 – *continued from previous page*

| Module | Protein ID | Function |
|---|---|---|
| | P_0234 | PTS glucose transporter subunit IIA |
| | P_0141 | Peptide chain release factor 1 |
| | P_0220 | 6-Phosphofructokinase |
| | P_0421 | Uncharacterized protein |
| | P_0840 | Antitermination protein |
| | P_0094 | Uncharacterized protein |
| | P_0199 | 50S ribosomal protein L35 |
| | P_0154 | Uncharacterized peptidase |
| | P_0773 | Ribonucleoside-diphosphate reductase subunit beta |
| | P_0482 | 30S ribosomal protein S21 |
| | P_0046 | Recombination protein |
| | P_0695 | ATP-dependent DNA helicase |
| | P_0689 | Glutamyl-tRNA amidotransferase subunit C |
| | P_0044 | DNA polymerase III subunit delta |
| | P_0544 | Heat-inducible transcription repressor |
| | P_0692 | Uncharacterized pseudouridine synthase |
| | P_0301 | Ribosome assembly cofactor |
| | P_0011 | Nucleoside ABC transporter substrate-binding protein |
| | P_0428 | Phosphate transport system regulatory protein |
| | P_0133 | Toxin-antitoxin serine protease |
| | P_0040 | tRNA lysidine(34) synthetase |
| | P_0794 | F0F1 ATP synthase subunit B |
| | P_0283 | Double-stranded RNA binding RNase HI |
| | P_0329 | Uncharacterized pseudouridine synthase |

*continued on next page*

Table S2 – *continued from previous page*

| **Module** | **Protein ID** | **Function** |
|---|---|---|
| | P_0549 | Non-canonical purine NTP pyrophosphatase |
| | P_0875 | CDP-diacylglycerol–glycerol-3-phosphate 3-phosphatidyltransferase |
| | P_0129 | CTP synthase |
| | P_0515 | Cytidine deaminase |
| | P_0109 | Deoxyribonuclease IV |
| | P_0361 | 23S rRNA (pseudouridine(1915)-N3)-methyltransferase |
| | P_0439 | Uncharacterized lipoprotein |
| | P_0448 | Uncharacterized rRNA methyltransferase |
| | P_0375 | Uncharacterized protein |
| | P_0114 | Glycolipid synthase B |
| | P_0415 | Chromosome segregation protein |
| | P_0651 | Adenylate kinase |
| | P_0526 | 50S ribosomal protein L32 |
| | P_0010 | Nucleoside ABC transporter ATP-binding protein |
| | P_0143 | Uncharacterized protein |
| | P_0645 | DNA-directed RNA polymerase subunit alpha |
| | P_0254 | Excinuclease ABC subunit C |
| | P_0257 | RNase J family beta-CASP ribonuclease |
| | P_0517 | Uncharacterized RNA pseudouridine synthase |
| | P_0314 | Uncharacterized ECF transporter S component |
| | P_0688 | Glutamyl-tRNA amidotransferase subunit A |
| | P_0429 | Signal recognition particle-docking protein |
| | P_0200 | Translation initiation factor IF-3 |

*continued on next page*

Table S2 – *continued from previous page*

| Module | Protein ID | Function |
|---|---|---|
| | P_0043 | Uncharacterized methyltransferase |
| | P_0390 | Methionyl-tRNA formyltransferase |
| | P_0592 | Uncharacterized protein |
| | P_0348 | Ribosome biogenesis GTPase |
| | P_0504 | 16S rRNA (cytidine(1402)-2'-O)-methyltransferase |
| | P_0081 | tRNA uridine(34) 5-carboxymethylaminomethyl synthesis GTPase |
| | P_0146 | Uncharacterized protein |
| | P_0344 | Inorganic diphosphatase |
| | P_0377 | Ribosome GTPase |
| | P_0838 | 23S rRNA (guanosine(2251)-2'-O)-methyltransferase |
| | P_0874 | 16S rRNA (guanine(527)-N(7))-methyltransferase |
| | P_0168 | Oligopeptide ABC transporter ATP-binding protein |
| | P_0388 | Uncharacterized protein |
| | P_0116 | Uncharacterized protein |
| | P_0346 | Uncharacterized protein |
| | P_0371 | Flippase A |
| | P_0726 | Glucosamine-6-phosphate deaminase |
| | P_0732 | Deoxyribose-phosphate aldolase |
| Module 10 | P_0932 | 50S ribosomal protein L33 |

*continued on next page*

Table S2 – *continued from previous page*

| Module | Protein ID | Function |
|---|---|---|
| | P_0271 | tRNA N6-adenosine(37)-N6-threonylcarbamoyltransferase complex dimerization subunit |
| | P_0687 | Glutamyl-tRNA amidotransferase subunit B |
| | P_0879 | magnesium-importing ATPase |
| | P_0733 | Phosphopentomutase |
| | P_0108 | Uncharacterized lipoprotein |
| | P_0117 | Acyl-phosphate glycerol 3-phosphate acyltransferase |
| | P_0440 | Uncharacterized lipoprotein |
| | P_0608 | Primosomal protein |
| | P_0645 | DNA-directed RNA polymerase subunit alpha |
| | P_0823 | Dihydrofolate synthase |
| | P_0728 | Low specificity hydrolase |
| | P_0438 | Uncharacterized protein |
| | P_0113 | Glycolipid synthase A |
| | P_0499 | 50S ribosomal protein L27 |
| | P_0300 | Transcription termination/antitermination protein NusA |
| | P_0203 | Guanylate kinase |
| | P_0851 | Uncharacterized lipoprotein |
| | P_0257 | RNase J family beta-CASP ribonuclease |
| | P_0479 | Uncharacterized peptidase |
| | P_0079 | tRNA (adenosine(37)-N6)-threonylcarbamoyltransferase complex transferase subunit D |

*continued on next page*

Table S2 – *continued from previous page*

| Module | Protein ID | Function |
|---|---|---|
| | P_0042 | Uncharacterized transcriptional regulator |
| | P_0249 | Uncharacterized protein |
| | P_0240 | tRNA 4-thiouridine(8) synthase |
| | P_0200 | Translation initiation factor IF-3 |
| | P_0371 | Flippase A |
| | P_0301 | Ribosome assembly cofactor |
| | P_0908 | Membrane protein insertase |
| | P_0406 | DNA primase |
| | P_0411 | Uncharacterized protein |
| | P_0316 | Transketolase |
| | P_0006 | DNA gyrase subunit B |
| | P_0707 | Thiamine ABC transporter ATP-binding protein |
| | P_0202 | 16S rRNA (guanine(966)-N(2))-methyltransferase |
| | P_0444 | Oligopeptidase |
| | P_0039 | Membrane anchored AAA+ protease |
| | P_0697 | Uncharacterized glycosyl transferase |
| | P_0054 | Uncharacterized peroxiredoxin |
| | P_0796 | F0F1 ATP synthase subunit A |
| | P_0081 | tRNA uridine(34) 5-carboxymethylaminomethyl synthesis GTPase |
| | P_0195 | Spermidine/putrescine ABC transporter permease |
| | P_0453 | DNA topoisomerase IV subunit A |
| | P_0452 | DNA topoisomerase IV subunit B |

*continued on next page*

Table S2 – *continued from previous page*

| **Module** | **Protein ID** | **Function** |
|---|---|---|
| | P_0543 | Nucleotide exchange factor |
| | P_0168 | Oligopeptide ABC transporter ATP-binding protein |
| | P_0228 | Dihydrolipoyl dehydrogenase |
| | P_0610 | DNA-formamidopyrimidine glycosylase |
| | P_0512 | Acyl-phosphate glycerol 3-phosphate acyltransferase |
| | P_0445 | Glucose-6-phosphate isomerase |
| | P_0289 | Ribosome-binding factor A |
| | P_0772 | Ribonucleotide reductase assembly protein |
| Module 11 | P_0151 | Translation elongation factor Tu |
| | P_0380 | Nicotinate (nicotinamide) nucleotide adenylyltransferase |
| | P_0378 | NAD(+) synthase |
| | P_0505 | Uncharacterized lipoprotein |
| | P_0661 | 50S ribosomal protein L14 |
| | P_0479 | Uncharacterized peptidase |
| | P_0375 | Uncharacterized protein |
| | P_0005 | Uncharacterized protein |
| | P_0353 | Cell division DivIVA/GpsB protein |
| | P_0400 | Uncharacterized protease |
| | P_0887 | Coenzyme A Disulfide Reductase |
| | P_0403 | Ribosome GTPase |
| | P_0441 | Cysteine desulfurase |
| | P_0642 | ECF transporter ATPase |

*continued on next page*

Table S2 – *continued from previous page*

| Module | Protein ID | Function |
|---|---|---|
| | P_0168 | Oligopeptide ABC transporter ATP-binding protein |
| | P_0332 | Uncharacterized protein |
| | P_0482 | 30S ribosomal protein S21 |
| | P_0686 | Potassium transporter |
| | P_0302 | Flavin reductase |
| | P_0387 | tRNA 2-thiouridine(34) synthase |
| | P_0347 | Cytidylate kinase |
| | P_0214 | Phosphatidylglycerophosphatase |
| | P_0392 | Uncharacterized protein |
| | P_0316 | Transketolase |
| | P_0690 | DNA ligase (NAD(+)) |
| | P_0166 | Oligopeptide ABC transporter permease |
| | P_0799 | Serine hydroxymethyltransferase |
| | P_0530 | Uncharacterized protein |
| | P_0143 | Uncharacterized protein |
| | P_0549 | Non-canonical purine NTP pyrophosphatase |
| | P_0009 | Nucleoside ABC transporter permease |
| | P_0418 | Ribonuclease III |
| | P_0044 | DNA polymerase III subunit delta |
| | P_0452 | DNA topoisomerase IV subunit B |
| | P_0805 | Uncharacterized protein |
| | P_0877 | Riboflavin ECF transporter S component |
| | P_0065 | Thioredoxin |
| | P_0095 | Preprotein translocase subunit A |
| | P_0139 | NanoRNAse |

*continued on next page*

Table S2 – *continued from previous page*

| Module | Protein ID | Function |
|---|---|---|
| | P_0402 | rRNA maturation RNase |
| | P_0523 | Cell division protein |
| | P_0428 | Phosphate transport system regulatory protein |
| | P_0137 | 50S ribosomal protein L31 |
| | P_0906 | Uncharacterized protein |
| | P_0147 | Cardiolipin synthase |
| | P_0361 | 23S rRNA (pseudouridine(1915)-N3)-methyltransferase |
| | P_0439 | Uncharacterized lipoprotein |
| | P_0004 | 16S rRNA (adenine(1518)-N(6)/adenine(1519)-N(6))- dimethyltransferase |
| | P_0537 | UMP kinase |
| | P_0228 | Dihydrolipoyl dehydrogenase |
| | P_0063 | Uncharacterized tRNA dihydrouridine synthase |
| | P_0544 | Heat-inducible transcription repressor |
| | P_0517 | Uncharacterized RNA pseudouridine synthase |
| | P_0792 | F0F1 ATP synthase subunit alpha |
| | P_0821 | HPr(Ser) kinase/phosphatase |
| | P_0301 | Ribosome assembly cofactor |
| | P_0346 | Uncharacterized protein |
| | P_0823 | Dihydrofolate synthase |
| | P_0707 | Thiamine ABC transporter ATP-binding protein |
| | P_0002 | DNA polymerase III subunit beta |
| Module 12 | P_0304 | Phosphatidate cytidylyltransferase |
| | P_0792 | F0F1 ATP synthase subunit alpha |

*continued on next page*

Table S2 – *continued from previous page*

| **Module** | **Protein ID** | **Function** |
|---|---|---|
| | P_0605 | Uncharacterized protein |
| | P_0115 | UTP–glucose-1-phosphate uridylyltransferase |
| | P_0046 | Recombination protein |
| | P_0418 | Ribonuclease III |
| | P_0791 | F0F1 ATP synthase subunit gamma |
| | P_0876 | Uncharacterized amino acid permease |
| | P_0271 | tRNA N6-adenosine(37)-N6-threonylcarbamoyltransferase complex dimerization subunit |
| | P_0422 | 50S ribosomal protein L28 |
| | P_0592 | Uncharacterized protein |
| | P_0106 | Exodeoxyribonuclease VII large subunit |
| | P_0695 | ATP-dependent DNA helicase |
| | P_0726 | Glucosamine-6-phosphate deaminase |
| | P_0132 | Toxin-antitoxin AAA ATPase |
| | P_0148 | 30S ribosomal protein S12 |
| | P_0636 | Uncharacterized lipoprotein |
| | P_0651 | Adenylate kinase |
| | P_0859 | DNA topoisomerase I |
| | P_0377 | Ribosome GTPase |
| | P_0314 | Uncharacterized ECF transporter S component |
| | P_0215 | Pre-16S rRNA nuclease |
| | P_0360 | Signal recognition particle protein |
| | P_0834 | Replicative DNA helicase |
| | P_0054 | Uncharacterized peroxiredoxin |
| | P_0047 | DNA polymerase III subunit gamma and tau |

*continued on next page*

Table S2 – *continued from previous page*

| Module | Protein ID | Function |
|---|---|---|
| | P_0500 | Maturation protease for ribosomal protein L27 |
| | P_0838 | 23S rRNA (guanosine(2251)-2'-O)-methyltransferase |
| | P_0316 | Transketolase |
| | P_0138 | Uncharacterized protein |
| | P_0301 | Ribosome assembly cofactor |
| | P_0515 | Cytidine deaminase |
| | P_0430 | Uncharacterized DNA-binding protein |
| | P_0684 | Bifunctional 5,10-methylene-tetrahydrofolate dehydrogenase/5,10-methylene-tetrahydrofolate cyclohydrolase |
| | P_0113 | Glycolipid synthase A |
| | P_0063 | Uncharacterized tRNA dihydrouridine synthase |
| | P_0453 | DNA topoisomerase IV subunit A |
| | P_0825 | Excinuclease ABC subunit B |
| | P_0114 | Glycolipid synthase B |
| | P_0874 | 16S rRNA (guanine(527)-N(7))-methyltransferase |
| | P_0451 | Glyceraldehyde-3-phosphate dehydrogenase |
| | P_0364 | tRNA (guanosine(37)-N1)-methyltransferase |
| | P_0407 | RNA polymerase subunit sigma |
| | P_0523 | Cell division protein |
| | P_0799 | Serine hydroxymethyltransferase |
| | P_0065 | Thioredoxin |
| | P_0800 | Ribose 5-phosphate isomerase B |
| | P_0549 | Non-canonical purine NTP pyrophosphatase |

*continued on next page*

Table S2 – *continued from previous page*

| Module | Protein ID | Function |
|---|---|---|
| | P_0505 | Uncharacterized lipoprotein |
| | P_0195 | Spermidine/putrescine ABC transporter permease |
| | P_0793 | F0F1 ATP synthase subunit delta |
| | P_0794 | F0F1 ATP synthase subunit B |
| | P_0009 | Nucleoside ABC transporter permease |
| | P_0382 | Deoxynucleoside kinase |
| | P_0524 | 16S rRNA (cytosine(1402)-N(4))-methyltransferase |
| | P_0302 | Flavin reductase |
| | P_0538 | Uncharacterized protein |
| | P_0908 | Membrane protein insertase |
| | P_0030 | Uncharacterized ABC transporter ATP-binding protein |
| | P_0081 | tRNA uridine(34) 5-carboxymethylaminomethyl synthesis GTPase |
| | P_0885 | tRNA uridine(34) 5-carboxymethylaminomethyl synthesis enzyme |
| | P_0097 | Flap endonuclease and 5' exonuclease |
| | P_0710 | Low specificity hydrolase |
| | P_0821 | HPr(Ser) kinase/phosphatase |
| | P_0644 | 50S ribosomal protein L17 |
| Module 13 | P_0348 | Ribosome biogenesis GTPase |
| | P_0387 | tRNA 2-thiouridine(34) synthase |
| | P_0407 | RNA polymerase subunit sigma |
| | P_0513 | ACP synthase |

*continued on next page*

Table S2 – *continued from previous page*

| Module | Protein ID | Function |
|---|---|---|
| | P_0527 | Uncharacterized protein |
| | P_0501 | 50S ribosomal protein L21 |
| | P_0710 | Low specificity hydrolase |
| | P_0065 | Thioredoxin |
| | P_0494 | N-acetylmannosamine-6-phosphate 2-epimerase |
| | P_0775 | Ribonuclease R |
| | P_0427 | Phosphate ABC transporter ATP-binding protein |
| | P_0689 | Glutamyl-tRNA amidotransferase subunit C |
| | P_0168 | Oligopeptide ABC transporter ATP-binding protein |
| | P_0605 | Uncharacterized protein |
| | P_0789 | F0F1 ATP synthase subunit delta/epsilon |
| | P_0453 | DNA topoisomerase IV subunit A |
| | P_0197 | Spermidine/putrescine ABC transporter ATP-binding protein |
| | P_0821 | HPr(Ser) kinase/phosphatase |
| | P_0687 | Glutamyl-tRNA amidotransferase subunit B |
| | P_0139 | NanoRNAse |
| | P_0429 | Signal recognition particle-docking protein |
| | P_0113 | Glycolipid synthase A |
| | P_0825 | Excinuclease ABC subunit B |
| | P_0249 | Uncharacterized protein |
| | P_0344 | Inorganic diphosphatase |
| | P_0039 | Membrane anchored AAA+ protease |

*continued on next page*

Table S2 – *continued from previous page*

| Module | Protein ID | Function |
|---|---|---|
| | P_0346 | Uncharacterized protein |
| | P_0422 | 50S ribosomal protein L28 |
| | P_0229 | Phosphate acetyltransferase |
| | P_0005 | Uncharacterized protein |
| | P_0684 | Bifunctional 5,10-methylene-tetrahydrofolate dehydrogenase/5,10-methylene-tetrahydrofolate cyclohydrolase |
| | P_0805 | Uncharacterized protein |
| | P_0511 | Uncharacterized protein |
| | P_0140 | Thymidine kinase |
| | P_0166 | Oligopeptide ABC transporter permease |
| | P_0885 | tRNA uridine(34) 5-carboxymethylaminomethyl synthesis enzyme |
| | P_0167 | Oligopeptide ABC transporter ATP-binding protein |
| | P_0876 | Uncharacterized amino acid permease |
| | P_0530 | Uncharacterized protein |
| | P_0377 | Ribosome GTPase |
| | P_0199 | 50S ribosomal protein L35 |
| | P_0202 | 16S rRNA (guanine(966)-N(2))-methyltransferase |
| | P_0347 | Cytidylate kinase |
| | P_0145 | Uncharacterized acetyltransferase |
| | P_0451 | Glyceraldehyde-3-phosphate dehydrogenase |
| | P_0026 | Single-stranded DNA-binding protein |
| | P_0042 | Uncharacterized transcriptional regulator |

*continued on next page*

Table S2 – *continued from previous page*

| Module | Protein ID | Function |
|---|---|---|
| | P_0434 | 23S rRNA (uridine(1939)-m5)-methyltransferase |
| | P_0006 | DNA gyrase subunit B |
| | P_0544 | Heat-inducible transcription repressor |
| Module 14 | P_0623 | Uncharacterized protein |
| | P_0501 | 50S ribosomal protein L21 |
| | P_0131 | Fructose-1,6-bisphosphate aldolase |
| | P_0440 | Uncharacterized lipoprotein |
| | P_0167 | Oligopeptide ABC transporter ATP-binding protein |
| | P_0305 | Prolidase |
| | P_0330 | Deoxyguanosine kinase |
| | P_0451 | Glyceraldehyde-3-phosphate dehydrogenase |
| | P_0094 | Uncharacterized protein |
| | P_0291 | FAD synthetase |
| | P_0129 | CTP synthase |
| | P_0005 | Uncharacterized protein |
| | P_0851 | Uncharacterized lipoprotein |
| | P_0611 | DNA polymerase I |
| | P_0792 | F0F1 ATP synthase subunit alpha |
| | P_0372 | Flippase B |
| | P_0445 | Glucose-6-phosphate isomerase |
| | P_0887 | Coenzyme A Disulfide Reductase |
| | P_0615 | Uncharacterized protein |
| | P_0377 | Ribosome GTPase |
| | P_0249 | Uncharacterized protein |

*continued on next page*

Table S2 – *continued from previous page*

| Module | Protein ID | Function |
|---|---|---|
| | P_0350 | DNA-binding protein |
| | P_0513 | ACP synthase |
| | P_0133 | Toxin-antitoxin serine protease |
| | P_0420 | Fatty acid kinase subunit A |
| | P_0826 | DNA polymerase III subunit delta |
| | P_0044 | DNA polymerase III subunit delta |
| | P_0045 | dTMP kinase |
| | P_0063 | Uncharacterized tRNA dihydrouridine synthase |
| | P_0827 | Uncharacterized protein |
| | P_0026 | Single-stranded DNA-binding protein |
| | P_0140 | Thymidine kinase |
| | P_0708 | Thiamine ABC transporter substrate-binding protein |
| | P_0651 | Adenylate kinase |
| | P_0054 | Uncharacterized peroxiredoxin |
| | P_0407 | RNA polymerase subunit sigma |
| | P_0113 | Glycolipid synthase A |
| | P_0541 | Molecular chaperone |
| | P_0344 | Inorganic diphosphatase |
| | P_0429 | Signal recognition particle-docking protein |
| | P_0250 | Uncharacterized protein |
| | P_0840 | Antitermination protein |
| | P_0359 | Ribonuclease Y |
| | P_0881 | Uncharacterized MFS transporter |
| | P_0799 | Serine hydroxymethyltransferase |
| | P_0338 | Uncharacterized lipoprotein |

*continued on next page*

Table S2 – *continued from previous page*

| Module | Protein ID | Function |
|---|---|---|
| | P_0165 | Oligopeptide ABC transporter permease |
| | P_0814 | UDP-galactopyranose mutase |
| | P_0434 | 23S rRNA (uridine(1939)-m5)-methyltransferase |
| | P_0389 | Uncharacterized protein |
| | P_0548 | tRNA (cytidine(34)-2'-O)-methyltransferase |
| | P_0294 | 30S ribosomal protein S15 |
| | P_0127 | Uncharacterized phosphohydrolase |
| | P_0201 | Peptide deformylase |
| | P_0200 | Translation initiation factor IF-3 |
| | P_0495 | Uncharacterized kinase |
| Module 15 | P_0080 | Uncharacterized protein |
| | P_0238 | 30S ribosomal protein S4 |
| | P_0140 | Thymidine kinase |
| | P_0652 | Preprotein translocase subunit |
| | P_0645 | DNA-directed RNA polymerase subunit alpha |
| | P_0443 | 5-Formyltetrahydrofolate cyclo-ligase |
| | P_0410 | Degradosome RNA helicase |
| | P_0771 | Ribonucleotide-diphosphate reductase subunit alpha |
| | P_0440 | Uncharacterized lipoprotein |
| | P_0644 | 50S ribosomal protein L17 |
| | P_0668 | 50S ribosomal protein L2 |
| | P_0671 | 50S ribosomal protein L3 |
| | P_0146 | Uncharacterized protein |
| | P_0065 | Thioredoxin |

*continued on next page*

Table S2 – *continued from previous page*

| Module | Protein ID | Function |
|---|---|---|
| | P_0026 | Single-stranded DNA-binding protein |
| | P_0549 | Non-canonical purine NTP pyrophosphatase |
| | P_0526 | 50S ribosomal protein L32 |
| | P_0695 | ATP-dependent DNA helicase |
| | P_0794 | F0F1 ATP synthase subunit B |
| | P_0710 | Low specificity hydrolase |
| | P_0347 | Cytidylate kinase |
| | P_0544 | Heat-inducible transcription repressor |
| | P_0500 | Maturation protease for ribosomal protein L27 |
| | P_0047 | DNA polymerase III subunit gamma and tau |
| | P_0451 | Glyceraldehyde-3-phosphate dehydrogenase |
| | P_0870 | Uncharacterized C4-dicarboxylate ABC transporter |
| | P_0148 | 30S ribosomal protein S12 |
| | P_0415 | Chromosome segregation protein |
| | P_0832 | Aminoacyl-tRNA hydrolase |
| | P_0516 | Uncharacterized protein |
| | P_0371 | Flippase A |
| | P_0730 | Uncharacterized protein |
| | P_0499 | 50S ribosomal protein L27 |
| | P_0684 | Bifunctional 5,10-methylene-tetrahydrofolate dehydrogenase/5,10-methylene-tetrahydrofolate cyclohydrolase |
| | P_0294 | 30S ribosomal protein S15 |
| | P_0799 | Serine hydroxymethyltransferase |
| | P_0776 | SsrA-binding protein |

*continued on next page*

Table S2 – *continued from previous page*

| Module | Protein ID | Function |
|---|---|---|
| | P_0005 | Uncharacterized protein |
| | P_0316 | Transketolase |
| | P_0537 | UMP kinase |
| | P_0009 | Nucleoside ABC transporter permease |
| | P_0444 | Oligopeptidase |
| | P_0834 | Replicative DNA helicase |
| | P_0113 | Glycolipid synthase A |
| | P_0003 | Ribonuclease M5 |
| | P_0330 | Deoxyguanosine kinase |
| | P_0378 | NAD(+) synthase |
| | P_0029 | FMN reductase |
| | P_0202 | 16S rRNA (guanine(966)-N(2))-methyltransferase |
| | P_0359 | Ribonuclease Y |
| | P_0117 | Acyl-phosphate glycerol 3-phosphate acyltransferase |
| | P_0420 | Fatty acid kinase subunit A |
| | P_0302 | Flavin reductase |
| | P_0727 | Triose-phosphate isomerase |
| | P_0421 | Uncharacterized protein |
| | P_0707 | Thiamine ABC transporter ATP-binding protein |
| | P_0390 | Methionyl-tRNA formyltransferase |
| | P_0414 | Guanosine-3',5'-bis(diphosphate) 3'-pyrophosphohydrolase |

*continued on next page*

Table S2 – *continued from previous page*

| Module | Protein ID | Function |
|---|---|---|
| | P_0874 | 16S rRNA (guanine(527)-N(7))-methyltransferase |
| | P_0887 | Coenzyme A Disulfide Reductase |
| | P_0827 | Uncharacterized protein |
| | P_0792 | F0F1 ATP synthase subunit alpha |
| | P_0651 | Adenylate kinase |
| | P_0886 | Proton-glutamate symporter |
| Module 16 | P_0431 | Uncharacterized metallophosphoesterase |
| | P_0426 | Phosphate ABC transporter permease |
| | P_0670 | 50S ribosomal protein L4 |
| | P_0650 | Type I methionyl aminopeptidase |
| | P_0643 | ECF transporter ATPase |
| | P_0434 | 23S rRNA (uridine(1939)-m5)-methyltransferase |
| | P_0392 | Uncharacterized protein |
| | P_0644 | 50S ribosomal protein L17 |
| | P_0693 | Uncharacterized protease |
| | P_0453 | DNA topoisomerase IV subunit A |
| | P_0407 | RNA polymerase subunit sigma |
| | P_0151 | Translation elongation factor Tu |
| | P_0138 | Uncharacterized protein |
| | P_0389 | Uncharacterized protein |
| | P_0669 | 50S ribosomal protein L23 |
| | P_0726 | Glucosamine-6-phosphate deaminase |
| | P_0007 | DNA gyrase subunit A |
| | P_0346 | Uncharacterized protein |

*continued on next page*

Table S2 – *continued from previous page*

| Module | Protein ID | Function |
|---|---|---|
| | P_0421 | Uncharacterized protein |
| | P_0520 | Uncharacterized hydrolase |
| | P_0435 | Mannose-6-phosphate isomerase |
| | P_0126 | Glutamate–tRNA ligase |
| | P_0792 | F0F1 ATP synthase subunit alpha |
| | P_0127 | Uncharacterized phosphohydrolase |
| | P_0772 | Ribonucleotide reductase assembly protein |
| | P_0432 | Methionine adenosyltransferase |
| | P_0365 | 50S ribosomal protein L19 |
| | P_0690 | DNA ligase (NAD(+)) |
| | P_0375 | Uncharacterized protein |
| | P_0605 | Uncharacterized protein |
| | P_0388 | Uncharacterized protein |
| | P_0515 | Cytidine deaminase |
| | P_0505 | Uncharacterized lipoprotein |
| | P_0799 | Serine hydroxymethyltransferase |
| | P_0418 | Ribonuclease III |
| | P_0042 | Uncharacterized transcriptional regulator |
| | P_0789 | F0F1 ATP synthase subunit delta/epsilon |
| | P_0600 | Ribonuclease J |
| | P_0494 | N-acetylmannosamine-6-phosphate 2-epimerase |
| | P_0410 | Degradosome RNA helicase |
| | P_0642 | ECF transporter ATPase |
| | P_0592 | Uncharacterized protein |
| | P_0137 | 50S ribosomal protein L31 |

*continued on next page*

Table S2 – *continued from previous page*

| Module | Protein ID | Function |
|---|---|---|
| | P_0154 | Uncharacterized peptidase |
| | P_0362 | 30S ribosomal protein S16 |
| | P_0614 | Nicotinate phosphoribosyltransferase |
| | P_0391 | Elongation factor P |
| | P_0684 | Bifunctional 5,10-methylene-tetrahydrofolate dehydrogenase/5,10-methylene-tetrahydrofolate cyclohydrolase |
| | P_0687 | Glutamyl-tRNA amidotransferase subunit B |
| | P_0387 | tRNA 2-thiouridine(34) synthase |
| | P_0870 | Uncharacterized C4-dicarboxylate ABC transporter |
| | P_0443 | 5-Formyltetrahydrofolate cyclo-ligase |
| | P_0218 | Glycerol kinase |
| | P_0314 | Uncharacterized ECF transporter S component |
| | P_0316 | Transketolase |
| | P_0026 | Single-stranded DNA-binding protein |
| | P_0373 | Uncharacterized protein |
| | P_0686 | Potassium transporter |
| | P_0501 | 50S ribosomal protein L21 |
| | P_0697 | Uncharacterized glycosyl transferase |
| | P_0113 | Glycolipid synthase A |
| | P_0513 | ACP synthase |
| | P_0116 | Uncharacterized protein |
| | P_0445 | Glucose-6-phosphate isomerase |
| Module 17 | P_0372 | Flippase B |
| | P_0499 | 50S ribosomal protein L27 |

*continued on next page*

Table S2 – *continued from previous page*

| Module | Protein ID | Function |
|---|---|---|
| | P_0824 | Excinuclease ABC subunit A |
| | P_0127 | Uncharacterized phosphohydrolase |
| | P_0669 | 50S ribosomal protein L23 |
| | P_0733 | Phosphopentomutase |
| | P_0378 | NAD(+) synthase |
| | P_0835 | Uncharacterized lipoprotein |
| | P_0710 | Low specificity hydrolase |
| | P_0148 | 30S ribosomal protein S12 |
| | P_0042 | Uncharacterized transcriptional regulator |
| | P_0314 | Uncharacterized ECF transporter S component |
| | P_0411 | Uncharacterized protein |
| | P_0347 | Cytidylate kinase |
| | P_0516 | Uncharacterized protein |
| | P_0094 | Uncharacterized protein |
| | P_0513 | ACP synthase |
| | P_0203 | Guanylate kinase |
| | P_0165 | Oligopeptide ABC transporter permease |
| | P_0435 | Mannose-6-phosphate isomerase |
| | P_0517 | Uncharacterized RNA pseudouridine synthase |
| | P_0549 | Non-canonical purine NTP pyrophosphatase |
| | P_0652 | Preprotein translocase subunit |
| | P_0429 | Signal recognition particle-docking protein |
| | P_0697 | Uncharacterized glycosyl transferase |
| | P_0414 | Guanosine-3',5'-bis(diphosphate) 3'-pyrophosphohydrolase |
| | P_0799 | Serine hydroxymethyltransferase |

*continued on next page*

Table S2 – *continued from previous page*

| Module | Protein ID | Function |
|---|---|---|
| | P_0007 | DNA gyrase subunit A |
| | P_0002 | DNA polymerase III subunit beta |
| | P_0129 | CTP synthase |
| | P_0608 | Primosomal protein |
| | P_0526 | 50S ribosomal protein L32 |
| | P_0169 | Oligopeptide ABC transporter substrate-binding protein |
| | P_0338 | Uncharacterized lipoprotein |
| | P_0081 | tRNA uridine(34) 5-carboxymethylaminomethyl synthesis GTPase |
| | P_0872 | Uncharacterized ATPase |
| | P_0359 | Ribonuclease Y |
| | P_0859 | DNA topoisomerase I |
| | P_0821 | HPr(Ser) kinase/phosphatase |
| | P_0482 | 30S ribosomal protein S21 |
| | P_0026 | Single-stranded DNA-binding protein |
| | P_0200 | Translation initiation factor IF-3 |
| | P_0167 | Oligopeptide ABC transporter ATP-binding protein |
| | P_0814 | UDP-galactopyranose mutase |
| | P_0406 | DNA primase |
| | P_0285 | Elongation factor 4 |
| | P_0617 | Fatty acid binding protein |
| | P_0116 | Uncharacterized protein |
| | P_0639 | Uncharacterized efflux ABC transporter permease |

*continued on next page*

Table S2 – *continued from previous page*

| Module | Protein ID | Function |
|---|---|---|
| | P_0063 | Uncharacterized tRNA dihydrouridine synthase |
| | P_0302 | Flavin reductase |
| | P_0541 | Molecular chaperone |
| | P_0392 | Uncharacterized protein |
| | P_0044 | DNA polymerase III subunit delta |
| | P_0728 | Low specificity hydrolase |
| | P_0197 | Spermidine/putrescine ABC transporter ATP-binding protein |
| | P_0544 | Heat-inducible transcription repressor |
| | P_0010 | Nucleoside ABC transporter ATP-binding protein |
| | P_0114 | Glycolipid synthase B |
| Module 18 | P_0779 | PTS sugar transporter |
| | P_0271 | tRNA N6-adenosine(37)-N6-threonylcarbamoyltransferase complex dimerization subunit |
| | P_0851 | Uncharacterized lipoprotein |
| | P_0601 | Uncharacterized protein |
| | P_0688 | Glutamyl-tRNA amidotransferase subunit A |
| | P_0814 | UDP-galactopyranose mutase |
| | P_0687 | Glutamyl-tRNA amidotransferase subunit B |
| | P_0409 | Uncharacterized protein |
| | P_0908 | Membrane protein insertase |
| | P_0668 | 50S ribosomal protein L2 |
| | P_0661 | 50S ribosomal protein L14 |
| | P_0045 | dTMP kinase |

*continued on next page*

Table S2 – *continued from previous page*

| Module | Protein ID | Function |
|---|---|---|
| | P_0329 | Uncharacterized pseudouridine synthase |
| | P_0026 | Single-stranded DNA-binding protein |
| | P_0648 | 50S ribosomal protein L36 |
| | P_0302 | Flavin reductase |
| | P_0082 | 30S ribosomal protein S20 |
| | P_0116 | Uncharacterized protein |
| | P_0787 | Magnesium-translocating P-type ATPase |
| | P_0165 | Oligopeptide ABC transporter permease |
| | P_0544 | Heat-inducible transcription repressor |
| | P_0410 | Degradosome RNA helicase |
| | P_0195 | Spermidine/putrescine ABC transporter permease |
| | P_0200 | Translation initiation factor IF-3 |
| | P_0726 | Glucosamine-6-phosphate deaminase |
| | P_0650 | Type I methionyl aminopeptidase |
| | P_0710 | Low specificity hydrolase |
| | P_0338 | Uncharacterized lipoprotein |
| | P_0600 | Ribonuclease J |
| | P_0132 | Toxin-antitoxin AAA ATPase |
| | P_0361 | 23S rRNA (pseudouridine(1915)-N3)-methyltransferase |
| | P_0377 | Ribosome GTPase |
| | P_0530 | Uncharacterized protein |
| | P_0727 | Triose-phosphate isomerase |
| | P_0453 | DNA topoisomerase IV subunit A |
| | P_0444 | Oligopeptidase |

*continued on next page*

Table S2 – *continued from previous page*

| **Module** | **Protein ID** | **Function** |
|---|---|---|
| | P_0452 | DNA topoisomerase IV subunit B |
| | P_0874 | 16S rRNA (guanine(527)-N(7))-methyltransferase |
| | P_0615 | Uncharacterized protein |
| | P_0537 | UMP kinase |
| | P_0695 | ATP-dependent DNA helicase |
| | P_0793 | F0F1 ATP synthase subunit delta |
| | P_0140 | Thymidine kinase |
| | P_0127 | Uncharacterized phosphohydrolase |
| | P_0247 | Ribosome biogenesis GTP-binding protein |
| | P_0366 | L16-binding dependent 50S subunit-maturation GTPase |
| | P_0391 | Elongation factor P |
| | P_0168 | Oligopeptide ABC transporter ATP-binding protein |
| | P_0610 | DNA-formamidopyrimidine glycosylase |
| | P_0001 | Chromosomal replication initiator protein |
| | P_0316 | Transketolase |
| | P_0412 | Uncharacterized protein |
| | P_0512 | Acyl-phosphate glycerol 3-phosphate acyltransferase |
| | P_0859 | DNA topoisomerase I |
| | P_0728 | Low specificity hydrolase |
| Module 19 | P_0005 | Uncharacterized protein |
| | P_0045 | dTMP kinase |
| | P_0805 | Uncharacterized protein |

*continued on next page*

Table S2 – *continued from previous page*

| Module | Protein ID | Function |
|---|---|---|
| | P_0410 | Degradosome RNA helicase |
| | P_0601 | Uncharacterized protein |
| | P_0428 | Phosphate transport system regulatory protein |
| | P_0881 | Uncharacterized MFS transporter |
| | P_0407 | RNA polymerase subunit sigma |
| | P_0775 | Ribonuclease R |
| | P_0521 | Cell division protein |
| | P_0639 | Uncharacterized efflux ABC transporter permease |
| | P_0789 | F0F1 ATP synthase subunit delta/epsilon |
| | P_0137 | 50S ribosomal protein L31 |
| | P_0097 | Flap endonuclease and 5' exonuclease |
| | P_0132 | Toxin-antitoxin AAA ATPase |
| | P_0145 | Uncharacterized acetyltransferase |
| | P_0875 | CDP-diacylglycerol–glycerol-3-phosphate 3-phosphatidyltransferase |
| | P_0791 | F0F1 ATP synthase subunit gamma |
| | P_0728 | Low specificity hydrolase |
| | P_0228 | Dihydrolipoyl dehydrogenase |
| | P_0371 | Flippase A |
| | P_0427 | Phosphate ABC transporter ATP-binding protein |
| | P_0195 | Spermidine/putrescine ABC transporter permease |
| | P_0872 | Uncharacterized ATPase |
| | P_0006 | DNA gyrase subunit B |

*continued on next page*

Table S2 – *continued from previous page*

| Module | Protein ID | Function |
| --- | --- | --- |
| | P_0512 | Acyl-phosphate glycerol 3-phosphate acyltransferase |
| | P_0116 | Uncharacterized protein |
| | P_0729 | Phosphoglycerate mutase (2,3-diphosphoglycerate-independent) |
| | P_0772 | Ribonucleotide reductase assembly protein |
| | P_0452 | DNA topoisomerase IV subunit B |
| | P_0411 | Uncharacterized protein |
| | P_0391 | Elongation factor P |
| | P_0302 | Flavin reductase |
| | P_0541 | Molecular chaperone |
| | P_0821 | HPr(Ser) kinase/phosphatase |
| | P_0494 | N-acetylmannosamine-6-phosphate 2-epimerase |
| | P_0203 | Guanylate kinase |
| | P_0726 | Glucosamine-6-phosphate deaminase |
| | P_0451 | Glyceraldehyde-3-phosphate dehydrogenase |
| | P_0138 | Uncharacterized protein |
| | P_0859 | DNA topoisomerase I |
| | P_0438 | Uncharacterized protein |
| | P_0197 | Spermidine/putrescine ABC transporter ATP-binding protein |
| | P_0549 | Non-canonical purine NTP pyrophosphatase |
| | P_0614 | Nicotinate phosphoribosyltransferase |
| | P_0199 | 50S ribosomal protein L35 |

*continued on next page*

Table S2 – *continued from previous page*

| Module | Protein ID | Function |
|---|---|---|
| | P_0366 | L16-binding dependent 50S subunit-maturation GTPase |
| | P_0054 | Uncharacterized peroxiredoxin |
| | P_0707 | Thiamine ABC transporter ATP-binding protein |
| | P_0316 | Transketolase |
| | P_0200 | Translation initiation factor IF-3 |
| | P_0794 | F0F1 ATP synthase subunit B |
| | P_0359 | Ribonuclease Y |
| | P_0732 | Deoxyribose-phosphate aldolase |
| | P_0838 | 23S rRNA (guanosine(2251)-2'-O)-methyltransferase |
| Module 20 | P_0140 | Thymidine kinase |
| | P_0285 | Elongation factor 4 |
| | P_0697 | Uncharacterized glycosyl transferase |
| | P_0824 | Excinuclease ABC subunit A |
| | P_0011 | Nucleoside ABC transporter substrate-binding protein |
| | P_0439 | Uncharacterized lipoprotein |
| | P_0402 | rRNA maturation RNase |
| | P_0407 | RNA polymerase subunit sigma |
| | P_0687 | Glutamyl-tRNA amidotransferase subunit B |
| | P_0825 | Excinuclease ABC subunit B |
| | P_0203 | Guanylate kinase |
| | P_0793 | F0F1 ATP synthase subunit delta |
| | P_0733 | Phosphopentomutase |

*continued on next page*

Table S2 – *continued from previous page*

| Module | Protein ID | Function |
|---|---|---|
| | P_0427 | Phosphate ABC transporter ATP-binding protein |
| | P_0494 | N-acetylmannosamine-6-phosphate 2-epimerase |
| | P_0005 | Uncharacterized protein |
| | P_0365 | 50S ribosomal protein L19 |
| | P_0314 | Uncharacterized ECF transporter S component |
| | P_0832 | Aminoacyl-tRNA hydrolase |
| | P_0400 | Uncharacterized protease |
| | P_0499 | 50S ribosomal protein L27 |
| | P_0168 | Oligopeptide ABC transporter ATP-binding protein |
| | P_0453 | DNA topoisomerase IV subunit A |
| | P_0874 | 16S rRNA (guanine(527)-N(7))-methyltransferase |
| | P_0872 | Uncharacterized ATPase |
| | P_0116 | Uncharacterized protein |
| | P_0420 | Fatty acid kinase subunit A |
| | P_0836 | Uncharacterized ECF transporter S component |
| | P_0800 | Ribose 5-phosphate isomerase B |
| | P_0775 | Ribonuclease R |
| | P_0641 | ECF transporter T component |
| | P_0033 | Uncharacterized protein |
| | P_0821 | HPr(Ser) kinase/phosphatase |
| | P_0195 | Spermidine/putrescine ABC transporter permease |

*continued on next page*

Table S2 – *continued from previous page*

| **Module** | **Protein ID** | **Function** |
|---|---|---|
| | P_0434 | 23S rRNA (uridine(1939)-m5)-methyltransferase |
| | P_0412 | Uncharacterized protein |
| | P_0835 | Uncharacterized lipoprotein |
| | P_0428 | Phosphate transport system regulatory protein |
| | P_0512 | Acyl-phosphate glycerol 3-phosphate acyltransferase |
| | P_0605 | Uncharacterized protein |
| | P_0063 | Uncharacterized tRNA dihydrouridine synthase |
| | P_0887 | Coenzyme A Disulfide Reductase |
| | P_0141 | Peptide chain release factor 1 |
| | P_0707 | Thiamine ABC transporter ATP-binding protein |
| | P_0859 | DNA topoisomerase I |
| | P_0805 | Uncharacterized protein |
| | P_0875 | CDP-diacylglycerol–glycerol-3-phosphate 3-phosphatidyltransferase |
| | P_0001 | Chromosomal replication initiator protein |
| | P_0834 | Replicative DNA helicase |
| | P_0201 | Peptide deformylase |
| | P_0549 | Non-canonical purine NTP pyrophosphatase |
| | P_0451 | Glyceraldehyde-3-phosphate dehydrogenase |
| | P_0291 | FAD synthetase |
| | P_0378 | NAD(+) synthase |
| | P_0688 | Glutamyl-tRNA amidotransferase subunit A |
| | P_0139 | NanoRNAse |

*continued on next page*

Table S2 – *continued from previous page*

| **Module** | **Protein ID** | **Function** |
|---|---|---|
| | P_0065 | Thioredoxin |
| Module 21 | P_0642 | ECF transporter ATPase |
| | P_0326 | Uncharacterized protein |
| | P_0114 | Glycolipid synthase B |
| | P_0505 | Uncharacterized lipoprotein |
| | P_0299 | Uncharacterized protein |
| | P_0063 | Uncharacterized tRNA dihydrouridine synthase |
| | P_0814 | UDP-galactopyranose mutase |
| | P_0129 | CTP synthase |
| | P_0407 | RNA polymerase subunit sigma |
| | P_0254 | Excinuclease ABC subunit C |
| | P_0366 | L16-binding dependent 50S subunit-maturation GTPase |
| | P_0011 | Nucleoside ABC transporter substrate-binding protein |
| | P_0371 | Flippase A |
| | P_0065 | Thioredoxin |
| | P_0249 | Uncharacterized protein |
| | P_0410 | Degradosome RNA helicase |
| | P_0045 | dTMP kinase |
| | P_0247 | Ribosome biogenesis GTP-binding protein |
| | P_0799 | Serine hydroxymethyltransferase |
| | P_0435 | Mannose-6-phosphate isomerase |
| | P_0301 | Ribosome assembly cofactor |
| | P_0859 | DNA topoisomerase I |
| | P_0710 | Low specificity hydrolase |

*continued on next page*

Table S2 – *continued from previous page*

| **Module** | **Protein ID** | **Function** |
|---|---|---|
| | P_0145 | Uncharacterized acetyltransferase |
| | P_0168 | Oligopeptide ABC transporter ATP-binding protein |
| | P_0420 | Fatty acid kinase subunit A |
| | P_0932 | 50S ribosomal protein L33 |
| | P_0800 | Ribose 5-phosphate isomerase B |
| | P_0006 | DNA gyrase subunit B |
| | P_0029 | FMN reductase |
| | P_0792 | F0F1 ATP synthase subunit alpha |
| | P_0813 | UDP-glucose 4-epimerase GalE |
| | P_0202 | 16S rRNA (guanine(966)-N(2))-methyltransferase |
| | P_0144 | tRNA (adenosine(37)-N6)-threonylcarbamoyltransferase complex transferase subunit |
| | P_0040 | tRNA lysidine(34) synthetase |
| | P_0708 | Thiamine ABC transporter substrate-binding protein |
| | P_0054 | Uncharacterized peroxiredoxin |
| | P_0387 | tRNA 2-thiouridine(34) synthase |
| | P_0195 | Spermidine/putrescine ABC transporter permease |
| | P_0043 | Uncharacterized methyltransferase |
| | P_0616 | Fatty acid binding protein |
| | P_0885 | tRNA uridine(34) 5-carboxymethylaminomethyl synthesis enzyme |

*continued on next page*

Table S2 – *continued from previous page*

| **Module** | **Protein ID** | **Function** |
|---|---|---|
| | P_0001 | Chromosomal replication initiator protein |
| | P_0004 | 16S rRNA (adenine(1518)-N(6)/adenine(1519)-N(6))- dimethyltransferase |
| | P_0199 | 50S ribosomal protein L35 |
| | P_0615 | Uncharacterized protein |
| | P_0697 | Uncharacterized glycosyl transferase |
| | P_0132 | Toxin-antitoxin AAA ATPase |
| | P_0908 | Membrane protein insertase |
| | P_0511 | Uncharacterized protein |
| | P_0614 | Nicotinate phosphoribosyltransferase |
| | P_0375 | Uncharacterized protein |
| | P_0344 | Inorganic diphosphatase |
| | P_0131 | Fructose-1,6-bisphosphate aldolase |
| | P_0482 | 30S ribosomal protein S21 |
| | P_0113 | Glycolipid synthase A |
| | P_0409 | Uncharacterized protein |
| | P_0042 | Uncharacterized transcriptional regulator |
| | P_0726 | Glucosamine-6-phosphate deaminase |
| | P_0427 | Phosphate ABC transporter ATP-binding protein |
| | P_0691 | Uncharacterized protein |
| | P_0302 | Flavin reductase |
| | P_0530 | Uncharacterized protein |
| | P_0733 | Phosphopentomutase |
| | P_0228 | Dihydrolipoyl dehydrogenase |

*continued on next page*

Table S2 – *continued from previous page*

| **Module** | **Protein ID** | **Function** |
|---|---|---|
| | P_0524 | 16S rRNA (cytosine(1402)-N(4))-methyltransferase |
| | P_0377 | Ribosome GTPase |
| | P_0732 | Deoxyribose-phosphate aldolase |
| | P_0515 | Cytidine deaminase |
| | P_0429 | Signal recognition particle-docking protein |
| | P_0651 | Adenylate kinase |
| | P_0793 | F0F1 ATP synthase subunit delta |
| | P_0047 | DNA polymerase III subunit gamma and tau |
| | P_0452 | DNA topoisomerase IV subunit B |
| Module 22 | P_0652 | Preprotein translocase subunit |
| | P_0414 | Guanosine-3',5'-bis(diphosphate) 3'-pyrophosphohydrolase |
| | P_0435 | Mannose-6-phosphate isomerase |
| | P_0441 | Cysteine desulfurase |
| | P_0636 | Uncharacterized lipoprotein |
| | P_0391 | Elongation factor P |
| | P_0695 | ATP-dependent DNA helicase |
| | P_0203 | Guanylate kinase |
| | P_0140 | Thymidine kinase |
| | P_0115 | UTP–glucose-1-phosphate uridylyltransferase |
| | P_0479 | Uncharacterized peptidase |
| | P_0047 | DNA polymerase III subunit gamma and tau |
| | P_0885 | tRNA uridine(34) 5-carboxymethylaminomethyl synthesis enzyme |

*continued on next page*

Table S2 – *continued from previous page*

| **Module** | **Protein ID** | **Function** |
|---|---|---|
| | P_0708 | Thiamine ABC transporter substrate-binding protein |
| | P_0285 | Elongation factor 4 |
| | P_0005 | Uncharacterized protein |
| | P_0530 | Uncharacterized protein |
| | P_0773 | Ribonucleoside-diphosphate reductase subunit beta |
| | P_0648 | 50S ribosomal protein L36 |
| | P_0710 | Low specificity hydrolase |
| | P_0202 | 16S rRNA (guanine(966)-N(2))-methyltransferase |
| | P_0415 | Chromosome segregation protein |
| | P_0650 | Type I methionyl aminopeptidase |
| | P_0006 | DNA gyrase subunit B |
| | P_0303 | PolC-type DNA polymerase III |
| | P_0611 | DNA polymerase I |
| | P_0113 | Glycolipid synthase A |
| | P_0544 | Heat-inducible transcription repressor |
| | P_0776 | SsrA-binding protein |
| | P_0109 | Deoxyribonuclease IV |
| | P_0494 | N-acetylmannosamine-6-phosphate 2-epimerase |
| | P_0228 | Dihydrolipoyl dehydrogenase |
| | P_0347 | Cytidylate kinase |
| | P_0247 | Ribosome biogenesis GTP-binding protein |
| | P_0453 | DNA topoisomerase IV subunit A |

*continued on next page*

Table S2 – *continued from previous page*

| **Module** | **Protein ID** | **Function** |
|---|---|---|
| | P_0732 | Deoxyribose-phosphate aldolase |
| | P_0859 | DNA topoisomerase I |
| | P_0537 | UMP kinase |
| | P_0289 | Ribosome-binding factor A |
| | P_0799 | Serine hydroxymethyltransferase |
| | P_0139 | NanoRNAse |
| | P_0872 | Uncharacterized ATPase |
| | P_0229 | Phosphate acetyltransferase |
| | P_0166 | Oligopeptide ABC transporter permease |
| | P_0169 | Oligopeptide ABC transporter substrate-binding protein |
| | P_0689 | Glutamyl-tRNA amidotransferase subunit C |
| | P_0302 | Flavin reductase |
| | P_0668 | 50S ribosomal protein L2 |
| | P_0314 | Uncharacterized ECF transporter S component |
| | P_0791 | F0F1 ATP synthase subunit gamma |
| | P_0001 | Chromosomal replication initiator protein |
| | P_0359 | Ribonuclease Y |
| | P_0614 | Nicotinate phosphoribosyltransferase |
| | P_0411 | Uncharacterized protein |
| | P_0800 | Ribose 5-phosphate isomerase B |
| | P_0054 | Uncharacterized peroxiredoxin |
| | P_0254 | Excinuclease ABC subunit C |
| Module 23 | P_0818 | Diacylglyceryl transferase |
| | P_0107 | Transcription antitermination factor |
| | P_0813 | UDP-glucose 4-epimerase GalE |

*continued on next page*

Table S2 – *continued from previous page*

| Module | Protein ID | Function |
|---|---|---|
| | P_0133 | Toxin-antitoxin serine protease |
| | P_0824 | Excinuclease ABC subunit A |
| | P_0097 | Flap endonuclease and 5' exonuclease |
| | P_0150 | Translation elongation factor G |
| | P_0544 | Heat-inducible transcription repressor |
| | P_0196 | Spermidine/putrescine ABC transporter permease |
| | P_0390 | Methionyl-tRNA formyltransferase |
| | P_0615 | Uncharacterized protein |
| | P_0359 | Ribonuclease Y |
| | P_0872 | Uncharacterized ATPase |
| | P_0906 | Uncharacterized protein |
| | P_0695 | ATP-dependent DNA helicase |
| | P_0428 | Phosphate transport system regulatory protein |
| | P_0143 | Uncharacterized protein |
| | P_0800 | Ribose 5-phosphate isomerase B |
| | P_0197 | Spermidine/putrescine ABC transporter ATP-binding protein |
| | P_0859 | DNA topoisomerase I |
| | P_0117 | Acyl-phosphate glycerol 3-phosphate acyltransferase |
| | P_0168 | Oligopeptide ABC transporter ATP-binding protein |
| | P_0228 | Dihydrolipoyl dehydrogenase |
| | P_0247 | Ribosome biogenesis GTP-binding protein |
| | P_0419 | Phosphate acyltransferase |

*continued on next page*

Table S2 – *continued from previous page*

| Module | Protein ID | Function |
|---|---|---|
| | P_0410 | Degradosome RNA helicase |
| | P_0838 | 23S rRNA (guanosine(2251)-2'-O)-methyltransferase |
| | P_0042 | Uncharacterized transcriptional regulator |
| | P_0438 | Uncharacterized protein |
| | P_0002 | DNA polymerase III subunit beta |
| | P_0505 | Uncharacterized lipoprotein |
| | P_0453 | DNA topoisomerase IV subunit A |
| | P_0415 | Chromosome segregation protein |
| | P_0406 | DNA primase |
| | P_0132 | Toxin-antitoxin AAA ATPase |
| | P_0420 | Fatty acid kinase subunit A |
| | P_0875 | CDP-diacylglycerol–glycerol-3-phosphate 3-phosphatidyltransferase |
| | P_0294 | 30S ribosomal protein S15 |
| | P_0009 | Nucleoside ABC transporter permease |
| | P_0541 | Molecular chaperone |
| | P_0063 | Uncharacterized tRNA dihydrouridine synthase |
| | P_0650 | Type I methionyl aminopeptidase |
| | P_0165 | Oligopeptide ABC transporter permease |
| | P_0775 | Ribonuclease R |
| | P_0380 | Nicotinate (nicotinamide) nucleotide adenylyl-transferase |
| | P_0840 | Antitermination protein |
| | P_0827 | Uncharacterized protein |
| | P_0043 | Uncharacterized methyltransferase |

*continued on next page*

Table S2 – *continued from previous page*

| **Module** | **Protein ID** | **Function** |
|---|---|---|
| | P_0834 | Replicative DNA helicase |
| | P_0316 | Transketolase |
| | P_0200 | Translation initiation factor IF-3 |
| | P_0250 | Uncharacterized protein |
| | P_0201 | Peptide deformylase |
| | P_0138 | Uncharacterized protein |
| | P_0145 | Uncharacterized acetyltransferase |
| | P_0832 | Aminoacyl-tRNA hydrolase |
| | P_0081 | tRNA uridine(34) 5-carboxymethylaminomethyl synthesis GTPase |
| | P_0814 | UDP-galactopyranose mutase |
| | P_0249 | Uncharacterized protein |
| | P_0776 | SsrA-binding protein |
| | P_0651 | Adenylate kinase |
| | P_0791 | F0F1 ATP synthase subunit gamma |
| | P_0065 | Thioredoxin |
| | P_0879 | magnesium-importing ATPase |
| | P_0299 | Uncharacterized protein |
| | P_0441 | Cysteine desulfurase |
| Module 24 | P_0002 | DNA polymerase III subunit beta |
| | P_0281 | Uncharacterized protein |
| | P_0151 | Translation elongation factor Tu |
| | P_0637 | 30S ribosomal protein S9 |
| | P_0664 | 50S ribosomal protein L16 |
| | P_0599 | Uncharacterized protein |
| | P_0733 | Phosphopentomutase |

*continued on next page*

Table S2 – *continued from previous page*

| Module | Protein ID | Function |
|---|---|---|
| | P_0380 | Nicotinate (nicotinamide) nucleotide adenylyl-transferase |
| | P_0833 | 50S ribosomal protein L9 |
| | P_0348 | Ribosome biogenesis GTPase |
| | P_0215 | Pre-16S rRNA nuclease |
| | P_0421 | Uncharacterized protein |
| | P_0114 | Glycolipid synthase B |
| | P_0710 | Low specificity hydrolase |
| | P_0220 | 6-Phosphofructokinase |
| | P_0005 | Uncharacterized protein |
| | P_0108 | Uncharacterized lipoprotein |
| | P_0420 | Fatty acid kinase subunit A |
| | P_0301 | Ribosome assembly cofactor |
| | P_0779 | PTS sugar transporter |
| | P_0433 | Uncharacterized protein |
| | P_0646 | 30S ribosomal protein S11 |
| | P_0686 | Potassium transporter |
| | P_0239 | Uncharacterized protein |
| | P_0364 | tRNA (guanosine(37)-N1)-methyltransferase |
| | P_0611 | DNA polymerase I |
| | P_0601 | Uncharacterized protein |
| | P_0410 | Degradosome RNA helicase |
| | P_0501 | 50S ribosomal protein L21 |
| | P_0651 | Adenylate kinase |
| | P_0418 | Ribonuclease III |
| | P_0065 | Thioredoxin |

*continued on next page*

Table S2 – *continued from previous page*

| Module | Protein ID | Function |
|---|---|---|
| | P_0294 | 30S ribosomal protein S15 |
| | P_0881 | Uncharacterized MFS transporter |
| | P_0549 | Non-canonical purine NTP pyrophosphatase |
| | P_0010 | Nucleoside ABC transporter ATP-binding protein |
| | P_0254 | Excinuclease ABC subunit C |
| | P_0859 | DNA topoisomerase I |
| | P_0372 | Flippase B |
| | P_0541 | Molecular chaperone |
| | P_0831 | Phosphoribosylpyrophosphate synthetase |
| | P_0201 | Peptide deformylase |
| | P_0435 | Mannose-6-phosphate isomerase |
| | P_0499 | 50S ribosomal protein L27 |
| | P_0773 | Ribonucleoside-diphosphate reductase subunit beta |
| | P_0886 | Proton-glutamate symporter |
| | P_0131 | Fructose-1,6-bisphosphate aldolase |
| | P_0127 | Uncharacterized phosphohydrolase |
| | P_0054 | Uncharacterized peroxiredoxin |
| | P_0409 | Uncharacterized protein |
| | P_0872 | Uncharacterized ATPase |
| | P_0544 | Heat-inducible transcription repressor |
| | P_0316 | Transketolase |
| | P_0060 | Uncharacterized protein |
| | P_0799 | Serine hydroxymethyltransferase |
| | P_0614 | Nicotinate phosphoribosyltransferase |

*continued on next page*

Table S2 – *continued from previous page*

| **Module** | **Protein ID** | **Function** |
|---|---|---|
| | P_0412 | Uncharacterized protein |
| | P_0043 | Uncharacterized methyltransferase |
| | P_0813 | UDP-glucose 4-epimerase GalE |
| | P_0361 | 23S rRNA (pseudouridine(1915)-N3)-methyltransferase |
| | P_0132 | Toxin-antitoxin AAA ATPase |
| | P_0692 | Uncharacterized pseudouridine synthase |
| | P_0794 | F0F1 ATP synthase subunit B |
| | P_0117 | Acyl-phosphate glycerol 3-phosphate acyltransferase |
| | P_0824 | Excinuclease ABC subunit A |
| | P_0387 | tRNA 2-thiouridine(34) synthase |
| | P_0365 | 50S ribosomal protein L19 |
| | P_0697 | Uncharacterized glycosyl transferase |
| | P_0427 | Phosphate ABC transporter ATP-binding protein |
| | P_0885 | tRNA uridine(34) 5-carboxymethylaminomethyl synthesis enzyme |
| Module 25 | P_0007 | DNA gyrase subunit A |
| | P_0494 | N-acetylmannosamine-6-phosphate 2-epimerase |
| | P_0642 | ECF transporter ATPase |
| | P_0907 | Low specificity hydrolase |
| | P_0445 | Glucose-6-phosphate isomerase |
| | P_0108 | Uncharacterized lipoprotein |

*continued on next page*

Table S2 – *continued from previous page*

| **Module** | **Protein ID** | **Function** |
|---|---|---|
| | P_0004 | 16S rRNA (adenine(1518)-N(6)/adenine(1519)-N(6))- dimethyltransferase |
| | P_0537 | UMP kinase |
| | P_0886 | Proton-glutamate symporter |
| | P_0779 | PTS sugar transporter |
| | P_0814 | UDP-galactopyranose mutase |
| | P_0389 | Uncharacterized protein |
| | P_0611 | DNA polymerase I |
| | P_0372 | Flippase B |
| | P_0501 | 50S ribosomal protein L21 |
| | P_0218 | Glycerol kinase |
| | P_0729 | Phosphoglycerate mutase (2,3-diphosphoglycerate-independent) |
| | P_0005 | Uncharacterized protein |
| | P_0420 | Fatty acid kinase subunit A |
| | P_0165 | Oligopeptide ABC transporter permease |
| | P_0614 | Nicotinate phosphoribosyltransferase |
| | P_0824 | Excinuclease ABC subunit A |
| | P_0299 | Uncharacterized protein |
| | P_0516 | Uncharacterized protein |
| | P_0876 | Uncharacterized amino acid permease |
| | P_0410 | Degradosome RNA helicase |
| | P_0006 | DNA gyrase subunit B |
| | P_0081 | tRNA uridine(34) 5-carboxymethylaminomethyl synthesis GTPase |
| | P_0392 | Uncharacterized protein |

*continued on next page*

Table S2 – *continued from previous page*

| Module | Protein ID | Function |
|---|---|---|
| | P_0001 | Chromosomal replication initiator protein |
| | P_0796 | F0F1 ATP synthase subunit A |
| | P_0435 | Mannose-6-phosphate isomerase |
| | P_0169 | Oligopeptide ABC transporter substrate-binding protein |
| | P_0688 | Glutamyl-tRNA amidotransferase subunit A |
| | P_0378 | NAD(+) synthase |
| | P_0615 | Uncharacterized protein |
| | P_0382 | Deoxynucleoside kinase |
| | P_0412 | Uncharacterized protein |
| | P_0686 | Potassium transporter |
| | P_0792 | F0F1 ATP synthase subunit alpha |
| | P_0139 | NanoRNAse |
| | P_0799 | Serine hydroxymethyltransferase |
| | P_0689 | Glutamyl-tRNA amidotransferase subunit C |
| | P_0859 | DNA topoisomerase I |
| | P_0195 | Spermidine/putrescine ABC transporter permease |
| | P_0294 | 30S ribosomal protein S15 |
| | P_0371 | Flippase A |
| | P_0687 | Glutamyl-tRNA amidotransferase subunit B |
| | P_0344 | Inorganic diphosphatase |
| | P_0885 | tRNA uridine(34) 5-carboxymethylaminomethyl synthesis enzyme |
| | P_0262 | Ribulose-phosphate 3-epimerase |
| | P_0316 | Transketolase |

*continued on next page*

Table S2 – *continued from previous page*

| Module | Protein ID | Function |
|---|---|---|
| | P_0600 | Ribonuclease J |
| | P_0202 | 16S rRNA (guanine(966)-N(2))-methyltransferase |
| | P_0726 | Glucosamine-6-phosphate deaminase |
| | P_0377 | Ribosome GTPase |
| | P_0691 | Uncharacterized protein |
| | P_0117 | Acyl-phosphate glycerol 3-phosphate acyltransferase |
| | P_0650 | Type I methionyl aminopeptidase |
| | P_0639 | Uncharacterized efflux ABC transporter permease |
| | P_0645 | DNA-directed RNA polymerase subunit alpha |
| | P_0250 | Uncharacterized protein |
| | P_0651 | Adenylate kinase |
| | P_0835 | Uncharacterized lipoprotein |
| | P_0427 | Phosphate ABC transporter ATP-binding protein |
| | P_0708 | Thiamine ABC transporter substrate-binding protein |
| | P_0366 | L16-binding dependent 50S subunit-maturation GTPase |
| | P_0452 | DNA topoisomerase IV subunit B |
| | P_0353 | Cell division DivIVA/GpsB protein |
| | P_0391 | Elongation factor P |
| | P_0303 | PolC-type DNA polymerase III |
| Module 26 | P_0137 | 50S ribosomal protein L31 |

*continued on next page*

Table S2 – *continued from previous page*

| Module | Protein ID | Function |
|---|---|---|
| | P_0611 | DNA polymerase I |
| | P_0108 | Uncharacterized lipoprotein |
| | P_0372 | Flippase B |
| | P_0814 | UDP-galactopyranose mutase |
| | P_0692 | Uncharacterized pseudouridine synthase |
| | P_0140 | Thymidine kinase |
| | P_0879 | magnesium-importing ATPase |
| | P_0329 | Uncharacterized pseudouridine synthase |
| | P_0387 | tRNA 2-thiouridine(34) synthase |
| | P_0772 | Ribonucleotide reductase assembly protein |
| | P_0906 | Uncharacterized protein |
| | P_0600 | Ribonuclease J |
| | P_0908 | Membrane protein insertase |
| | P_0257 | RNase J family beta-CASP ribonuclease |
| | P_0271 | tRNA N6-adenosine(37)-N6-threonylcarbamoyltransferase complex dimerization subunit |
| | P_0415 | Chromosome segregation protein |
| | P_0026 | Single-stranded DNA-binding protein |
| | P_0004 | 16S rRNA (adenine(1518)-N(6)/adenine(1519)-N(6))- dimethyltransferase |
| | P_0796 | F0F1 ATP synthase subunit A |
| | P_0371 | Flippase A |
| | P_0294 | 30S ribosomal protein S15 |
| | P_0824 | Excinuclease ABC subunit A |
| | P_0549 | Non-canonical purine NTP pyrophosphatase |

*continued on next page*

Table S2 – *continued from previous page*

| Module | Protein ID | Function |
|---|---|---|
| | P_0859 | DNA topoisomerase I |
| | P_0390 | Methionyl-tRNA formyltransferase |
| | P_0195 | Spermidine/putrescine ABC transporter permease |
| | P_0109 | Deoxyribonuclease IV |
| | P_0009 | Nucleoside ABC transporter permease |
| | P_0431 | Uncharacterized metallophosphoesterase |
| | P_0005 | Uncharacterized protein |
| | P_0800 | Ribose 5-phosphate isomerase B |
| | P_0344 | Inorganic diphosphatase |
| | P_0524 | 16S rRNA (cytosine(1402)-N(4))-methyltransferase |
| | P_0453 | DNA topoisomerase IV subunit A |
| | P_0775 | Ribonuclease R |
| | P_0689 | Glutamyl-tRNA amidotransferase subunit C |
| | P_0512 | Acyl-phosphate glycerol 3-phosphate acyltransferase |
| | P_0697 | Uncharacterized glycosyl transferase |
| | P_0042 | Uncharacterized transcriptional regulator |
| | P_0054 | Uncharacterized peroxiredoxin |
| | P_0283 | Double-stranded RNA binding RNase HI |
| | P_0610 | DNA-formamidopyrimidine glycosylase |
| | P_0302 | Flavin reductase |
| | P_0834 | Replicative DNA helicase |
| | P_0505 | Uncharacterized lipoprotein |
| | P_0420 | Fatty acid kinase subunit A |

*continued on next page*

Table S2 – *continued from previous page*

| **Module** | **Protein ID** | **Function** |
|---|---|---|
| | P_0726 | Glucosamine-6-phosphate deaminase |
| | P_0011 | Nucleoside ABC transporter substrate-binding protein |
| | P_0615 | Uncharacterized protein |
| | P_0361 | 23S rRNA (pseudouridine(1915)-N3)-methyltransferase |
| | P_0548 | tRNA (cytidine(34)-2'-O)-methyltransferase |
| | P_0113 | Glycolipid synthase A |
| | P_0517 | Uncharacterized RNA pseudouridine synthase |
| | P_0366 | L16-binding dependent 50S subunit-maturation GTPase |
| | P_0819 | Thioredoxin-disulfide reductase |
| | P_0650 | Type I methionyl aminopeptidase |
| | P_0688 | Glutamyl-tRNA amidotransferase subunit A |
| | P_0412 | Uncharacterized protein |
| Module 27 | P_0538 | Uncharacterized protein |
| | P_0044 | DNA polymerase III subunit delta |
| | P_0875 | CDP-diacylglycerol–glycerol-3-phosphate 3-phosphatidyltransferase |
| | P_0827 | Uncharacterized protein |
| | P_0697 | Uncharacterized glycosyl transferase |
| | P_0879 | magnesium-importing ATPase |
| | P_0132 | Toxin-antitoxin AAA ATPase |
| | P_0813 | UDP-glucose 4-epimerase GalE |
| | P_0652 | Preprotein translocase subunit |

*continued on next page*

Table S2 – *continued from previous page*

| Module | Protein ID | Function |
|---|---|---|
| | P_0202 | 16S rRNA (guanine(966)-N(2))-methyltransferase |
| | P_0689 | Glutamyl-tRNA amidotransferase subunit C |
| | P_0215 | Pre-16S rRNA nuclease |
| | P_0247 | Ribosome biogenesis GTP-binding protein |
| | P_0139 | NanoRNAse |
| | P_0127 | Uncharacterized phosphohydrolase |
| | P_0650 | Type I methionyl aminopeptidase |
| | P_0010 | Nucleoside ABC transporter ATP-binding protein |
| | P_0294 | 30S ribosomal protein S15 |
| | P_0605 | Uncharacterized protein |
| | P_0011 | Nucleoside ABC transporter substrate-binding protein |
| | P_0428 | Phosphate transport system regulatory protein |
| | P_0695 | ATP-dependent DNA helicase |
| | P_0009 | Nucleoside ABC transporter permease |
| | P_0380 | Nicotinate (nicotinamide) nucleotide adenylyltransferase |
| | P_0832 | Aminoacyl-tRNA hydrolase |
| | P_0499 | 50S ribosomal protein L27 |
| | P_0686 | Potassium transporter |
| | P_0042 | Uncharacterized transcriptional regulator |
| | P_0617 | Fatty acid binding protein |
| | P_0359 | Ribonuclease Y |
| | P_0779 | PTS sugar transporter |

*continued on next page*

Table S2 – *continued from previous page*

| **Module** | **Protein ID** | **Function** |
|---|---|---|
| | P_0029 | FMN reductase |
| | P_0438 | Uncharacterized protein |
| | P_0195 | Spermidine/putrescine ABC transporter permease |
| | P_0145 | Uncharacterized acetyltransferase |
| | P_0201 | Peptide deformylase |
| | P_0887 | Coenzyme A Disulfide Reductase |
| | P_0453 | DNA topoisomerase IV subunit A |
| | P_0146 | Uncharacterized protein |
| | P_0799 | Serine hydroxymethyltransferase |
| | P_0794 | F0F1 ATP synthase subunit B |
| | P_0410 | Degradosome RNA helicase |
| | P_0289 | Ribosome-binding factor A |
| | P_0515 | Cytidine deaminase |
| | P_0199 | 50S ribosomal protein L35 |
| | P_0262 | Ribulose-phosphate 3-epimerase |
| | P_0200 | Translation initiation factor IF-3 |
| | P_0517 | Uncharacterized RNA pseudouridine synthase |
| | P_0407 | RNA polymerase subunit sigma |
| | P_0141 | Peptide chain release factor 1 |
| Module 28 | P_0144 | tRNA (adenosine(37)-N6)-threonylcarbamoyltransferase complex transferase subunit |
| | P_0684 | Bifunctional 5,10-methylene-tetrahydrofolate dehydrogenase/5,10-methylene-tetrahydrofolate cyclohydrolase |

*continued on next page*

Table S2 – *continued from previous page*

| Module | Protein ID | Function |
|---|---|---|
| | P_0482 | 30S ribosomal protein S21 |
| | P_0326 | Uncharacterized protein |
| | P_0657 | 30S ribosomal protein S8 |
| | P_0060 | Uncharacterized protein |
| | P_0390 | Methionyl-tRNA formyltransferase |
| | P_0303 | PolC-type DNA polymerase III |
| | P_0776 | SsrA-binding protein |
| | P_0800 | Ribose 5-phosphate isomerase B |
| | P_0823 | Dihydrofolate synthase |
| | P_0382 | Deoxynucleoside kinase |
| | P_0445 | Glucose-6-phosphate isomerase |
| | P_0918 | Imidazoleglycerol-phosphate dehydratase |
| | P_0452 | DNA topoisomerase IV subunit B |
| | P_0002 | DNA polymerase III subunit beta |
| | P_0109 | Deoxyribonuclease IV |
| | P_0835 | Uncharacterized lipoprotein |
| | P_0771 | Ribonucleotide-diphosphate reductase subunit alpha |
| | P_0439 | Uncharacterized lipoprotein |
| | P_0133 | Toxin-antitoxin serine protease |
| | P_0516 | Uncharacterized protein |
| | P_0813 | UDP-glucose 4-epimerase GalE |
| | P_0414 | Guanosine-3',5'-bis(diphosphate) 3'-pyrophosphohydrolase |
| | P_0033 | Uncharacterized protein |
| | P_0643 | ECF transporter ATPase |

*continued on next page*

Table S2 – *continued from previous page*

| Module | Protein ID | Function |
|---|---|---|
| | P_0359 | Ribonuclease Y |
| | P_0353 | Cell division DivIVA/GpsB protein |
| | P_0081 | tRNA uridine(34) 5-carboxymethylaminomethyl synthesis GTPase |
| | P_0799 | Serine hydroxymethyltransferase |
| | P_0604 | Uncharacterized protein |
| | P_0431 | Uncharacterized metallophosphoesterase |
| | P_0609 | Chromosome replication initiation protein |
| | P_0202 | 16S rRNA (guanine(966)-N(2))-methyltransferase |
| | P_0347 | Cytidylate kinase |
| | P_0440 | Uncharacterized lipoprotein |
| | P_0127 | Uncharacterized phosphohydrolase |
| | P_0140 | Thymidine kinase |
| | P_0653 | 50S ribosomal protein L15 |
| | P_0641 | ECF transporter T component |
| | P_0608 | Primosomal protein |
| | P_0426 | Phosphate ABC transporter permease |
| | P_0537 | UMP kinase |
| | P_0789 | F0F1 ATP synthase subunit delta/epsilon |
| | P_0779 | PTS sugar transporter |
| | P_0289 | Ribosome-binding factor A |
| | P_0875 | CDP-diacylglycerol–glycerol-3-phosphate 3-phosphatidyltransferase |
| | P_0687 | Glutamyl-tRNA amidotransferase subunit B |
| | P_0825 | Excinuclease ABC subunit B |

*continued on next page*

Table S2 – *continued from previous page*

| Module | Protein ID | Function |
|---|---|---|
| | P_0201 | Peptide deformylase |
| | P_0695 | ATP-dependent DNA helicase |
| | P_0870 | Uncharacterized C4-dicarboxylate ABC transporter |
| | P_0409 | Uncharacterized protein |
| | P_0832 | Aminoacyl-tRNA hydrolase |
| | P_0262 | Ribulose-phosphate 3-epimerase |
| | P_0165 | Oligopeptide ABC transporter permease |
| | P_0139 | NanoRNAse |
| | P_0129 | CTP synthase |
| | P_0690 | DNA ligase (NAD(+)) |
| | P_0617 | Fatty acid binding protein |
| | P_0434 | 23S rRNA (uridine(1939)-m5)-methyltransferase |
| | P_0418 | Ribonuclease III |
| | P_0329 | Uncharacterized pseudouridine synthase |
| | P_0517 | Uncharacterized RNA pseudouridine synthase |
| | P_0257 | RNase J family beta-CASP ribonuclease |
| | P_0271 | tRNA N6-adenosine(37)-N6-threonylcarbamoyltransferase complex dimerization subunit |
| | P_0371 | Flippase A |
| | P_0441 | Cysteine desulfurase |
| | P_0794 | F0F1 ATP synthase subunit B |
| | P_0730 | Uncharacterized protein |
| | P_0094 | Uncharacterized protein |

*continued on next page*

Table S2 – *continued from previous page*

| **Module** | **Protein ID** | **Function** |
|---|---|---|
| | P_0707 | Thiamine ABC transporter ATP-binding protein |
| | P_0885 | tRNA uridine(34) 5-carboxymethylaminomethyl synthesis enzyme |
| | P_0117 | Acyl-phosphate glycerol 3-phosphate acyltransferase |
| | P_0614 | Nicotinate phosphoribosyltransferase |
| | P_0429 | Signal recognition particle-docking protein |
| | P_0040 | tRNA lysidine(34) synthetase |
| | P_0113 | Glycolipid synthase A |
| | P_0422 | 50S ribosomal protein L28 |
| | P_0710 | Low specificity hydrolase |
| | P_0010 | Nucleoside ABC transporter ATP-binding protein |
| | P_0239 | Uncharacterized protein |
| | P_0728 | Low specificity hydrolase |
| | P_0824 | Excinuclease ABC subunit A |
| | P_0530 | Uncharacterized protein |
| | P_0543 | Nucleotide exchange factor |
| | P_0652 | Preprotein translocase subunit |
| Module 29 | P_0346 | Uncharacterized protein |
| | P_0501 | 50S ribosomal protein L21 |
| | P_0653 | 50S ribosomal protein L15 |
| | P_0109 | Deoxyribonuclease IV |
| | P_0835 | Uncharacterized lipoprotein |
| | P_0819 | Thioredoxin-disulfide reductase |

*continued on next page*

Table S2 – *continued from previous page*

| Module | Protein ID | Function |
|---|---|---|
| | P_0402 | rRNA maturation RNase |
| | P_0044 | DNA polymerase III subunit delta |
| | P_0132 | Toxin-antitoxin AAA ATPase |
| | P_0526 | 50S ribosomal protein L32 |
| | P_0414 | Guanosine-3',5'-bis(diphosphate) 3'-pyrophosphohydrolase |
| | P_0601 | Uncharacterized protein |
| | P_0727 | Triose-phosphate isomerase |
| | P_0114 | Glycolipid synthase B |
| | P_0361 | 23S rRNA (pseudouridine(1915)-N3)-methyltransferase |
| | P_0642 | ECF transporter ATPase |
| | P_0042 | Uncharacterized transcriptional regulator |
| | P_0127 | Uncharacterized phosphohydrolase |
| | P_0081 | tRNA uridine(34) 5-carboxymethylaminomethyl synthesis GTPase |
| | P_0195 | Spermidine/putrescine ABC transporter permease |
| | P_0800 | Ribose 5-phosphate isomerase B |
| | P_0419 | Phosphate acyltransferase |
| | P_0438 | Uncharacterized protein |
| | P_0451 | Glyceraldehyde-3-phosphate dehydrogenase |
| | P_0203 | Guanylate kinase |
| | P_0362 | 30S ribosomal protein S16 |
| | P_0874 | 16S rRNA (guanine(527)-N(7))-methyltransferase |

*continued on next page*

Table S2 – *continued from previous page*

| Module | Protein ID | Function |
|---|---|---|
| | P_0908 | Membrane protein insertase |
| | P_0145 | Uncharacterized acetyltransferase |
| | P_0821 | HPr(Ser) kinase/phosphatase |
| | P_0010 | Nucleoside ABC transporter ATP-binding protein |
| | P_0139 | NanoRNAse |
| | P_0661 | 50S ribosomal protein L14 |
| | P_0441 | Cysteine desulfurase |
| | P_0887 | Coenzyme A Disulfide Reductase |
| | P_0648 | 50S ribosomal protein L36 |
| | P_0375 | Uncharacterized protein |
| | P_0240 | tRNA 4-thiouridine(8) synthase |
| | P_0512 | Acyl-phosphate glycerol 3-phosphate acyltransferase |
| | P_0116 | Uncharacterized protein |
| | P_0054 | Uncharacterized peroxiredoxin |
| | P_0814 | UDP-galactopyranose mutase |
| | P_0775 | Ribonuclease R |
| | P_0289 | Ribosome-binding factor A |
| | P_0733 | Phosphopentomutase |
| | P_0338 | Uncharacterized lipoprotein |
| | P_0218 | Glycerol kinase |
| | P_0452 | DNA topoisomerase IV subunit B |
| | P_0428 | Phosphate transport system regulatory protein |
| | P_0117 | Acyl-phosphate glycerol 3-phosphate acyltransferase |

*continued on next page*

Table S2 – *continued from previous page*

| Module | Protein ID | Function |
|---|---|---|
| | P_0371 | Flippase A |
| | P_0794 | F0F1 ATP synthase subunit B |
| | P_0832 | Aminoacyl-tRNA hydrolase |
| | P_0517 | Uncharacterized RNA pseudouridine synthase |
| | P_0011 | Nucleoside ABC transporter substrate-binding protein |
| | P_0201 | Peptide deformylase |
| | P_0138 | Uncharacterized protein |
| | P_0250 | Uncharacterized protein |
| | P_0314 | Uncharacterized ECF transporter S component |
| | P_0229 | Phosphate acetyltransferase |
| | P_0617 | Fatty acid binding protein |
| | P_0329 | Uncharacterized pseudouridine synthase |
| Module 30 | P_0402 | rRNA maturation RNase |
| | P_0004 | 16S rRNA (adenine(1518)-N(6)/adenine(1519)-N(6))- dimethyltransferase |
| | P_0451 | Glyceraldehyde-3-phosphate dehydrogenase |
| | P_0147 | Cardiolipin synthase |
| | P_0365 | 50S ribosomal protein L19 |
| | P_0079 | tRNA (adenosine(37)-N6)-threonylcarbamoyltransferase complex transferase subunit D |
| | P_0328 | Chromosome segregation protein B |
| | P_0707 | Thiamine ABC transporter ATP-binding protein |
| | P_0215 | Pre-16S rRNA nuclease |

*continued on next page*

Table S2 – *continued from previous page*

| Module | Protein ID | Function |
|---|---|---|
| | P_0388 | Uncharacterized protein |
| | P_0500 | Maturation protease for ribosomal protein L27 |
| | P_0218 | Glycerol kinase |
| | P_0229 | Phosphate acetyltransferase |
| | P_0831 | Phosphoribosylpyrophosphate synthetase |
| | P_0422 | 50S ribosomal protein L28 |
| | P_0805 | Uncharacterized protein |
| | P_0167 | Oligopeptide ABC transporter ATP-binding protein |
| | P_0029 | FMN reductase |
| | P_0285 | Elongation factor 4 |
| | P_0065 | Thioredoxin |
| | P_0807 | 50S ribosomal protein L10 |
| | P_0819 | Thioredoxin-disulfide reductase |
| | P_0330 | Deoxyguanosine kinase |
| | P_0197 | Spermidine/putrescine ABC transporter ATP-binding protein |
| | P_0517 | Uncharacterized RNA pseudouridine synthase |
| | P_0434 | 23S rRNA (uridine(1939)-m5)-methyltransferase |
| | P_0796 | F0F1 ATP synthase subunit A |
| | P_0840 | Antitermination protein |
| | P_0143 | Uncharacterized protein |
| | P_0605 | Uncharacterized protein |
| | P_0838 | 23S rRNA (guanosine(2251)-2'-O)-methyltransferase |

*continued on next page*

Table S2 – *continued from previous page*

| Module | Protein ID | Function |
| --- | --- | --- |
| | P_0726 | Glucosamine-6-phosphate deaminase |
| | P_0412 | Uncharacterized protein |
| | P_0790 | F0F1 ATP synthase subunit beta |
| | P_0823 | Dihydrofolate synthase |
| | P_0776 | SsrA-binding protein |
| | P_0234 | PTS glucose transporter subunit IIA |
| | P_0203 | Guanylate kinase |
| | P_0876 | Uncharacterized amino acid permease |
| | P_0377 | Ribosome GTPase |
| | P_0511 | Uncharacterized protein |
| | P_0007 | DNA gyrase subunit A |
| | P_0247 | Ribosome biogenesis GTP-binding protein |
| | P_0427 | Phosphate ABC transporter ATP-binding protein |
| | P_0400 | Uncharacterized protease |
| | P_0353 | Cell division DivIVA/GpsB protein |
| | P_0501 | 50S ribosomal protein L21 |
| | P_0366 | L16-binding dependent 50S subunit-maturation GTPase |
| | P_0392 | Uncharacterized protein |
| | P_0438 | Uncharacterized protein |
| | P_0548 | tRNA (cytidine(34)-2'-O)-methyltransferase |
| | P_0419 | Phosphate acyltransferase |
| | P_0728 | Low specificity hydrolase |
| | P_0044 | DNA polymerase III subunit delta |
| | P_0695 | ATP-dependent DNA helicase |

*continued on next page*

Table S2 – *continued from previous page*

| **Module** | **Protein ID** | **Function** |
|---|---|---|
| | P_0289 | Ribosome-binding factor A |
| | P_0650 | Type I methionyl aminopeptidase |
| | P_0338 | Uncharacterized lipoprotein |
| | P_0387 | tRNA 2-thiouridine(34) synthase |
| | P_0651 | Adenylate kinase |
| | P_0515 | Cytidine deaminase |
| | P_0885 | tRNA uridine(34) 5-carboxymethylaminomethyl synthesis enzyme |
| | P_0046 | Recombination protein |
| | P_0303 | PolC-type DNA polymerase III |
| | P_0779 | PTS sugar transporter |
| | P_0009 | Nucleoside ABC transporter permease |
| Module 31 | P_0792 | F0F1 ATP synthase subunit alpha |
| | P_0378 | NAD(+) synthase |
| | P_0419 | Phosphate acyltransferase |
| | P_0407 | RNA polymerase subunit sigma |
| | P_0859 | DNA topoisomerase I |
| | P_0687 | Glutamyl-tRNA amidotransferase subunit B |
| | P_0779 | PTS sugar transporter |
| | P_0254 | Excinuclease ABC subunit C |
| | P_0611 | DNA polymerase I |
| | P_0126 | Glutamate–tRNA ligase |
| | P_0505 | Uncharacterized lipoprotein |
| | P_0143 | Uncharacterized protein |
| | P_0010 | Nucleoside ABC transporter ATP-binding protein |

*continued on next page*

Table S2 – *continued from previous page*

| Module | Protein ID | Function |
|---|---|---|
| | P_0199 | 50S ribosomal protein L35 |
| | P_0415 | Chromosome segregation protein |
| | P_0145 | Uncharacterized acetyltransferase |
| | P_0203 | Guanylate kinase |
| | P_0427 | Phosphate ABC transporter ATP-binding protein |
| | P_0501 | 50S ribosomal protein L21 |
| | P_0688 | Glutamyl-tRNA amidotransferase subunit A |
| | P_0799 | Serine hydroxymethyltransferase |
| | P_0872 | Uncharacterized ATPase |
| | P_0201 | Peptide deformylase |
| | P_0544 | Heat-inducible transcription repressor |
| | P_0344 | Inorganic diphosphatase |
| | P_0732 | Deoxyribose-phosphate aldolase |
| | P_0326 | Uncharacterized protein |
| | P_0512 | Acyl-phosphate glycerol 3-phosphate acyltransferase |
| | P_0365 | 50S ribosomal protein L19 |
| | P_0005 | Uncharacterized protein |
| | P_0814 | UDP-galactopyranose mutase |
| | P_0689 | Glutamyl-tRNA amidotransferase subunit C |
| | P_0500 | Maturation protease for ribosomal protein L27 |
| | P_0726 | Glucosamine-6-phosphate deaminase |
| | P_0047 | DNA polymerase III subunit gamma and tau |
| | P_0372 | Flippase B |
| | P_0448 | Uncharacterized rRNA methyltransferase |

*continued on next page*

Table S2 – *continued from previous page*

| **Module** | **Protein ID** | **Function** |
|---|---|---|
| | P_0834 | Replicative DNA helicase |
| | P_0422 | 50S ribosomal protein L28 |
| | P_0392 | Uncharacterized protein |
| | P_0234 | PTS glucose transporter subunit IIA |
| | P_0696 | Uncharacterized transporter |
| | P_0789 | F0F1 ATP synthase subunit delta/epsilon |
| | P_0139 | NanoRNAse |
| | P_0708 | Thiamine ABC transporter substrate-binding protein |
| Module 32 | P_0027 | 30S ribosomal protein S6 |
| | P_0729 | Phosphoglycerate mutase (2,3-diphosphoglycerate-independent) |
| | P_0133 | Toxin-antitoxin serine protease |
| | P_0932 | 50S ribosomal protein L33 |
| | P_0144 | tRNA (adenosine(37)-N6)-threonylcarbamoyltransferase complex transferase subunit |
| | P_0079 | tRNA (adenosine(37)-N6)-threonylcarbamoyltransferase complex transferase subunit D |
| | P_0819 | Thioredoxin-disulfide reductase |
| | P_0646 | 30S ribosomal protein S11 |
| | P_0516 | Uncharacterized protein |
| | P_0504 | 16S rRNA (cytidine(1402)-2'-O)-methyltransferase |
| | P_0250 | Uncharacterized protein |

*continued on next page*

Table S2 – *continued from previous page*

| **Module** | **Protein ID** | **Function** |
|---|---|---|
| | P_0303 | PolC-type DNA polymerase III |
| | P_0114 | Glycolipid synthase B |
| | P_0686 | Potassium transporter |
| | P_0389 | Uncharacterized protein |
| | P_0362 | 30S ribosomal protein S16 |
| | P_0870 | Uncharacterized C4-dicarboxylate ABC transporter |
| | P_0410 | Degradosome RNA helicase |
| | P_0390 | Methionyl-tRNA formyltransferase |
| | P_0776 | SsrA-binding protein |
| | P_0831 | Phosphoribosylpyrophosphate synthetase |
| | P_0537 | UMP kinase |
| | P_0813 | UDP-glucose 4-epimerase GalE |
| | P_0726 | Glucosamine-6-phosphate deaminase |
| | P_0684 | Bifunctional 5,10-methylene-tetrahydrofolate dehydrogenase/5,10-methylene-tetrahydrofolate cyclohydrolase |
| | P_0373 | Uncharacterized protein |
| | P_0218 | Glycerol kinase |
| | P_0429 | Signal recognition particle-docking protein |
| | P_0215 | Pre-16S rRNA nuclease |
| | P_0805 | Uncharacterized protein |
| | P_0002 | DNA polymerase III subunit beta |
| | P_0794 | F0F1 ATP synthase subunit B |
| | P_0438 | Uncharacterized protein |

*continued on next page*

Table S2 – *continued from previous page*

| Module | Protein ID | Function |
|---|---|---|
| | P_0434 | 23S rRNA (uridine(1939)-m5)-methyltransferase |
| | P_0479 | Uncharacterized peptidase |
| | P_0435 | Mannose-6-phosphate isomerase |
| | P_0360 | Signal recognition particle protein |
| | P_0406 | DNA primase |
| | P_0229 | Phosphate acetyltransferase |
| | P_0453 | DNA topoisomerase IV subunit A |
| | P_0824 | Excinuclease ABC subunit A |
| | P_0538 | Uncharacterized protein |
| | P_0814 | UDP-galactopyranose mutase |
| | P_0885 | tRNA uridine(34) 5-carboxymethylaminomethyl synthesis enzyme |
| | P_0600 | Ribonuclease J |
| | P_0876 | Uncharacterized amino acid permease |
| | P_0047 | DNA polymerase III subunit gamma and tau |
| | P_0375 | Uncharacterized protein |
| | P_0908 | Membrane protein insertase |
| | P_0692 | Uncharacterized pseudouridine synthase |
| | P_0366 | L16-binding dependent 50S subunit-maturation GTPase |
| | P_0039 | Membrane anchored AAA+ protease |
| | P_0689 | Glutamyl-tRNA amidotransferase subunit C |
| | P_0137 | 50S ribosomal protein L31 |
| | P_0140 | Thymidine kinase |
| | P_0138 | Uncharacterized protein |

*continued on next page*

Table S2 – *continued from previous page*

| Module | Protein ID | Function |
|---|---|---|
| | P_0615 | Uncharacterized protein |
| | P_0304 | Phosphatidate cytidylyltransferase |
| | P_0641 | ECF transporter T component |
| | P_0202 | 16S rRNA (guanine(966)-N(2))-methyltransferase |
| | P_0617 | Fatty acid binding protein |
| | P_0030 | Uncharacterized ABC transporter ATP-binding protein |
| | P_0271 | tRNA N6-adenosine(37)-N6-threonylcarbamoyltransferase complex dimerization subunit |
| | P_0887 | Coenzyme A Disulfide Reductase |
| | P_0503 | Uncharacterized protein |
| | P_0097 | Flap endonuclease and 5' exonuclease |
| | P_0517 | Uncharacterized RNA pseudouridine synthase |
| | P_0874 | 16S rRNA (guanine(527)-N(7))-methyltransferase |
| | P_0833 | 50S ribosomal protein L9 |
| | P_0063 | Uncharacterized tRNA dihydrouridine synthase |
| | P_0859 | DNA topoisomerase I |
| | P_0775 | Ribonuclease R |
| | P_0800 | Ribose 5-phosphate isomerase B |
| Module 33 | P_0654 | 30S ribosomal protein S5 |
| | P_0657 | 30S ribosomal protein S8 |
| | P_0203 | Guanylate kinase |
| | P_0601 | Uncharacterized protein |

*continued on next page*

Table S2 – *continued from previous page*

| **Module** | **Protein ID** | **Function** |
|---|---|---|
| | P_0840 | Antitermination protein |
| | P_0047 | DNA polymerase III subunit gamma and tau |
| | P_0440 | Uncharacterized lipoprotein |
| | P_0011 | Nucleoside ABC transporter substrate-binding protein |
| | P_0411 | Uncharacterized protein |
| | P_0346 | Uncharacterized protein |
| | P_0615 | Uncharacterized protein |
| | P_0729 | Phosphoglycerate mutase (2,3-diphosphoglycerate-independent) |
| | P_0874 | 16S rRNA (guanine(527)-N(7))-methyltransferase |
| | P_0138 | Uncharacterized protein |
| | P_0726 | Glucosamine-6-phosphate deaminase |
| | P_0817 | Uncharacterized DNA-binding protein |
| | P_0390 | Methionyl-tRNA formyltransferase |
| | P_0421 | Uncharacterized protein |
| | P_0608 | Primosomal protein |
| | P_0792 | F0F1 ATP synthase subunit alpha |
| | P_0600 | Ribonuclease J |
| | P_0688 | Glutamyl-tRNA amidotransferase subunit A |
| | P_0526 | 50S ribosomal protein L32 |
| | P_0687 | Glutamyl-tRNA amidotransferase subunit B |
| | P_0549 | Non-canonical purine NTP pyrophosphatase |
| | P_0523 | Cell division protein |

*continued on next page*

Table S2 – *continued from previous page*

| Module | Protein ID | Function |
|---|---|---|
| | P_0684 | Bifunctional 5,10-methylene-tetrahydrofolate dehydrogenase/5,10-methylene-tetrahydrofolate cyclohydrolase |
| | P_0708 | Thiamine ABC transporter substrate-binding protein |
| | P_0285 | Elongation factor 4 |
| | P_0859 | DNA topoisomerase I |
| | P_0451 | Glyceraldehyde-3-phosphate dehydrogenase |
| | P_0592 | Uncharacterized protein |
| | P_0733 | Phosphopentomutase |
| | P_0438 | Uncharacterized protein |
| | P_0108 | Uncharacterized lipoprotein |
| | P_0610 | DNA-formamidopyrimidine glycosylase |
| | P_0002 | DNA polymerase III subunit beta |
| | P_0344 | Inorganic diphosphatase |
| | P_0253 | Transcription elongation factor |
| | P_0544 | Heat-inducible transcription repressor |
| | P_0389 | Uncharacterized protein |
| | P_0611 | DNA polymerase I |
| | P_0372 | Flippase B |
| | P_0247 | Ribosome biogenesis GTP-binding protein |
| | P_0195 | Spermidine/putrescine ABC transporter permease |
| | P_0033 | Uncharacterized protein |
| | P_0452 | DNA topoisomerase IV subunit B |
| | P_0140 | Thymidine kinase |

*continued on next page*

Table S2 – *continued from previous page*

| Module | Protein ID | Function |
|---|---|---|
| | P_0139 | NanoRNAse |
| | P_0791 | F0F1 ATP synthase subunit gamma |
| | P_0107 | Transcription antitermination factor |
| | P_0885 | tRNA uridine(34) 5-carboxymethylaminomethyl synthesis enzyme |
| | P_0501 | 50S ribosomal protein L21 |
| | P_0881 | Uncharacterized MFS transporter |
| | P_0006 | DNA gyrase subunit B |
| | P_0834 | Replicative DNA helicase |
| | P_0362 | 30S ribosomal protein S16 |
| | P_0537 | UMP kinase |
| | P_0202 | 16S rRNA (guanine(966)-N(2))-methyltransferase |
| | P_0289 | Ribosome-binding factor A |
| | P_0240 | tRNA 4-thiouridine(8) synthase |
| | P_0426 | Phosphate ABC transporter permease |
| | P_0710 | Low specificity hydrolase |
| | P_0875 | CDP-diacylglycerol–glycerol-3-phosphate 3-phosphatidyltransferase |
| | P_0530 | Uncharacterized protein |
| | P_0652 | Preprotein translocase subunit |
| | P_0001 | Chromosomal replication initiator protein |
| | P_0838 | 23S rRNA (guanosine(2251)-2'-O)-methyltransferase |
| | P_0137 | 50S ribosomal protein L31 |
| | P_0727 | Triose-phosphate isomerase |

*continued on next page*

Table S2 – *continued from previous page*

| Module | Protein ID | Function |
|---|---|---|
| | P_0515 | Cytidine deaminase |
| | P_0431 | Uncharacterized metallophosphoesterase |
| | P_0832 | Aminoacyl-tRNA hydrolase |
| | P_0026 | Single-stranded DNA-binding protein |
| Module 34 | P_0793 | F0F1 ATP synthase subunit delta |
| | P_0346 | Uncharacterized protein |
| | P_0440 | Uncharacterized lipoprotein |
| | P_0054 | Uncharacterized peroxiredoxin |
| | P_0144 | tRNA (adenosine(37)-N6)-threonylcarbamoyltransferase complex transferase subunit |
| | P_0378 | NAD(+) synthase |
| | P_0851 | Uncharacterized lipoprotein |
| | P_0448 | Uncharacterized rRNA methyltransferase |
| | P_0047 | DNA polymerase III subunit gamma and tau |
| | P_0294 | 30S ribosomal protein S15 |
| | P_0879 | magnesium-importing ATPase |
| | P_0726 | Glucosamine-6-phosphate deaminase |
| | P_0494 | N-acetylmannosamine-6-phosphate 2-epimerase |
| | P_0530 | Uncharacterized protein |
| | P_0453 | DNA topoisomerase IV subunit A |
| | P_0302 | Flavin reductase |
| | P_0045 | dTMP kinase |
| | P_0451 | Glyceraldehyde-3-phosphate dehydrogenase |
| | P_0791 | F0F1 ATP synthase subunit gamma |

*continued on next page*

Table S2 – *continued from previous page*

| **Module** | **Protein ID** | **Function** |
|---|---|---|
| | P_0728 | Low specificity hydrolase |
| | P_0169 | Oligopeptide ABC transporter substrate-binding protein |
| | P_0697 | Uncharacterized glycosyl transferase |
| | P_0197 | Spermidine/putrescine ABC transporter ATP-binding protein |
| | P_0372 | Flippase B |
| | P_0425 | Phosphate ABC transporter substrate-binding protein |
| | P_0042 | Uncharacterized transcriptional regulator |
| | P_0695 | ATP-dependent DNA helicase |
| | P_0874 | 16S rRNA (guanine(527)-N(7))-methyltransferase |
| | P_0203 | Guanylate kinase |
| | P_0361 | 23S rRNA (pseudouridine(1915)-N3)-methyltransferase |
| | P_0303 | PolC-type DNA polymerase III |
| | P_0859 | DNA topoisomerase I |
| | P_0732 | Deoxyribose-phosphate aldolase |
| | P_0511 | Uncharacterized protein |
| | P_0482 | 30S ribosomal protein S21 |
| | P_0885 | tRNA uridine(34) 5-carboxymethylaminomethyl synthesis enzyme |
| | P_0512 | Acyl-phosphate glycerol 3-phosphate acyltransferase |
| | P_0614 | Nicotinate phosphoribosyltransferase |

*continued on next page*

Table S2 – *continued from previous page*

| **Module** | **Protein ID** | **Function** |
|---|---|---|
| | P_0615 | Uncharacterized protein |
| | P_0910 | 50S ribosomal protein L34 |
| Module 35 | P_0690 | DNA ligase (NAD(+)) |
| | P_0501 | 50S ribosomal protein L21 |
| | P_0414 | Guanosine-3',5'-bis(diphosphate) 3'-pyrophosphohydrolase |
| | P_0387 | tRNA 2-thiouridine(34) synthase |
| | P_0094 | Uncharacterized protein |
| | P_0412 | Uncharacterized protein |
| | P_0445 | Glucose-6-phosphate isomerase |
| | P_0644 | 50S ribosomal protein L17 |
| | P_0389 | Uncharacterized protein |
| | P_0346 | Uncharacterized protein |
| | P_0029 | FMN reductase |
| | P_0167 | Oligopeptide ABC transporter ATP-binding protein |
| | P_0353 | Cell division DivIVA/GpsB protein |
| | P_0348 | Ribosome biogenesis GTPase |
| | P_0641 | ECF transporter T component |
| | P_0439 | Uncharacterized lipoprotein |
| | P_0047 | DNA polymerase III subunit gamma and tau |
| | P_0166 | Oligopeptide ABC transporter permease |
| | P_0143 | Uncharacterized protein |
| | P_0639 | Uncharacterized efflux ABC transporter permease |
| | P_0840 | Antitermination protein |

*continued on next page*

Table S2 – *continued from previous page*

| Module | Protein ID | Function |
|---|---|---|
| | P_0299 | Uncharacterized protein |
| | P_0063 | Uncharacterized tRNA dihydrouridine synthase |
| | P_0431 | Uncharacterized metallophosphoesterase |
| | P_0042 | Uncharacterized transcriptional regulator |
| | P_0813 | UDP-glucose 4-epimerase GalE |
| | P_0108 | Uncharacterized lipoprotein |
| | P_0271 | tRNA N6-adenosine(37)-N6-threonylcarbamoyltransferase complex dimerization subunit |
| | P_0611 | DNA polymerase I |
| | P_0886 | Proton-glutamate symporter |
| | P_0440 | Uncharacterized lipoprotein |
| | P_0429 | Signal recognition particle-docking protein |
| | P_0046 | Recombination protein |
| | P_0879 | magnesium-importing ATPase |
| | P_0438 | Uncharacterized protein |
| | P_0002 | DNA polymerase III subunit beta |
| | P_0040 | tRNA lysidine(34) synthetase |
| | P_0165 | Oligopeptide ABC transporter permease |
| | P_0451 | Glyceraldehyde-3-phosphate dehydrogenase |
| | P_0254 | Excinuclease ABC subunit C |
| | P_0203 | Guanylate kinase |
| | P_0415 | Chromosome segregation protein |
| | P_0794 | F0F1 ATP synthase subunit B |
| | P_0229 | Phosphate acetyltransferase |

*continued on next page*

Table S2 – *continued from previous page*

| **Module** | **Protein ID** | **Function** |
|---|---|---|
| | P_0512 | Acyl-phosphate glycerol 3-phosphate acyltransferase |
| | P_0116 | Uncharacterized protein |
| | P_0240 | tRNA 4-thiouridine(8) synthase |
| | P_0859 | DNA topoisomerase I |
| | P_0800 | Ribose 5-phosphate isomerase B |
| | P_0908 | Membrane protein insertase |
| | P_0250 | Uncharacterized protein |
| | P_0145 | Uncharacterized acetyltransferase |
| | P_0005 | Uncharacterized protein |
| | P_0832 | Aminoacyl-tRNA hydrolase |
| | P_0390 | Methionyl-tRNA formyltransferase |
| | P_0419 | Phosphate acyltransferase |
| | P_0600 | Ribonuclease J |
| | P_0791 | F0F1 ATP synthase subunit gamma |
| | P_0316 | Transketolase |
| | P_0592 | Uncharacterized protein |
| Module 36 | P_0365 | 50S ribosomal protein L19 |
| | P_0646 | 30S ribosomal protein S11 |
| | P_0819 | Thioredoxin-disulfide reductase |
| | P_0608 | Primosomal protein |
| | P_0166 | Oligopeptide ABC transporter permease |
| | P_0773 | Ribonucleoside-diphosphate reductase subunit beta |
| | P_0027 | 30S ribosomal protein S6 |
| | P_0517 | Uncharacterized RNA pseudouridine synthase |

*continued on next page*

Table S2 – *continued from previous page*

| Module | Protein ID | Function |
|---|---|---|
| | P_0617 | Fatty acid binding protein |
| | P_0115 | UTP–glucose-1-phosphate uridylyltransferase |
| | P_0684 | Bifunctional 5,10-methylene-tetrahydrofolate dehydrogenase/5,10-methylene-tetrahydrofolate cyclohydrolase |
| | P_0338 | Uncharacterized lipoprotein |
| | P_0859 | DNA topoisomerase I |
| | P_0165 | Oligopeptide ABC transporter permease |
| | P_0796 | F0F1 ATP synthase subunit A |
| | P_0410 | Degradosome RNA helicase |
| | P_0452 | DNA topoisomerase IV subunit B |
| | P_0827 | Uncharacterized protein |
| | P_0605 | Uncharacterized protein |
| | P_0042 | Uncharacterized transcriptional regulator |
| | P_0800 | Ribose 5-phosphate isomerase B |
| | P_0775 | Ribonuclease R |
| | P_0029 | FMN reductase |
| | P_0772 | Ribonucleotide reductase assembly protein |
| | P_0879 | magnesium-importing ATPase |
| | P_0710 | Low specificity hydrolase |
| | P_0145 | Uncharacterized acetyltransferase |
| | P_0441 | Cysteine desulfurase |
| | P_0789 | F0F1 ATP synthase subunit delta/epsilon |
| | P_0392 | Uncharacterized protein |
| | P_0537 | UMP kinase |
| | P_0077 | Low specificity hydrolase |

*continued on next page*

Table S2 – *continued from previous page*

| **Module** | **Protein ID** | **Function** |
|---|---|---|
| | P_0139 | NanoRNAse |
| | P_0872 | Uncharacterized ATPase |
| | P_0838 | 23S rRNA (guanosine(2251)-2'-O)-methyltransferase |
| | P_0289 | Ribosome-binding factor A |
| | P_0730 | Uncharacterized protein |
| | P_0908 | Membrane protein insertase |
| | P_0116 | Uncharacterized protein |
| | P_0044 | DNA polymerase III subunit delta |
| | P_0422 | 50S ribosomal protein L28 |
| | P_0771 | Ribonucleotide-diphosphate reductase subunit alpha |
| | P_0391 | Elongation factor P |
| | P_0526 | 50S ribosomal protein L32 |
| | P_0332 | Uncharacterized protein |
| | P_0010 | Nucleoside ABC transporter ATP-binding protein |
| | P_0361 | 23S rRNA (pseudouridine(1915)-N3)-methyltransferase |
| | P_0832 | Aminoacyl-tRNA hydrolase |
| | P_0444 | Oligopeptidase |
| | P_0366 | L16-binding dependent 50S subunit-maturation GTPase |
| | P_0412 | Uncharacterized protein |
| | P_0799 | Serine hydroxymethyltransferase |
| | P_0006 | DNA gyrase subunit B |

*continued on next page*

Table S2 – *continued from previous page*

| Module | Protein ID | Function |
|---|---|---|
| Module 37 | P_0389 | Uncharacterized protein |
| | P_0271 | tRNA N6-adenosine(37)-N6-threonylcarbamoyltransferase complex dimerization subunit |
| | P_0592 | Uncharacterized protein |
| | P_0375 | Uncharacterized protein |
| | P_0650 | Type I methionyl aminopeptidase |
| | P_0886 | Proton-glutamate symporter |
| | P_0444 | Oligopeptidase |
| | P_0422 | 50S ribosomal protein L28 |
| | P_0708 | Thiamine ABC transporter substrate-binding protein |
| | P_0409 | Uncharacterized protein |
| | P_0791 | F0F1 ATP synthase subunit gamma |
| | P_0006 | DNA gyrase subunit B |
| | P_0329 | Uncharacterized pseudouridine synthase |
| | P_0257 | RNase J family beta-CASP ribonuclease |
| | P_0202 | 16S rRNA (guanine(966)-N(2))-methyltransferase |
| | P_0834 | Replicative DNA helicase |
| | P_0391 | Elongation factor P |
| | P_0249 | Uncharacterized protein |
| | P_0661 | 50S ribosomal protein L14 |
| | P_0005 | Uncharacterized protein |
| | P_0133 | Toxin-antitoxin serine protease |
| | P_0165 | Oligopeptide ABC transporter permease |

*continued on next page*

Table S2 – *continued from previous page*

| **Module** | **Protein ID** | **Function** |
|---|---|---|
| | P_0543 | Nucleotide exchange factor |
| | P_0537 | UMP kinase |
| | P_0875 | CDP-diacylglycerol–glycerol-3-phosphate 3-phosphatidyltransferase |
| | P_0116 | Uncharacterized protein |
| | P_0645 | DNA-directed RNA polymerase subunit alpha |
| | P_0330 | Deoxyguanosine kinase |
| | P_0262 | Ribulose-phosphate 3-epimerase |
| | P_0651 | Adenylate kinase |
| | P_0009 | Nucleoside ABC transporter permease |
| | P_0047 | DNA polymerase III subunit gamma and tau |
| | P_0435 | Mannose-6-phosphate isomerase |
| | P_0132 | Toxin-antitoxin AAA ATPase |
| | P_0526 | 50S ribosomal protein L32 |
| | P_0377 | Ribosome GTPase |
| | P_0772 | Ribonucleotide reductase assembly protein |
| | P_0303 | PolC-type DNA polymerase III |
| | P_0726 | Glucosamine-6-phosphate deaminase |
| | P_0390 | Methionyl-tRNA formyltransferase |
| | P_0228 | Dihydrolipoyl dehydrogenase |
| | P_0523 | Cell division protein |
| | P_0338 | Uncharacterized lipoprotein |
| | P_0600 | Ribonuclease J |
| | P_0548 | tRNA (cytidine(34)-2'-O)-methyltransferase |
| | P_0010 | Nucleoside ABC transporter ATP-binding protein |

*continued on next page*

Table S2 – *continued from previous page*

| Module | Protein ID | Function |
|---|---|---|
| | P_0505 | Uncharacterized lipoprotein |
| | P_0610 | DNA-formamidopyrimidine glycosylase |
| | P_0314 | Uncharacterized ECF transporter S component |
| | P_0065 | Thioredoxin |
| | P_0732 | Deoxyribose-phosphate aldolase |
| | P_0452 | DNA topoisomerase IV subunit B |
| | P_0042 | Uncharacterized transcriptional regulator |
| | P_0872 | Uncharacterized ATPase |
| | P_0039 | Membrane anchored AAA+ protease |
| Module 38 | P_0876 | Uncharacterized amino acid permease |
| | P_0428 | Phosphate transport system regulatory protein |
| | P_0791 | F0F1 ATP synthase subunit gamma |
| | P_0316 | Transketolase |
| | P_0420 | Fatty acid kinase subunit A |
| | P_0511 | Uncharacterized protein |
| | P_0432 | Methionine adenosyltransferase |
| | P_0139 | NanoRNAse |
| | P_0651 | Adenylate kinase |
| | P_0707 | Thiamine ABC transporter ATP-binding protein |
| | P_0429 | Signal recognition particle-docking protein |
| | P_0377 | Ribosome GTPase |
| | P_0009 | Nucleoside ABC transporter permease |
| | P_0695 | ATP-dependent DNA helicase |
| | P_0686 | Potassium transporter |

*continued on next page*

Table S2 – *continued from previous page*

| Module | Protein ID | Function |
|---|---|---|
| | P_0010 | Nucleoside ABC transporter ATP-binding protein |
| | P_0541 | Molecular chaperone |
| | P_0874 | 16S rRNA (guanine(527)-N(7))-methyltransferase |
| | P_0821 | HPr(Ser) kinase/phosphatase |
| | P_0039 | Membrane anchored AAA+ protease |
| | P_0827 | Uncharacterized protein |
| | P_0517 | Uncharacterized RNA pseudouridine synthase |
| | P_0108 | Uncharacterized lipoprotein |
| | P_0879 | magnesium-importing ATPase |
| | P_0203 | Guanylate kinase |
| | P_0732 | Deoxyribose-phosphate aldolase |
| | P_0859 | DNA topoisomerase I |
| | P_0832 | Aminoacyl-tRNA hydrolase |
| | P_0592 | Uncharacterized protein |
| | P_0195 | Spermidine/putrescine ABC transporter permease |
| | P_0302 | Flavin reductase |
| | P_0126 | Glutamate–tRNA ligase |
| | P_0065 | Thioredoxin |
| | P_0885 | tRNA uridine(34) 5-carboxymethylaminomethyl synthesis enzyme |
| | P_0689 | Glutamyl-tRNA amidotransferase subunit C |
| | P_0117 | Acyl-phosphate glycerol 3-phosphate acyltransferase |

*continued on next page*

Table S2 – *continued from previous page*

| Module | Protein ID | Function |
|---|---|---|
| | P_0001 | Chromosomal replication initiator protein |
| | P_0419 | Phosphate acyltransferase |
| | P_0138 | Uncharacterized protein |
| Module 39 | P_0044 | DNA polymerase III subunit delta |
| | P_0332 | Uncharacterized protein |
| | P_0605 | Uncharacterized protein |
| | P_0604 | Uncharacterized protein |
| | P_0851 | Uncharacterized lipoprotein |
| | P_0285 | Elongation factor 4 |
| | P_0002 | DNA polymerase III subunit beta |
| | P_0045 | dTMP kinase |
| | P_0304 | Phosphatidate cytidylyltransferase |
| | P_0520 | Uncharacterized hydrolase |
| | P_0728 | Low specificity hydrolase |
| | P_0776 | SsrA-binding protein |
| | P_0063 | Uncharacterized tRNA dihydrouridine synthase |
| | P_0421 | Uncharacterized protein |
| | P_0229 | Phosphate acetyltransferase |
| | P_0361 | 23S rRNA (pseudouridine(1915)-N3)-methyltransferase |
| | P_0114 | Glycolipid synthase B |
| | P_0451 | Glyceraldehyde-3-phosphate dehydrogenase |
| | P_0824 | Excinuclease ABC subunit A |
| | P_0377 | Ribosome GTPase |
| | P_0517 | Uncharacterized RNA pseudouridine synthase |
| | P_0779 | PTS sugar transporter |

*continued on next page*

Table S2 – *continued from previous page*

| Module | Protein ID | Function |
|---|---|---|
| | P_0422 | 50S ribosomal protein L28 |
| | P_0427 | Phosphate ABC transporter ATP-binding protein |
| | P_0240 | tRNA 4-thiouridine(8) synthase |
| | P_0648 | 50S ribosomal protein L36 |
| | P_0526 | 50S ribosomal protein L32 |
| | P_0789 | F0F1 ATP synthase subunit delta/epsilon |
| | P_0380 | Nicotinate (nicotinamide) nucleotide adenylyltransferase |
| | P_0791 | F0F1 ATP synthase subunit gamma |
| | P_0689 | Glutamyl-tRNA amidotransferase subunit C |
| | P_0081 | tRNA uridine(34) 5-carboxymethylaminomethyl synthesis GTPase |
| | P_0249 | Uncharacterized protein |
| | P_0145 | Uncharacterized acetyltransferase |
| | P_0908 | Membrane protein insertase |
| | P_0410 | Degradosome RNA helicase |
| | P_0390 | Methionyl-tRNA formyltransferase |
| | P_0773 | Ribonucleoside-diphosphate reductase subunit beta |
| | P_0444 | Oligopeptidase |
| | P_0117 | Acyl-phosphate glycerol 3-phosphate acyltransferase |
| | P_0039 | Membrane anchored AAA+ protease |
| | P_0371 | Flippase A |
| | P_0516 | Uncharacterized protein |

*continued on next page*

Table S2 – *continued from previous page*

| **Module** | **Protein ID** | **Function** |
|---|---|---|
| | P_0139 | NanoRNAse |
| | P_0043 | Uncharacterized methyltransferase |
| | P_0813 | UDP-glucose 4-epimerase GalE |
| | P_0906 | Uncharacterized protein |
| | P_0732 | Deoxyribose-phosphate aldolase |
| | P_0115 | UTP–glucose-1-phosphate uridylyltransferase |
| | P_0116 | Uncharacterized protein |
| | P_0859 | DNA topoisomerase I |
| | P_0065 | Thioredoxin |
| | P_0775 | Ribonuclease R |
| | P_0726 | Glucosamine-6-phosphate deaminase |
| | P_0314 | Uncharacterized ECF transporter S component |
| | P_0688 | Glutamyl-tRNA amidotransferase subunit A |
| | P_0636 | Uncharacterized lipoprotein |
| | P_0601 | Uncharacterized protein |
| | P_0530 | Uncharacterized protein |
| | P_0429 | Signal recognition particle-docking protein |
| | P_0294 | 30S ribosomal protein S15 |
| | P_0291 | FAD synthetase |
| | P_0006 | DNA gyrase subunit B |
| Module 40 | P_0006 | DNA gyrase subunit B |
| | P_0831 | Phosphoribosylpyrophosphate synthetase |
| | P_0639 | Uncharacterized efflux ABC transporter permease |
| | P_0428 | Phosphate transport system regulatory protein |
| | P_0254 | Excinuclease ABC subunit C |

*continued on next page*

Table S2 – *continued from previous page*

| Module | Protein ID | Function |
|---|---|---|
| | P_0353 | Cell division DivIVA/GpsB protein |
| | P_0005 | Uncharacterized protein |
| | P_0441 | Cysteine desulfurase |
| | P_0378 | NAD(+) synthase |
| | P_0419 | Phosphate acyltransferase |
| | P_0144 | tRNA (adenosine(37)-N6)-threonylcarbamoyltransferase complex transferase subunit |
| | P_0611 | DNA polymerase I |
| | P_0412 | Uncharacterized protein |
| | P_0772 | Ribonucleotide reductase assembly protein |
| | P_0329 | Uncharacterized pseudouridine synthase |
| | P_0439 | Uncharacterized lipoprotein |
| | P_0372 | Flippase B |
| | P_0819 | Thioredoxin-disulfide reductase |
| | P_0779 | PTS sugar transporter |
| | P_0301 | Ribosome assembly cofactor |
| | P_0642 | ECF transporter ATPase |
| | P_0214 | Phosphatidylglycerophosphatase |
| | P_0411 | Uncharacterized protein |
| | P_0004 | 16S rRNA (adenine(1518)-N(6)/adenine(1519)-N(6))- dimethyltransferase |
| | P_0794 | F0F1 ATP synthase subunit B |
| | P_0203 | Guanylate kinase |
| | P_0885 | tRNA uridine(34) 5-carboxymethylaminomethyl synthesis enzyme |

*continued on next page*

Table S2 – *continued from previous page*

| Module | Protein ID | Function |
|---|---|---|
| | P_0452 | DNA topoisomerase IV subunit B |
| | P_0511 | Uncharacterized protein |
| | P_0652 | Preprotein translocase subunit |
| | P_0228 | Dihydrolipoyl dehydrogenase |
| | P_0249 | Uncharacterized protein |
| | P_0840 | Antitermination protein |
| | P_0202 | 16S rRNA (guanine(966)-N(2))-methyltransferase |
| | P_0537 | UMP kinase |
| | P_0710 | Low specificity hydrolase |
| | P_0610 | DNA-formamidopyrimidine glycosylase |
| | P_0648 | 50S ribosomal protein L36 |
| | P_0330 | Deoxyguanosine kinase |
| | P_0799 | Serine hydroxymethyltransferase |
| | P_0608 | Primosomal protein |
| | P_0824 | Excinuclease ABC subunit A |
| | P_0517 | Uncharacterized RNA pseudouridine synthase |
| | P_0145 | Uncharacterized acetyltransferase |
| | P_0201 | Peptide deformylase |
| | P_0482 | 30S ribosomal protein S21 |
| | P_0872 | Uncharacterized ATPase |
| | P_0886 | Proton-glutamate symporter |
| | P_0250 | Uncharacterized protein |
| | P_0422 | 50S ribosomal protein L28 |
| | P_0800 | Ribose 5-phosphate isomerase B |
| | P_0887 | Coenzyme A Disulfide Reductase |

*continued on next page*

Table S2 – *continued from previous page*

| Module | Protein ID | Function |
|---|---|---|
| | P_0932 | 50S ribosomal protein L33 |
| | P_0838 | 23S rRNA (guanosine(2251)-2'-O)-methyltransferase |
| | P_0859 | DNA topoisomerase I |
| | P_0645 | DNA-directed RNA polymerase subunit alpha |
| | P_0697 | Uncharacterized glycosyl transferase |
| | P_0294 | 30S ribosomal protein S15 |
| | P_0881 | Uncharacterized MFS transporter |
| | P_0728 | Low specificity hydrolase |
| | P_0512 | Acyl-phosphate glycerol 3-phosphate acyltransferase |
| | P_0695 | ATP-dependent DNA helicase |
| | P_0302 | Flavin reductase |
| | P_0516 | Uncharacterized protein |
| | P_0832 | Aminoacyl-tRNA hydrolase |
| | P_0371 | Flippase A |
| | P_0733 | Phosphopentomutase |
| | P_0001 | Chromosomal replication initiator protein |
| | P_0361 | 23S rRNA (pseudouridine(1915)-N3)-methyltransferase |
| | P_0392 | Uncharacterized protein |
| Module 41 | P_0407 | RNA polymerase subunit sigma |
| | P_0505 | Uncharacterized lipoprotein |
| | P_0166 | Oligopeptide ABC transporter permease |
| | P_0840 | Antitermination protein |
| | P_0060 | Uncharacterized protein |

*continued on next page*

Table S2 – *continued from previous page*

| Module | Protein ID | Function |
|---|---|---|
| | P_0430 | Uncharacterized DNA-binding protein |
| | P_0414 | Guanosine-3',5'-bis(diphosphate) 3'-pyrophosphohydrolase |
| | P_0538 | Uncharacterized protein |
| | P_0823 | Dihydrofolate synthase |
| | P_0148 | 30S ribosomal protein S12 |
| | P_0918 | Imidazoleglycerol-phosphate dehydratase |
| | P_0420 | Fatty acid kinase subunit A |
| | P_0301 | Ribosome assembly cofactor |
| | P_0029 | FMN reductase |
| | P_0639 | Uncharacterized efflux ABC transporter permease |
| | P_0126 | Glutamate–tRNA ligase |
| | P_0835 | Uncharacterized lipoprotein |
| | P_0429 | Signal recognition particle-docking protein |
| | P_0932 | 50S ribosomal protein L33 |
| | P_0344 | Inorganic diphosphatase |
| | P_0604 | Uncharacterized protein |
| | P_0138 | Uncharacterized protein |
| | P_0065 | Thioredoxin |
| | P_0813 | UDP-glucose 4-epimerase GalE |
| | P_0636 | Uncharacterized lipoprotein |
| | P_0733 | Phosphopentomutase |
| | P_0141 | Peptide chain release factor 1 |
| | P_0410 | Degradosome RNA helicase |
| | P_0228 | Dihydrolipoyl dehydrogenase |

*continued on next page*

Table S2 – *continued from previous page*

| Module | Protein ID | Function |
|---|---|---|
| | P_0834 | Replicative DNA helicase |
| | P_0623 | Uncharacterized protein |
| | P_0006 | DNA gyrase subunit B |
| | P_0378 | NAD(+) synthase |
| | P_0197 | Spermidine/putrescine ABC transporter ATP-binding protein |
| | P_0262 | Ribulose-phosphate 3-epimerase |
| | P_0499 | 50S ribosomal protein L27 |
| | P_0651 | Adenylate kinase |
| | P_0877 | Riboflavin ECF transporter S component |
| | P_0526 | 50S ribosomal protein L32 |
| | P_0641 | ECF transporter T component |
| | P_0776 | SsrA-binding protein |
| | P_0691 | Uncharacterized protein |
| | P_0045 | dTMP kinase |
| | P_0167 | Oligopeptide ABC transporter ATP-binding protein |
| | P_0664 | 50S ribosomal protein L16 |
| | P_0838 | 23S rRNA (guanosine(2251)-2'-O)-methyltransferase |
| | P_0799 | Serine hydroxymethyltransferase |
| | P_0438 | Uncharacterized protein |
| | P_0689 | Glutamyl-tRNA amidotransferase subunit C |
| | P_0291 | FAD synthetase |
| | P_0549 | Non-canonical purine NTP pyrophosphatase |
| | P_0791 | F0F1 ATP synthase subunit gamma |

*continued on next page*

Table S2 – *continued from previous page*

| Module | Protein ID | Function |
|---|---|---|
| | P_0147 | Cardiolipin synthase |
| | P_0271 | tRNA N6-adenosine(37)-N6-threonylcarbamoyltransferase complex dimerization subunit |
| | P_0195 | Spermidine/putrescine ABC transporter permease |
| | P_0726 | Glucosamine-6-phosphate deaminase |
| | P_0079 | tRNA (adenosine(37)-N6)-threonylcarbamoyltransferase complex transferase subunit D |
| | P_0077 | Low specificity hydrolase |
| | P_0044 | DNA polymerase III subunit delta |
| | P_0359 | Ribonuclease Y |
| | P_0418 | Ribonuclease III |
| | P_0203 | Guanylate kinase |
| | P_0415 | Chromosome segregation protein |
| | P_0240 | tRNA 4-thiouridine(8) synthase |
| Module 42 | P_0126 | Glutamate–tRNA ligase |
| | P_0689 | Glutamyl-tRNA amidotransferase subunit C |
| | P_0819 | Thioredoxin-disulfide reductase |
| | P_0026 | Single-stranded DNA-binding protein |
| | P_0140 | Thymidine kinase |
| | P_0775 | Ribonuclease R |
| | P_0410 | Degradosome RNA helicase |
| | P_0044 | DNA polymerase III subunit delta |

*continued on next page*

Table S2 – *continued from previous page*

| Module | Protein ID | Function |
|---|---|---|
| | P_0197 | Spermidine/putrescine ABC transporter ATP-binding protein |
| | P_0294 | 30S ribosomal protein S15 |
| | P_0541 | Molecular chaperone |
| | P_0129 | CTP synthase |
| | P_0289 | Ribosome-binding factor A |
| | P_0145 | Uncharacterized acetyltransferase |
| | P_0065 | Thioredoxin |
| | P_0592 | Uncharacterized protein |
| | P_0148 | 30S ribosomal protein S12 |
| | P_0402 | rRNA maturation RNase |
| | P_0615 | Uncharacterized protein |
| | P_0407 | RNA polymerase subunit sigma |
| | P_0544 | Heat-inducible transcription repressor |
| | P_0870 | Uncharacterized C4-dicarboxylate ABC transporter |
| | P_0428 | Phosphate transport system regulatory protein |
| | P_0228 | Dihydrolipoyl dehydrogenase |
| | P_0364 | tRNA (guanosine(37)-N1)-methyltransferase |
| | P_0708 | Thiamine ABC transporter substrate-binding protein |
| | P_0113 | Glycolipid synthase A |
| | P_0695 | ATP-dependent DNA helicase |
| | P_0792 | F0F1 ATP synthase subunit alpha |
| | P_0614 | Nicotinate phosphoribosyltransferase |
| | P_0301 | Ribosome assembly cofactor |

*continued on next page*

Table S2 – *continued from previous page*

| Module | Protein ID | Function |
|---|---|---|
| | P_0610 | DNA-formamidopyrimidine glycosylase |
| | P_0195 | Spermidine/putrescine ABC transporter permease |
| | P_0047 | DNA polymerase III subunit gamma and tau |
| | P_0611 | DNA polymerase I |
| | P_0249 | Uncharacterized protein |
| | P_0165 | Oligopeptide ABC transporter permease |
| | P_0908 | Membrane protein insertase |
| | P_0010 | Nucleoside ABC transporter ATP-binding protein |
| | P_0139 | NanoRNAse |
| | P_0199 | 50S ribosomal protein L35 |
| | P_0684 | Bifunctional 5,10-methylene-tetrahydrofolate dehydrogenase/5,10-methylene-tetrahydrofolate cyclohydrolase |
| | P_0429 | Signal recognition particle-docking protein |
| | P_0515 | Cytidine deaminase |
| Module 43 | P_0831 | Phosphoribosylpyrophosphate synthetase |
| | P_0439 | Uncharacterized lipoprotein |
| | P_0304 | Phosphatidate cytidylyltransferase |
| | P_0482 | 30S ribosomal protein S21 |
| | P_0328 | Chromosome segregation protein B |
| | P_0827 | Uncharacterized protein |
| | P_0360 | Signal recognition particle protein |
| | P_0544 | Heat-inducible transcription repressor |

*continued on next page*

Table S2 – *continued from previous page*

| Module | Protein ID | Function |
|---|---|---|
| | P_0875 | CDP-diacylglycerol–glycerol-3-phosphate 3-phosphatidyltransferase |
| | P_0881 | Uncharacterized MFS transporter |
| | P_0434 | 23S rRNA (uridine(1939)-m5)-methyltransferase |
| | P_0364 | tRNA (guanosine(37)-N1)-methyltransferase |
| | P_0347 | Cytidylate kinase |
| | P_0346 | Uncharacterized protein |
| | P_0695 | ATP-dependent DNA helicase |
| | P_0138 | Uncharacterized protein |
| | P_0415 | Chromosome segregation protein |
| | P_0645 | DNA-directed RNA polymerase subunit alpha |
| | P_0790 | F0F1 ATP synthase subunit beta |
| | P_0338 | Uncharacterized lipoprotein |
| | P_0294 | 30S ribosomal protein S15 |
| | P_0610 | DNA-formamidopyrimidine glycosylase |
| | P_0407 | RNA polymerase subunit sigma |
| | P_0029 | FMN reductase |
| | P_0420 | Fatty acid kinase subunit A |
| | P_0834 | Replicative DNA helicase |
| | P_0608 | Primosomal protein |
| | P_0432 | Methionine adenosyltransferase |
| | P_0918 | Imidazoleglycerol-phosphate dehydratase |
| | P_0249 | Uncharacterized protein |
| | P_0650 | Type I methionyl aminopeptidase |
| | P_0353 | Cell division DivIVA/GpsB protein |

*continued on next page*

Table S2 – *continued from previous page*

| Module | Protein ID | Function |
|---|---|---|
| | P_0197 | Spermidine/putrescine ABC transporter ATP-binding protein |
| | P_0303 | PolC-type DNA polymerase III |
| | P_0247 | Ribosome biogenesis GTP-binding protein |
| | P_0649 | Translation initiation factor IF-1 |
| | P_0708 | Thiamine ABC transporter substrate-binding protein |
| | P_0819 | Thioredoxin-disulfide reductase |
| | P_0203 | Guanylate kinase |
| | P_0779 | PTS sugar transporter |
| | P_0906 | Uncharacterized protein |
| | P_0412 | Uncharacterized protein |
| | P_0428 | Phosphate transport system regulatory protein |
| | P_0005 | Uncharacterized protein |
| | P_0402 | rRNA maturation RNase |
| | P_0512 | Acyl-phosphate glycerol 3-phosphate acyltransferase |
| | P_0010 | Nucleoside ABC transporter ATP-binding protein |
| | P_0301 | Ribosome assembly cofactor |
| | P_0669 | 50S ribosomal protein L23 |
| | P_0452 | DNA topoisomerase IV subunit B |
| | P_0047 | DNA polymerase III subunit gamma and tau |
| | P_0689 | Glutamyl-tRNA amidotransferase subunit C |
| | P_0148 | 30S ribosomal protein S12 |
| | P_0316 | Transketolase |

*continued on next page*

Table S2 – *continued from previous page*

| Module | Protein ID | Function |
|---|---|---|
| | P_0910 | 50S ribosomal protein L34 |
| | P_0373 | Uncharacterized protein |
| | P_0445 | Glucose-6-phosphate isomerase |
| | P_0636 | Uncharacterized lipoprotein |
| | P_0299 | Uncharacterized protein |
| | P_0371 | Flippase A |
| | P_0887 | Coenzyme A Disulfide Reductase |
| | P_0430 | Uncharacterized DNA-binding protein |
| | P_0002 | DNA polymerase III subunit beta |
| | P_0614 | Nicotinate phosphoribosyltransferase |
| | P_0011 | Nucleoside ABC transporter substrate-binding protein |
| | P_0238 | 30S ribosomal protein S4 |
| | P_0289 | Ribosome-binding factor A |
| | P_0063 | Uncharacterized tRNA dihydrouridine synthase |
| | P_0081 | tRNA uridine(34) 5-carboxymethylaminomethyl synthesis GTPase |
| | P_0201 | Peptide deformylase |
| | P_0611 | DNA polymerase I |
| | P_0389 | Uncharacterized protein |
| | P_0140 | Thymidine kinase |
| | P_0054 | Uncharacterized peroxiredoxin |
| | P_0688 | Glutamyl-tRNA amidotransferase subunit A |
| | P_0239 | Uncharacterized protein |
| | P_0617 | Fatty acid binding protein |
| | P_0824 | Excinuclease ABC subunit A |

*continued on next page*

Table S2 – *continued from previous page*

| **Module** | **Protein ID** | **Function** |
|---|---|---|
| | P_0832 | Aminoacyl-tRNA hydrolase |
| | P_0733 | Phosphopentomutase |
| | P_0548 | tRNA (cytidine(34)-2'-O)-methyltransferase |
| | P_0686 | Potassium transporter |
| | P_0065 | Thioredoxin |
| | P_0821 | HPr(Ser) kinase/phosphatase |
| Module 44 | P_0639 | Uncharacterized efflux ABC transporter permease |
| | P_0113 | Glycolipid synthase A |
| | P_0661 | 50S ribosomal protein L14 |
| | P_0271 | tRNA N6-adenosine(37)-N6-threonylcarbamoyltransferase complex dimerization subunit |
| | P_0835 | Uncharacterized lipoprotein |
| | P_0410 | Degradosome RNA helicase |
| | P_0692 | Uncharacterized pseudouridine synthase |
| | P_0366 | L16-binding dependent 50S subunit-maturation GTPase |
| | P_0434 | 23S rRNA (uridine(1939)-m5)-methyltransferase |
| | P_0126 | Glutamate–tRNA ligase |
| | P_0427 | Phosphate ABC transporter ATP-binding protein |
| | P_0047 | DNA polymerase III subunit gamma and tau |
| | P_0347 | Cytidylate kinase |
| | P_0799 | Serine hydroxymethyltransferase |

*continued on next page*

Table S2 – *continued from previous page*

| **Module** | **Protein ID** | **Function** |
| --- | --- | --- |
| | P_0779 | PTS sugar transporter |
| | P_0516 | Uncharacterized protein |
| | P_0636 | Uncharacterized lipoprotein |
| | P_0870 | Uncharacterized C4-dicarboxylate ABC transporter |
| | P_0285 | Elongation factor 4 |
| | P_0109 | Deoxyribonuclease IV |
| | P_0365 | 50S ribosomal protein L19 |
| | P_0362 | 30S ribosomal protein S16 |
| | P_0695 | ATP-dependent DNA helicase |
| | P_0377 | Ribosome GTPase |
| | P_0065 | Thioredoxin |
| | P_0409 | Uncharacterized protein |
| | P_0145 | Uncharacterized acetyltransferase |
| | P_0827 | Uncharacterized protein |
| | P_0294 | 30S ribosomal protein S15 |
| | P_0515 | Cytidine deaminase |
| | P_0605 | Uncharacterized protein |
| | P_0234 | PTS glucose transporter subunit IIA |
| | P_0314 | Uncharacterized ECF transporter S component |
| | P_0010 | Nucleoside ABC transporter ATP-binding protein |
| | P_0316 | Transketolase |
| | P_0549 | Non-canonical purine NTP pyrophosphatase |
| | P_0168 | Oligopeptide ABC transporter ATP-binding protein |

*continued on next page*

Table S2 – *continued from previous page*

| Module | Protein ID | Function |
|---|---|---|
| | P_0690 | DNA ligase (NAD(+)) |
| | P_0131 | Fructose-1,6-bisphosphate aldolase |
| | P_0200 | Translation initiation factor IF-3 |
| | P_0411 | Uncharacterized protein |
| | P_0228 | Dihydrolipoyl dehydrogenase |
| | P_0838 | 23S rRNA (guanosine(2251)-2'-O)-methyltransferase |
| | P_0026 | Single-stranded DNA-binding protein |
| | P_0254 | Excinuclease ABC subunit C |
| | P_0601 | Uncharacterized protein |
| | P_0202 | 16S rRNA (guanine(966)-N(2))-methyltransferase |
| | P_0650 | Type I methionyl aminopeptidase |
| | P_0651 | Adenylate kinase |
| | P_0229 | Phosphate acetyltransferase |
| | P_0592 | Uncharacterized protein |
| | P_0414 | Guanosine-3',5'-bis(diphosphate) 3'-pyrophosphohydrolase |
| | P_0453 | DNA topoisomerase IV subunit A |
| | P_0859 | DNA topoisomerase I |
| | P_0116 | Uncharacterized protein |
| | P_0042 | Uncharacterized transcriptional regulator |
| | P_0133 | Toxin-antitoxin serine protease |
| | P_0794 | F0F1 ATP synthase subunit B |
| Module 45 | P_0637 | 30S ribosomal protein S9 |
| | P_0365 | 50S ribosomal protein L19 |

*continued on next page*

Table S2 – *continued from previous page*

| **Module** | **Protein ID** | **Function** |
|---|---|---|
| | P_0881 | Uncharacterized MFS transporter |
| | P_0727 | Triose-phosphate isomerase |
| | P_0006 | DNA gyrase subunit B |
| | P_0388 | Uncharacterized protein |
| | P_0281 | Uncharacterized protein |
| | P_0839 | Preprotein translocase subunit |
| | P_0827 | Uncharacterized protein |
| | P_0664 | 50S ribosomal protein L16 |
| | P_0918 | Imidazoleglycerol-phosphate dehydratase |
| | P_0886 | Proton-glutamate symporter |
| | P_0381 | 5'-Methylthioadenosine nucleosidase |
| | P_0007 | DNA gyrase subunit A |
| | P_0686 | Potassium transporter |
| | P_0422 | 50S ribosomal protein L28 |
| | P_0114 | Glycolipid synthase B |
| | P_0373 | Uncharacterized protein |
| | P_0249 | Uncharacterized protein |
| | P_0262 | Ribulose-phosphate 3-epimerase |
| | P_0407 | RNA polymerase subunit sigma |
| | P_0005 | Uncharacterized protein |
| | P_0347 | Cytidylate kinase |
| | P_0239 | Uncharacterized protein |
| | P_0115 | UTP–glucose-1-phosphate uridylyltransferase |
| | P_0814 | UDP-galactopyranose mutase |
| | P_0360 | Signal recognition particle protein |
| | P_0097 | Flap endonuclease and 5' exonuclease |

*continued on next page*

Table S2 – *continued from previous page*

| **Module** | **Protein ID** | **Function** |
|---|---|---|
| | P_0819 | Thioredoxin-disulfide reductase |
| | P_0338 | Uncharacterized lipoprotein |
| | P_0165 | Oligopeptide ABC transporter permease |
| | P_0799 | Serine hydroxymethyltransferase |
| | P_0063 | Uncharacterized tRNA dihydrouridine synthase |
| | P_0439 | Uncharacterized lipoprotein |
| | P_0537 | UMP kinase |
| | P_0382 | Deoxynucleoside kinase |
| | P_0329 | Uncharacterized pseudouridine synthase |
| | P_0617 | Fatty acid binding protein |
| | P_0285 | Elongation factor 4 |
| | P_0004 | 16S rRNA (adenine(1518)-N(6)/adenine(1519)-N(6))- dimethyltransferase |
| | P_0444 | Oligopeptidase |
| | P_0440 | Uncharacterized lipoprotein |
| | P_0378 | NAD(+) synthase |
| | P_0728 | Low specificity hydrolase |
| | P_0330 | Deoxyguanosine kinase |
| | P_0512 | Acyl-phosphate glycerol 3-phosphate acyltransferase |
| | P_0362 | 30S ribosomal protein S16 |
| | P_0420 | Fatty acid kinase subunit A |
| | P_0623 | Uncharacterized protein |
| | P_0791 | F0F1 ATP synthase subunit gamma |
| | P_0430 | Uncharacterized DNA-binding protein |
| | P_0793 | F0F1 ATP synthase subunit delta |

*continued on next page*

Table S2 – *continued from previous page*

| **Module** | **Protein ID** | **Function** |
|---|---|---|
| | P_0823 | Dihydrofolate synthase |
| | P_0427 | Phosphate ABC transporter ATP-binding protein |
| | P_0835 | Uncharacterized lipoprotein |
| | P_0138 | Uncharacterized protein |
| | P_0299 | Uncharacterized protein |
| | P_0691 | Uncharacterized protein |
| | P_0054 | Uncharacterized peroxiredoxin |
| | P_0001 | Chromosomal replication initiator protein |
| | P_0530 | Uncharacterized protein |
| | P_0708 | Thiamine ABC transporter substrate-binding protein |
| | P_0146 | Uncharacterized protein |
| | P_0428 | Phosphate transport system regulatory protein |
| | P_0482 | 30S ribosomal protein S21 |
| | P_0684 | Bifunctional 5,10-methylene-tetrahydrofolate dehydrogenase/5,10-methylene-tetrahydrofolate cyclohydrolase |
| | P_0133 | Toxin-antitoxin serine protease |
| | P_0794 | F0F1 ATP synthase subunit B |
| | P_0410 | Degradosome RNA helicase |
| | P_0361 | 23S rRNA (pseudouridine(1915)-N3)-methyltransferase |
| | P_0438 | Uncharacterized protein |
| Module 46 | P_0303 | PolC-type DNA polymerase III |
| | P_0199 | 50S ribosomal protein L35 |

*continued on next page*

| **Module** | **Protein ID** | **Function** |
|---|---|---|
| | P_0250 | Uncharacterized protein |
| | P_0859 | DNA topoisomerase I |
| | P_0789 | F0F1 ATP synthase subunit delta/epsilon |
| | P_0610 | DNA-formamidopyrimidine glycosylase |
| | P_0548 | tRNA (cytidine(34)-2'-O)-methyltransferase |
| | P_0645 | DNA-directed RNA polymerase subunit alpha |
| | P_0026 | Single-stranded DNA-binding protein |
| | P_0097 | Flap endonuclease and 5' exonuclease |
| | P_0827 | Uncharacterized protein |
| | P_0195 | Spermidine/putrescine ABC transporter permease |
| | P_0824 | Excinuclease ABC subunit A |
| | P_0453 | DNA topoisomerase IV subunit A |
| | P_0819 | Thioredoxin-disulfide reductase |
| | P_0201 | Peptide deformylase |
| | P_0530 | Uncharacterized protein |
| | P_0427 | Phosphate ABC transporter ATP-binding protein |
| | P_0054 | Uncharacterized peroxiredoxin |
| | P_0684 | Bifunctional 5,10-methylene-tetrahydrofolate dehydrogenase/5,10-methylene-tetrahydrofolate cyclohydrolase |
| | P_0501 | 50S ribosomal protein L21 |
| | P_0775 | Ribonuclease R |
| | P_0228 | Dihydrolipoyl dehydrogenase |
| | P_0515 | Cytidine deaminase |

*continued on next page*

Table S2 – *continued from previous page*

| Module | Protein ID | Function |
|---|---|---|
| | P_0710 | Low specificity hydrolase |
| | P_0834 | Replicative DNA helicase |
| | P_0063 | Uncharacterized tRNA dihydrouridine synthase |
| | P_0127 | Uncharacterized phosphohydrolase |
| | P_0832 | Aminoacyl-tRNA hydrolase |
| | P_0428 | Phosphate transport system regulatory protein |
| | P_0452 | DNA topoisomerase IV subunit B |
| | P_0874 | 16S rRNA (guanine(527)-N(7))-methyltransferase |
| | P_0549 | Non-canonical purine NTP pyrophosphatase |
| | P_0081 | tRNA uridine(34) 5-carboxymethylaminomethyl synthesis GTPase |
| | P_0412 | Uncharacterized protein |
| | P_0371 | Flippase A |
| | P_0592 | Uncharacterized protein |
| | P_0611 | DNA polymerase I |
| | P_0444 | Oligopeptidase |
| | P_0419 | Phosphate acyltransferase |
| | P_0429 | Signal recognition particle-docking protein |
| Module 47 | P_0392 | Uncharacterized protein |
| | P_0516 | Uncharacterized protein |
| | P_0097 | Flap endonuclease and 5' exonuclease |
| | P_0047 | DNA polymerase III subunit gamma and tau |
| | P_0387 | tRNA 2-thiouridine(34) synthase |
| | P_0687 | Glutamyl-tRNA amidotransferase subunit B |
| | P_0517 | Uncharacterized RNA pseudouridine synthase |

*continued on next page*

Table S2 – *continued from previous page*

| Module | Protein ID | Function |
|---|---|---|
| | P_0727 | Triose-phosphate isomerase |
| | P_0642 | ECF transporter ATPase |
| | P_0108 | Uncharacterized lipoprotein |
| | P_0203 | Guanylate kinase |
| | P_0881 | Uncharacterized MFS transporter |
| | P_0380 | Nicotinate (nicotinamide) nucleotide adenylyltransferase |
| | P_0001 | Chromosomal replication initiator protein |
| | P_0271 | tRNA N6-adenosine(37)-N6-threonylcarbamoyltransferase complex dimerization subunit |
| | P_0407 | RNA polymerase subunit sigma |
| | P_0045 | dTMP kinase |
| | P_0005 | Uncharacterized protein |
| | P_0616 | Fatty acid binding protein |
| | P_0428 | Phosphate transport system regulatory protein |
| | P_0548 | tRNA (cytidine(34)-2'-O)-methyltransferase |
| | P_0026 | Single-stranded DNA-binding protein |
| | P_0378 | NAD(+) synthase |
| | P_0371 | Flippase A |
| | P_0116 | Uncharacterized protein |
| | P_0686 | Potassium transporter |
| | P_0729 | Phosphoglycerate mutase (2,3-diphosphoglycerate-independent) |
| | P_0002 | DNA polymerase III subunit beta |

*continued on next page*

Table S2 – *continued from previous page*

| Module | Protein ID | Function |
|---|---|---|
| | P_0875 | CDP-diacylglycerol–glycerol-3-phosphate 3-phosphatidyltransferase |
| | P_0344 | Inorganic diphosphatase |
| | P_0859 | DNA topoisomerase I |
| | P_0081 | tRNA uridine(34) 5-carboxymethylaminomethyl synthesis GTPase |
| | P_0537 | UMP kinase |
| | P_0851 | Uncharacterized lipoprotein |
| Module 48 | P_0063 | Uncharacterized tRNA dihydrouridine synthase |
| | P_0115 | UTP–glucose-1-phosphate uridylyltransferase |
| | P_0650 | Type I methionyl aminopeptidase |
| | P_0833 | 50S ribosomal protein L9 |
| | P_0137 | 50S ribosomal protein L31 |
| | P_0640 | tRNA pseudouridine(38-40) synthase |
| | P_0615 | Uncharacterized protein |
| | P_0289 | Ribosome-binding factor A |
| | P_0431 | Uncharacterized metallophosphoesterase |
| | P_0813 | UDP-glucose 4-epimerase GalE |
| | P_0165 | Oligopeptide ABC transporter permease |
| | P_0403 | Ribosome GTPase |
| | P_0011 | Nucleoside ABC transporter substrate-binding protein |
| | P_0819 | Thioredoxin-disulfide reductase |
| | P_0082 | 30S ribosomal protein S20 |
| | P_0005 | Uncharacterized protein |
| | P_0825 | Excinuclease ABC subunit B |

*continued on next page*

Table S2 – *continued from previous page*

| Module | Protein ID | Function |
|---|---|---|
| | P_0271 | tRNA N6-adenosine(37)-N6-threonylcarbamoyltransferase complex dimerization subunit |
| | P_0387 | tRNA 2-thiouridine(34) synthase |
| | P_0592 | Uncharacterized protein |
| | P_0517 | Uncharacterized RNA pseudouridine synthase |
| | P_0203 | Guanylate kinase |
| | P_0821 | HPr(Ser) kinase/phosphatase |
| | P_0420 | Fatty acid kinase subunit A |
| | P_0195 | Spermidine/putrescine ABC transporter permease |
| | P_0168 | Oligopeptide ABC transporter ATP-binding protein |
| | P_0081 | tRNA uridine(34) 5-carboxymethylaminomethyl synthesis GTPase |
| | P_0391 | Elongation factor P |
| | P_0040 | tRNA lysidine(34) synthetase |
| Module 49 | P_0819 | Thioredoxin-disulfide reductase |
| | P_0330 | Deoxyguanosine kinase |
| | P_0684 | Bifunctional 5,10-methylene-tetrahydrofolate dehydrogenase/5,10-methylene-tetrahydrofolate cyclohydrolase |
| | P_0420 | Fatty acid kinase subunit A |
| | P_0133 | Toxin-antitoxin serine protease |
| | P_0218 | Glycerol kinase |
| | P_0696 | Uncharacterized transporter |

*continued on next page*

Table S2 – *continued from previous page*

| Module | Protein ID | Function |
|---|---|---|
| | P_0240 | tRNA 4-thiouridine(8) synthase |
| | P_0825 | Excinuclease ABC subunit B |
| | P_0065 | Thioredoxin |
| | P_0029 | FMN reductase |
| | P_0234 | PTS glucose transporter subunit IIA |
| | P_0800 | Ribose 5-phosphate isomerase B |
| | P_0203 | Guanylate kinase |
| | P_0348 | Ribosome biogenesis GTPase |
| | P_0639 | Uncharacterized efflux ABC transporter permease |
| | P_0294 | 30S ribosomal protein S15 |
| | P_0799 | Serine hydroxymethyltransferase |
| | P_0805 | Uncharacterized protein |
| | P_0114 | Glycolipid synthase B |
| | P_0109 | Deoxyribonuclease IV |
| | P_0302 | Flavin reductase |
| | P_0289 | Ribosome-binding factor A |
| | P_0544 | Heat-inducible transcription repressor |
| | P_0392 | Uncharacterized protein |
| | P_0601 | Uncharacterized protein |
| | P_0827 | Uncharacterized protein |
| | P_0378 | NAD(+) synthase |
| | P_0651 | Adenylate kinase |
| | P_0835 | Uncharacterized lipoprotein |
| | P_0710 | Low specificity hydrolase |
| | P_0377 | Ribosome GTPase |

*continued on next page*

Table S2 – *continued from previous page*

| Module | Protein ID | Function |
|---|---|---|
| | P_0728 | Low specificity hydrolase |
| | P_0661 | 50S ribosomal protein L14 |
| | P_0422 | 50S ribosomal protein L28 |
| | P_0821 | HPr(Ser) kinase/phosphatase |
| | P_0688 | Glutamyl-tRNA amidotransferase subunit A |
| | P_0733 | Phosphopentomutase |
| | P_0338 | Uncharacterized lipoprotein |
| | P_0779 | PTS sugar transporter |
| | P_0530 | Uncharacterized protein |
| | P_0271 | tRNA N6-adenosine(37)-N6-threonylcarbamoyltransferase complex dimerization subunit |
| | P_0838 | 23S rRNA (guanosine(2251)-2'-O)-methyltransferase |
| | P_0285 | Elongation factor 4 |
| | P_0329 | Uncharacterized pseudouridine synthase |
| | P_0200 | Translation initiation factor IF-3 |
| | P_0517 | Uncharacterized RNA pseudouridine synthase |
| | P_0511 | Uncharacterized protein |
| | P_0415 | Chromosome segregation protein |
| | P_0108 | Uncharacterized lipoprotein |
| | P_0794 | F0F1 ATP synthase subunit B |
| | P_0887 | Coenzyme A Disulfide Reductase |
| | P_0045 | dTMP kinase |
| | P_0042 | Uncharacterized transcriptional regulator |
| | P_0608 | Primosomal protein |

*continued on next page*

Table S2 – *continued from previous page*

| Module | Protein ID | Function |
|---|---|---|
| | P_0832 | Aminoacyl-tRNA hydrolase |
| | P_0823 | Dihydrofolate synthase |
| | P_0344 | Inorganic diphosphatase |
| | P_0166 | Oligopeptide ABC transporter permease |
| | P_0834 | Replicative DNA helicase |
| | P_0361 | 23S rRNA (pseudouridine(1915)-N3)-methyltransferase |
| | P_0537 | UMP kinase |
| Module 50 | P_0108 | Uncharacterized lipoprotein |
| | P_0140 | Thymidine kinase |
| | P_0109 | Deoxyribonuclease IV |
| | P_0257 | RNase J family beta-CASP ribonuclease |
| | P_0330 | Deoxyguanosine kinase |
| | P_0642 | ECF transporter ATPase |
| | P_0825 | Excinuclease ABC subunit B |
| | P_0610 | DNA-formamidopyrimidine glycosylase |
| | P_0776 | SsrA-binding protein |
| | P_0440 | Uncharacterized lipoprotein |
| | P_0834 | Replicative DNA helicase |
| | P_0065 | Thioredoxin |
| | P_0392 | Uncharacterized protein |
| | P_0824 | Excinuclease ABC subunit A |
| | P_0045 | dTMP kinase |
| | P_0639 | Uncharacterized efflux ABC transporter permease |
| | P_0137 | 50S ribosomal protein L31 |

*continued on next page*

Table S2 – *continued from previous page*

| Module | Protein ID | Function |
|---|---|---|
| | P_0789 | F0F1 ATP synthase subunit delta/epsilon |
| | P_0169 | Oligopeptide ABC transporter substrate-binding protein |
| | P_0615 | Uncharacterized protein |
| | P_0004 | 16S rRNA (adenine(1518)-N(6)/adenine(1519)-N(6))- dimethyltransferase |
| | P_0544 | Heat-inducible transcription repressor |
| | P_0081 | tRNA uridine(34) 5-carboxymethylaminomethyl synthesis GTPase |
| | P_0726 | Glucosamine-6-phosphate deaminase |
| | P_0294 | 30S ribosomal protein S15 |
| | P_0799 | Serine hydroxymethyltransferase |
| | P_0114 | Glycolipid synthase B |
| | P_0794 | F0F1 ATP synthase subunit B |
| | P_0511 | Uncharacterized protein |
| | P_0063 | Uncharacterized tRNA dihydrouridine synthase |
| | P_0875 | CDP-diacylglycerol–glycerol-3-phosphate 3-phosphatidyltransferase |
| | P_0419 | Phosphate acyltransferase |
| | P_0380 | Nicotinate (nicotinamide) nucleotide adenylyl-transferase |
| | P_0054 | Uncharacterized peroxiredoxin |
| | P_0301 | Ribosome assembly cofactor |
| | P_0168 | Oligopeptide ABC transporter ATP-binding protein |
| | P_0537 | UMP kinase |

*continued on next page*

Table S2 – *continued from previous page*

| Module | Protein ID | Function |
|---|---|---|
| | P_0262 | Ribulose-phosphate 3-epimerase |
| | P_0316 | Transketolase |
| | P_0145 | Uncharacterized acetyltransferase |
| | P_0197 | Spermidine/putrescine ABC transporter ATP-binding protein |
| | P_0431 | Uncharacterized metallophosphoesterase |
| | P_0617 | Fatty acid binding protein |
| | P_0228 | Dihydrolipoyl dehydrogenase |
| | P_0838 | 23S rRNA (guanosine(2251)-2'-O)-methyltransferase |
| | P_0592 | Uncharacterized protein |
| | P_0906 | Uncharacterized protein |
| | P_0636 | Uncharacterized lipoprotein |
| | P_0710 | Low specificity hydrolase |
| | P_0859 | DNA topoisomerase I |
| | P_0009 | Nucleoside ABC transporter permease |
| | P_0732 | Deoxyribose-phosphate aldolase |
| | P_0390 | Methionyl-tRNA formyltransferase |
| | P_0010 | Nucleoside ABC transporter ATP-binding protein |
| | P_0697 | Uncharacterized glycosyl transferase |

Table S3: Metabolite functional modules.

| **Module** | **Metabolite ID** | **Metabolite Name** |
|---|---|---|
| Module 1 | M_gdp_c | GDP |
| | M_pyr_c | Pyruvate |
| | M_2pg_c | D-Glycerate 2-phosphate |
| | M_ile__L_c | L-Isoleucine |
| | M_nh3_c | NH3 |
| | M_udp_c | UDP |
| | M_asp__L_c | L-Aspartate |
| | M_sprm_c | spermine |
| | M_nac_c | Nicotinate |
| | M_r1p_c | D-ribose 1-phosphate |
| | M_pep_c | Phosphoenolpyruvate |
| | M_acald_c | Acetaldehyde |
| | M_k_c | Potassium |
| | M_ser__L_c | L-Serine |
| | M_trp__L_c | L-Tryptophan |
| | M_glu__L_c | L-Glutamate |
| | M_galfur12dgr_c | Gal-DAG |
| | M_12dgr_c | 1,2-diacylglycerol |
| | M_prpp_c | 5-Phospho-alpha-D-ribose 1-diphosphate |
| | M_cdpdag_c | CDP-diacylglycerol |
| Module 2 | M_dcyt_c | Deoxycytidine |
| | M_cmp_c | CMP |
| | M_dcmp_c | DCMP |
| | M_nadph_c | NADPH |
| | M_dtdp_c | DTDP |
| | M_lystrna_c | L-lysyl-tRNA(Lys) |

*continued on next page*

Table S3 – *continued from previous page*

| **Module** | **Metabolite ID** | **Metabolite Name** |
|---|---|---|
| | M_dcdp_c | DCDP |
| | M_actp_c | Acetyl phosphate |
| | M_sprm_c | spermine |
| | M_asn__L_c | L-Asparagine |
| | M_2pg_c | D-Glycerate 2-phosphate |
| | M_e4p_c | D-Erythrose 4-phosphate |
| | M_dump_c | DUMP |
| | M_pi_c | Phosphate |
| | M_ile__L_c | L-Isoleucine |
| | M_ppi_c | Diphosphate |
| | M_f6p_c | D-Fructose 6-phosphate |
| | M_dctp_c | DCTP |
| | M_ser__L_c | L-Serine |
| | M_gsn_c | Guanosine |
| Module 3 | M_lys__L_c | L-Lysine |
| | M_trnalys_c | TRNA(Lys) |
| | M_chsterol_c | cholesterol |
| | M_dcmp_c | DCMP |
| | M_clpn_c | cardiolipin |
| | M_gly_c | Glycine |
| | M_lac__L_c | L-Lactate |
| | M_tyr__L_c | L-Tyrosine |
| | M_f6p_c | D-Fructose 6-phosphate |
| | M_ile__L_c | L-Isoleucine |
| | M_pap_c | Adenosine 3',5'-bisphosphate |
| | M_ppi_c | Diphosphate |

*continued on next page*

Table S3 – *continued from previous page*

| **Module** | **Metabolite ID** | **Metabolite Name** |
| --- | --- | --- |
| | M_dump_c | DUMP |
| | M_nadp_c | NADP+ |
| | M_glyc3p_c | Glycerol 3-phosphate |
| | M_cmp_c | CMP |
| | M_val__L_c | L-Valine |
| | M_coa_c | CoA |
| | M_pyr_c | Pyruvate |
| | M_met__L_c | L-Methionine |
| Module 4 | M_ile__L_c | L-Isoleucine |
| | M_cys__L_c | L-Cysteine |
| | M_ala__L_c | L-Alanine |
| | M_val__L_c | L-Valine |
| | M_glyc3p_c | Glycerol 3-phosphate |
| | M_prpp_c | 5-Phospho-alpha-D-ribose 1-diphosphate |
| | M_2dr5p_c | 2-Deoxy-D-ribose 5-phosphate |
| | M_gln__L_c | L-Glutamine |
| | M_nad_c | NAD+ |
| | M_pg_c | phosphatidylglycerol |
| | M_sprm_c | spermine |
| | M_3pg_c | 3-Phospho-D-glycerate |
| | M_pc_c | Phosphatidylcholine |
| | M_adn_c | Adenosine |
| | M_nac_c | Nicotinate |
| | M_xu5p__D_c | D-Xylulose 5-phosphate |
| | M_ump_c | UMP |
| | M_cdpdag_c | CDP-diacylglycerol |

*continued on next page*

Table S3 – *continued from previous page*

| Module | Metabolite ID | Metabolite Name |
|---|---|---|
| | M_nh3_c | NH3 |
| | M_g1p_c | D-Glucose 1-phosphate |
| Module 5 | M_ala__L_c | L-Alanine |
| | M_pc_c | Phosphatidylcholine |
| | M_clpn_c | cardiolipin |
| | M_gsn_c | Guanosine |
| | M_leu__L_c | L-Leucine |
| | M_cys__L_c | L-Cysteine |
| | M_g1p_c | D-Glucose 1-phosphate |
| | M_tyr__L_c | L-Tyrosine |
| | M_dcdp_c | DCDP |
| | M_gtp_c | GTP |
| | M_lystrna_c | L-lysyl-tRNA(Lys) |
| | M_adn_c | Adenosine |
| | M_chsterol_c | cholesterol |
| | M_13dpg_c | 3-Phospho-D-glyceroyl phosphate |
| | M_pap_c | Adenosine 3',5'-bisphosphate |
| | M_ura_c | Uracil |
| | M_ump_c | UMP |
| | M_tyrtrna_c | L-Tyrosyl-tRNA(Tyr) |
| | M_ser__L_c | L-Serine |
| | M_asn__L_c | L-Asparagine |
| Module 6 | M_dgtp_c | DGTP |
| | M_datp_c | DATP |
| | M_1ag3p_c | 1-acyl-glycerol 3-phosphate |
| | M_dttp_c | DTTP |

*continued on next page*

Table S3 – *continued from previous page*

| **Module** | **Metabolite ID** | **Metabolite Name** |
|---|---|---|
| | M_g1p_c | D-Glucose 1-phosphate |
| | M_trnalys_c | TRNA(Lys) |
| | M_gdp_c | GDP |
| | M_adp_c | ADP |
| | M_gtp_c | GTP |
| | M_pa_c | phosphatidic acid |
| | M_coa_c | CoA |
| | M_dctp_c | DCTP |
| | M_r5p_c | D-ribose 5-phosphate |
| | M_3pg_c | 3-Phospho-D-glycerate |
| | M_ump_c | UMP |
| | M_dtmp_c | DTMP |
| | M_10fthfglu3_c | 10-formyltetrahydrofolate tri-L-glutamate |
| | M_amp_c | AMP |
| | M_pep_c | Phosphoenolpyruvate |
| | M_tag_c | triacylglycerol |
| Module 7 | M_ac_c | Acetate |
| | M_dump_c | DUMP |
| | M_ump_c | UMP |
| | M_thmpp_c | thiamin diphosphate |
| | M_clpn_c | cardiolipin |
| | M_udpgalfur_c | UDP-galactofuranose |
| | M_nadp_c | NADP+ |
| | M_cmp_c | CMP |
| | M_r5p_c | D-ribose 5-phosphate |
| | M_e4p_c | D-Erythrose 4-phosphate |

*continued on next page*

Table S3 – *continued from previous page*

| Module | Metabolite ID | Metabolite Name |
|---|---|---|
| | M_dtdp_c | DTDP |
| | M_ile__L_c | L-Isoleucine |
| | M_coa_c | CoA |
| | M_gln__L_c | L-Glutamine |
| | M_tyr__L_c | L-Tyrosine |
| | M_ade_c | Adenine |
| | M_dgtp_c | DGTP |
| | M_glyc3p_c | Glycerol 3-phosphate |
| | M_pa_c | phosphatidic acid |
| | M_10fthfglu3_c | 10-formyltetrahydrofolate tri-L-glutamate |
| Module 8 | M_cdp_c | CDP |
| | M_chsterol_c | cholesterol |
| | M_atp_c | ATP |
| | M_gsn_c | Guanosine |
| | M_g3p_c | Glyceraldehyde 3-phosphate |
| | M_dtdp_c | DTDP |
| | M_s7p_c | Sedoheptulose 7-phosphate |
| | M_ump_c | UMP |
| | M_dcdp_c | DCDP |
| | M_adn_c | Adenosine |
| | M_1ag3p_c | 1-acyl-glycerol 3-phosphate |
| | M_f6p_c | D-Fructose 6-phosphate |
| | M_gly_c | Glycine |
| | M_pi_c | Phosphate |
| | M_k_c | Potassium |
| | M_coa_c | CoA |

*continued on next page*

Table S3 – *continued from previous page*

| Module | Metabolite ID | Metabolite Name |
|---|---|---|
| | M_r1p_c | D-ribose 1-phosphate |
| | M_pc_c | Phosphatidylcholine |
| | M_gdp_c | GDP |
| | M_dtmp_c | DTMP |
| Module 9 | M_nadph_c | NADPH |
| | M_ppi_c | Diphosphate |
| | M_dump_c | DUMP |
| | M_udpgalfur_c | UDP-galactofuranose |
| | M_val__L_c | L-Valine |
| | M_gly_c | Glycine |
| | M_clpn_c | cardiolipin |
| | M_pap_c | Adenosine 3',5'-bisphosphate |
| | M_mg2_c | Magnesium |
| | M_cmp_c | CMP |
| | M_dctp_c | DCTP |
| | M_dgtp_c | DGTP |
| | M_datp_c | DATP |
| | M_ile__L_c | L-Isoleucine |
| | M_trp__L_c | L-Tryptophan |
| | M_nadp_c | NADP+ |
| | M_trnalys_c | TRNA(Lys) |
| | M_cys__L_c | L-Cysteine |
| | M_12dgr_c | 1,2-diacylglycerol |
| | M_sm_c | Sphingomyelin |
| Module 10 | M_gly_c | Glycine |
| | M_fdp_c | D-Fructose 1,6-bisphosphate |

*continued on next page*

Table S3 – *continued from previous page*

| Module | Metabolite ID | Metabolite Name |
|---|---|---|
| | M_ppi_c | Diphosphate |
| | M_g1p_c | D-Glucose 1-phosphate |
| | M_dctp_c | DCTP |
| | M_cmp_c | CMP |
| | M_clpn_c | cardiolipin |
| | M_sprm_c | spermine |
| | M_ile__L_c | L-Isoleucine |
| | M_lystrna_c | L-lysyl-tRNA(Lys) |
| | M_coa_c | CoA |
| | M_trnalys_c | TRNA(Lys) |
| | M_galfur12dgr_c | Gal-DAG |
| | M_acald_c | Acetaldehyde |
| | M_pyr_c | Pyruvate |
| | M_uri_c | Uridine |
| | M_pc_c | Phosphatidylcholine |
| | M_mg2_c | Magnesium |
| | M_xu5p__D_c | D-Xylulose 5-phosphate |
| | M_tyr__L_c | L-Tyrosine |
| Module 11 | M_xu5p__D_c | D-Xylulose 5-phosphate |
| | M_glyc3p_c | Glycerol 3-phosphate |
| | M_pyr_c | Pyruvate |
| | M_sprm_c | spermine |
| | M_ile__L_c | L-Isoleucine |
| | M_adp_c | ADP |
| | M_r5p_c | D-ribose 5-phosphate |
| | M_nh3_c | NH3 |

*continued on next page*

Table S3 – *continued from previous page*

| **Module** | **Metabolite ID** | **Metabolite Name** |
|---|---|---|
| | M_ura_c | Uracil |
| | M_dtmp_c | DTMP |
| | M_adn_c | Adenosine |
| | M_glu__L_c | L-Glutamate |
| | M_ru5p__D_c | D-Ribulose 5-phosphate |
| | M_coa_c | CoA |
| | M_12dgr_c | 1,2-diacylglycerol |
| | M_cdpdag_c | CDP-diacylglycerol |
| | M_1ag3p_c | 1-acyl-glycerol 3-phosphate |
| | M_nac_c | Nicotinate |
| | M_datp_c | DATP |
| | M_pg_c | phosphatidylglycerol |
| Module 12 | M_ppi_c | Diphosphate |
| | M_gsn_c | Guanosine |
| | M_pa_c | phosphatidic acid |
| | M_glyc3p_c | Glycerol 3-phosphate |
| | M_amp_c | AMP |
| | M_xu5p__D_c | D-Xylulose 5-phosphate |
| | M_dttp_c | DTTP |
| | M_uri_c | Uridine |
| | M_cdpdag_c | CDP-diacylglycerol |
| | M_udpg_c | UDP-glucose |
| | M_galfur12dgr_c | Gal-DAG |
| | M_dctp_c | DCTP |
| | M_mg2_c | Magnesium |
| | M_s7p_c | Sedoheptulose 7-phosphate |

*continued on next page*

Table S3 – *continued from previous page*

| Module | Metabolite ID | Metabolite Name |
|---|---|---|
| | M_nad_c | NAD+ |
| | M_accoa_c | Acetyl-CoA |
| | M_g3p_c | Glyceraldehyde 3-phosphate |
| | M_fmn_c | FMN |
| | M_lac__L_c | L-Lactate |
| | M_ile__L_c | L-Isoleucine |
| Module 13 | M_r5p_c | D-ribose 5-phosphate |
| | M_10fthfglu3_c | 10-formyltetrahydrofolate tri-L-glutamate |
| | M_13dpg_c | 3-Phospho-D-glyceroyl phosphate |
| | M_g6p_c | D-Glucose 6-phosphate |
| | M_nac_c | Nicotinate |
| | M_cmp_c | CMP |
| | M_xu5p__D_c | D-Xylulose 5-phosphate |
| | M_adp_c | ADP |
| | M_ser__L_c | L-Serine |
| | M_udpg_c | UDP-glucose |
| | M_prpp_c | 5-Phospho-alpha-D-ribose 1-diphosphate |
| | M_asp__L_c | L-Aspartate |
| | M_2pg_c | D-Glycerate 2-phosphate |
| | M_ile__L_c | L-Isoleucine |
| | M_ade_c | Adenine |
| | M_adn_c | Adenosine |
| | M_amp_c | AMP |
| | M_dtdp_c | DTDP |
| | M_s7p_c | Sedoheptulose 7-phosphate |
| | M_ctp_c | CTP |

*continued on next page*

Table S3 – *continued from previous page*

| Module | Metabolite ID | Metabolite Name |
|---|---|---|
| Module 14 | M_arg__L_c | L-Arginine |
| | M_5fthfglu3_c | 5-formyltetrahydrofolate tri-L-glutamate |
| | M_clpn_c | cardiolipin |
| | M_dump_c | DUMP |
| | M_dtdp_c | DTDP |
| | M_atp_c | ATP |
| | M_udp_c | UDP |
| | M_ru5p__D_c | D-Ribulose 5-phosphate |
| | M_ump_c | UMP |
| | M_pap_c | Adenosine 3',5'-bisphosphate |
| | M_prpp_c | 5-Phospho-alpha-D-ribose 1-diphosphate |
| | M_pc_c | Phosphatidylcholine |
| | M_phe__L_c | L-Phenylalanine |
| | M_ser__L_c | L-Serine |
| | M_g1p_c | D-Glucose 1-phosphate |
| | M_pro__L_c | L-Proline |
| | M_trp__L_c | L-Tryptophan |
| | M_e4p_c | D-Erythrose 4-phosphate |
| | M_asp__L_c | L-Aspartate |
| | M_fdp_c | D-Fructose 1,6-bisphosphate |
| Module 15 | M_cdp_c | CDP |
| | M_dcdp_c | DCDP |
| | M_nadph_c | NADPH |
| | M_his__L_c | L-Histidine |
| | M_thr__L_c | L-Threonine |
| | M_chsterol_c | cholesterol |

*continued on next page*

Table S3 – *continued from previous page*

| **Module** | **Metabolite ID** | **Metabolite Name** |
|---|---|---|
| | M_xu5p__D_c | D-Xylulose 5-phosphate |
| | M_ump_c | UMP |
| | M_val__L_c | L-Valine |
| | M_trptrna_c | L-Tryptophanyl-tRNA(Trp) |
| | M_sprm_c | spermine |
| | M_clpn_c | cardiolipin |
| | M_leu__L_c | L-Leucine |
| | M_gsn_c | Guanosine |
| | M_g1p_c | D-Glucose 1-phosphate |
| | M_f6p_c | D-Fructose 6-phosphate |
| | M_dctp_c | DCTP |
| | M_1ag3p_c | 1-acyl-glycerol 3-phosphate |
| | M_glyc3p_c | Glycerol 3-phosphate |
| | M_asp__L_c | L-Aspartate |
| Module 16 | M_g1p_c | D-Glucose 1-phosphate |
| | M_xu5p__D_c | D-Xylulose 5-phosphate |
| | M_ura_c | Uracil |
| | M_gdp_c | GDP |
| | M_pyr_c | Pyruvate |
| | M_adp_c | ADP |
| | M_nh3_c | NH3 |
| | M_adn_c | Adenosine |
| | M_pep_c | Phosphoenolpyruvate |
| | M_dctp_c | DCTP |
| | M_ile__L_c | L-Isoleucine |
| | M_accoa_c | Acetyl-CoA |

*continued on next page*

Table S3 – *continued from previous page*

| Module | Metabolite ID | Metabolite Name |
|---|---|---|
| | M_lystrna_c | L-lysyl-tRNA(Lys) |
| | M_coa_c | CoA |
| | M_ser__L_c | L-Serine |
| | M_amp_c | AMP |
| | M_12dgr_c | 1,2-diacylglycerol |
| | M_k_c | Potassium |
| | M_trp__L_c | L-Tryptophan |
| | M_udpg_c | UDP-glucose |
| Module 17 | M_cystrna_c | L-Cysteinyl-tRNA(Cys) |
| | M_nicrnt_c | Nicotinate D-ribonucleotide |
| | M_dump_c | DUMP |
| | M_2pg_c | D-Glycerate 2-phosphate |
| | M_actp_c | Acetyl phosphate |
| | M_val__L_c | L-Valine |
| | M_dcmp_c | DCMP |
| | M_trnalys_c | TRNA(Lys) |
| | M_nadp_c | NADP+ |
| | M_pc_c | Phosphatidylcholine |
| | M_chsterol_c | cholesterol |
| | M_ppi_c | Diphosphate |
| | M_1ag3p_c | 1-acyl-glycerol 3-phosphate |
| | M_udpgalfur_c | UDP-galactofuranose |
| | M_dctp_c | DCTP |
| | M_pep_c | Phosphoenolpyruvate |
| | M_thr__L_c | L-Threonine |
| | M_arg__L_c | L-Arginine |

*continued on next page*

Table S3 – *continued from previous page*

| **Module** | **Metabolite ID** | **Metabolite Name** |
|---|---|---|
| | M_nadph_c | NADPH |
| | M_mg2_c | Magnesium |
| Module 18 | M_cystrna_c | L-Cysteinyl-tRNA(Cys) |
| | M_ru5p__D_c | D-Ribulose 5-phosphate |
| | M_nicrnt_c | Nicotinate D-ribonucleotide |
| | M_sm_c | Sphingomyelin |
| | M_lpl_PdhC_c | lipoyl-PdhC |
| | M_trnalys_c | TRNA(Lys) |
| | M_gtp_c | GTP |
| | M_nadp_c | NADP+ |
| | M_chsterol_c | cholesterol |
| | M_glyc3p_c | Glycerol 3-phosphate |
| | M_pap_c | Adenosine 3',5'-bisphosphate |
| | M_gly_c | Glycine |
| | M_g6p_c | D-Glucose 6-phosphate |
| | M_alatrna_c | L-Alanyl-tRNA(Ala) |
| | M_datp_c | DATP |
| | M_gln__L_c | L-Glutamine |
| | M_udpgalfur_c | UDP-galactofuranose |
| | M_cys__L_c | L-Cysteine |
| | M_cdp_c | CDP |
| | M_ile__L_c | L-Isoleucine |
| Module 19 | M_amp_c | AMP |
| | M_dtdp_c | DTDP |
| | M_nac_c | Nicotinate |
| | M_12dgr_c | 1,2-diacylglycerol |

*continued on next page*

Table S3 – *continued from previous page*

| Module | Metabolite ID | Metabolite Name |
|---|---|---|
| | M_ade_c | Adenine |
| | M_mg2_c | Magnesium |
| | M_nad_c | NAD+ |
| | M_2dr5p_c | 2-Deoxy-D-ribose 5-phosphate |
| | M_2pg_c | D-Glycerate 2-phosphate |
| | M_gmp_c | GMP |
| | M_pa_c | phosphatidic acid |
| | M_udpg_c | UDP-glucose |
| | M_r1p_c | D-ribose 1-phosphate |
| | M_glu__L_c | L-Glutamate |
| | M_ca2_c | Calcium |
| | M_fa_c | fatty acid |
| | M_pg_c | phosphatidylglycerol |
| | M_asp__L_c | L-Aspartate |
| | M_galfur12dgr_c | Gal-DAG |
| | M_gln__L_c | L-Glutamine |
| Module 20 | M_chsterol_c | cholesterol |
| | M_dcdp_c | DCDP |
| | M_sm_c | Sphingomyelin |
| | M_dtdp_c | DTDP |
| | M_cdp_c | CDP |
| | M_his__L_c | L-Histidine |
| | M_arg__L_c | L-Arginine |
| | M_trnaasn_c | TRNA(Asn) |
| | M_pc_c | Phosphatidylcholine |
| | M_ppi_c | Diphosphate |

*continued on next page*

Table S3 – *continued from previous page*

| **Module** | **Metabolite ID** | **Metabolite Name** |
|---|---|---|
| | M_trnalys_c | TRNA(Lys) |
| | M_gua_c | Guanine |
| | M_ala__L_c | L-Alanine |
| | M_gmp_c | GMP |
| | M_gtp_c | GTP |
| | M_f6p_c | D-Fructose 6-phosphate |
| | M_datp_c | DATP |
| | M_xu5p__D_c | D-Xylulose 5-phosphate |
| | M_trnagln_c | TRNA(Gln) |
| | M_1ag3p_c | 1-acyl-glycerol 3-phosphate |
| Module 21 | M_udpgalfur_c | UDP-galactofuranose |
| | M_lys__L_c | L-Lysine |
| | M_phe__L_c | L-Phenylalanine |
| | M_ser__L_c | L-Serine |
| | M_adn_c | Adenosine |
| | M_lac__L_c | L-Lactate |
| | M_clpn_c | cardiolipin |
| | M_udpg_c | UDP-glucose |
| | M_g6p_c | D-Glucose 6-phosphate |
| | M_ppi_c | Diphosphate |
| | M_pyr_c | Pyruvate |
| | M_cdpdag_c | CDP-diacylglycerol |
| | M_ca2_c | Calcium |
| | M_r1p_c | D-ribose 1-phosphate |
| | M_gsn_c | Guanosine |
| | M_acald_c | Acetaldehyde |

*continued on next page*

Table S3 – *continued from previous page*

| Module | Metabolite ID | Metabolite Name |
|---|---|---|
| | M_utp_c | UTP |
| | M_gdp_c | GDP |
| | M_3pg_c | 3-Phospho-D-glycerate |
| | M_pa_c | phosphatidic acid |
| Module 22 | M_gln__L_c | L-Glutamine |
| | M_uri_c | Uridine |
| | M_k_c | Potassium |
| | M_glyc3p_c | Glycerol 3-phosphate |
| | M_amp_c | AMP |
| | M_adp_c | ADP |
| | M_ura_c | Uracil |
| | M_dtmp_c | DTMP |
| | M_dctp_c | DCTP |
| | M_asp__L_c | L-Aspartate |
| | M_2pg_c | D-Glycerate 2-phosphate |
| | M_pyr_c | Pyruvate |
| | M_fa_c | fatty acid |
| | M_cdpdag_c | CDP-diacylglycerol |
| | M_g1p_c | D-Glucose 1-phosphate |
| | M_r5p_c | D-ribose 5-phosphate |
| | M_phe__L_c | L-Phenylalanine |
| | M_datp_c | DATP |
| | M_ca2_c | Calcium |
| | M_atp_c | ATP |
| Module 23 | M_gdp_c | GDP |
| | M_g1p_c | D-Glucose 1-phosphate |

*continued on next page*

Table S3 – *continued from previous page*

| **Module** | **Metabolite ID** | **Metabolite Name** |
|---|---|---|
| | M_ade_c | Adenine |
| | M_pep_c | Phosphoenolpyruvate |
| | M_dtdp_c | DTDP |
| | M_cmp_c | CMP |
| | M_nad_c | NAD+ |
| | M_fdp_c | D-Fructose 1,6-bisphosphate |
| | M_glu__L_c | L-Glutamate |
| | M_utp_c | UTP |
| | M_coa_c | CoA |
| | M_nadp_c | NADP+ |
| | M_dtmp_c | DTMP |
| | M_sprm_c | spermine |
| | M_asp__L_c | L-Aspartate |
| | M_k_c | Potassium |
| | M_pa_c | phosphatidic acid |
| | M_2pg_c | D-Glycerate 2-phosphate |
| | M_ca2_c | Calcium |
| | M_cdpdag_c | CDP-diacylglycerol |
| Module 24 | M_ru5p__D_c | D-Ribulose 5-phosphate |
| | M_trnalys_c | TRNA(Lys) |
| | M_sm_c | Sphingomyelin |
| | M_nadph_c | NADPH |
| | M_nadp_c | NADP+ |
| | M_pc_c | Phosphatidylcholine |
| | M_dump_c | DUMP |
| | M_nac_c | Nicotinate |

*continued on next page*

Table S3 – *continued from previous page*

| Module | Metabolite ID | Metabolite Name |
|---|---|---|
| | M_met__L_c | L-Methionine |
| | M_his__L_c | L-Histidine |
| | M_val__L_c | L-Valine |
| | M_thr__L_c | L-Threonine |
| | M_glyc3p_c | Glycerol 3-phosphate |
| | M_dcmp_c | DCMP |
| | M_pro__L_c | L-Proline |
| | M_gln__L_c | L-Glutamine |
| | M_nad_c | NAD+ |
| | M_ac_c | Acetate |
| | M_dtdp_c | DTDP |
| | M_chsterol_c | cholesterol |
| Module 25 | M_nadph_c | NADPH |
| | M_cystrna_c | L-Cysteinyl-tRNA(Cys) |
| | M_gly_c | Glycine |
| | M_sm_c | Sphingomyelin |
| | M_pc_c | Phosphatidylcholine |
| | M_ser__L_c | L-Serine |
| | M_trnalys_c | TRNA(Lys) |
| | M_met__L_c | L-Methionine |
| | M_2pg_c | D-Glycerate 2-phosphate |
| | M_sprm_c | spermine |
| | M_g1p_c | D-Glucose 1-phosphate |
| | M_thr__L_c | L-Threonine |
| | M_cmp_c | CMP |
| | M_cdpdag_c | CDP-diacylglycerol |

*continued on next page*

Table S3 – *continued from previous page*

| Module | Metabolite ID | Metabolite Name |
|---|---|---|
| | M_arg__L_c | L-Arginine |
| | M_fmn_c | FMN |
| | M_e4p_c | D-Erythrose 4-phosphate |
| | M_accoa_c | Acetyl-CoA |
| | M_13dpg_c | 3-Phospho-D-glyceroyl phosphate |
| | M_f6p_c | D-Fructose 6-phosphate |
| Module 26 | M_ru5p__D_c | D-Ribulose 5-phosphate |
| | M_dtdp_c | DTDP |
| | M_dcdp_c | DCDP |
| | M_ala__L_c | L-Alanine |
| | M_dump_c | DUMP |
| | M_2pg_c | D-Glycerate 2-phosphate |
| | M_gly_c | Glycine |
| | M_cys__L_c | L-Cysteine |
| | M_prpp_c | 5-Phospho-alpha-D-ribose 1-diphosphate |
| | M_g6p_c | D-Glucose 6-phosphate |
| | M_actp_c | Acetyl phosphate |
| | M_dgtp_c | DGTP |
| | M_trnalys_c | TRNA(Lys) |
| | M_pg_c | phosphatidylglycerol |
| | M_nadp_c | NADP+ |
| | M_pc_c | Phosphatidylcholine |
| | M_pep_c | Phosphoenolpyruvate |
| | M_pi_c | Phosphate |
| | M_trp__L_c | L-Tryptophan |
| | M_glyc3p_c | Glycerol 3-phosphate |

*continued on next page*

Table S3 – *continued from previous page*

| Module | Metabolite ID | Metabolite Name |
|---|---|---|
| Module 27 | M_alatrna_c | L-Alanyl-tRNA(Ala) |
| | M_trnalys_c | TRNA(Lys) |
| | M_gly_c | Glycine |
| | M_clpn_c | cardiolipin |
| | M_met__L_c | L-Methionine |
| | M_udpgalfur_c | UDP-galactofuranose |
| | M_dump_c | DUMP |
| | M_pc_c | Phosphatidylcholine |
| | M_trnagln_c | TRNA(Gln) |
| | M_coa_c | CoA |
| | M_1ag3p_c | 1-acyl-glycerol 3-phosphate |
| | M_13dpg_c | 3-Phospho-D-glyceroyl phosphate |
| | M_nadp_c | NADP+ |
| | M_ump_c | UMP |
| | M_adn_c | Adenosine |
| | M_asp__L_c | L-Aspartate |
| | M_ru5p__D_c | D-Ribulose 5-phosphate |
| | M_cdp_c | CDP |
| | M_cmp_c | CMP |
| | M_nac_c | Nicotinate |
| Module 28 | M_ump_c | UMP |
| | M_ppi_c | Diphosphate |
| | M_fmn_c | FMN |
| | M_pc_c | Phosphatidylcholine |
| | M_ctp_c | CTP |
| | M_glyc3p_c | Glycerol 3-phosphate |

*continued on next page*

Table S3 – *continued from previous page*

| Module | Metabolite ID | Metabolite Name |
|---|---|---|
| | M_k_c | Potassium |
| | M_prpp_c | 5-Phospho-alpha-D-ribose 1-diphosphate |
| | M_pep_c | Phosphoenolpyruvate |
| | M_asn__L_c | L-Asparagine |
| | M_pa_c | phosphatidic acid |
| | M_acald_c | Acetaldehyde |
| | M_utp_c | UTP |
| | M_dtdp_c | DTDP |
| | M_1ag3p_c | 1-acyl-glycerol 3-phosphate |
| | M_dctp_c | DCTP |
| | M_2dr5p_c | 2-Deoxy-D-ribose 5-phosphate |
| | M_s7p_c | Sedoheptulose 7-phosphate |
| | M_ser__L_c | L-Serine |
| | M_dgtp_c | DGTP |
| Module 29 | M_pap_c | Adenosine 3',5'-bisphosphate |
| | M_ser__L_c | L-Serine |
| | M_gmp_c | GMP |
| | M_pc_c | Phosphatidylcholine |
| | M_pi_c | Phosphate |
| | M_ade_c | Adenine |
| | M_cmp_c | CMP |
| | M_cdpdag_c | CDP-diacylglycerol |
| | M_trnalys_c | TRNA(Lys) |
| | M_dctp_c | DCTP |
| | M_coa_c | CoA |
| | M_prpp_c | 5-Phospho-alpha-D-ribose 1-diphosphate |

*continued on next page*

Table S3 – *continued from previous page*

| **Module** | **Metabolite ID** | **Metabolite Name** |
|---|---|---|
| | M_gdp_c | GDP |
| | M_fa_c | fatty acid |
| | M_lac__L_c | L-Lactate |
| | M_g6p_c | D-Glucose 6-phosphate |
| | M_12dgr_c | 1,2-diacylglycerol |
| | M_pep_c | Phosphoenolpyruvate |
| | M_leu__L_c | L-Leucine |
| | M_ala__L_c | L-Alanine |
| Module 30 | M_gly_c | Glycine |
| | M_gua_c | Guanine |
| | M_ump_c | UMP |
| | M_ala__L_c | L-Alanine |
| | M_trnagln_c | TRNA(Gln) |
| | M_pro__L_c | L-Proline |
| | M_gmp_c | GMP |
| | M_arg__L_c | L-Arginine |
| | M_k_c | Potassium |
| | M_trnaile_c | TRNA(Ile) |
| | M_clpn_c | cardiolipin |
| | M_trnalys_c | TRNA(Lys) |
| | M_e4p_c | D-Erythrose 4-phosphate |
| | M_atp_c | ATP |
| | M_ca2_c | Calcium |
| | M_chsterol_c | cholesterol |
| | M_ctp_c | CTP |
| | M_mg2_c | Magnesium |

*continued on next page*

Table S3 – *continued from previous page*

| **Module** | **Metabolite ID** | **Metabolite Name** |
|---|---|---|
| | M_udpg_c | UDP-glucose |
| | M_acald_c | Acetaldehyde |
| Module 31 | M_asp__L_c | L-Aspartate |
| | M_mg2_c | Magnesium |
| | M_3pg_c | 3-Phospho-D-glycerate |
| | M_pi_c | Phosphate |
| | M_glu__L_c | L-Glutamate |
| | M_udpg_c | UDP-glucose |
| | M_ile__L_c | L-Isoleucine |
| | M_fdp_c | D-Fructose 1,6-bisphosphate |
| | M_dtdp_c | DTDP |
| | M_pro__L_c | L-Proline |
| | M_k_c | Potassium |
| | M_lystrna_c | L-lysyl-tRNA(Lys) |
| | M_datp_c | DATP |
| | M_adn_c | Adenosine |
| | M_udp_c | UDP |
| | M_2pg_c | D-Glycerate 2-phosphate |
| | M_accoa_c | Acetyl-CoA |
| | M_glyc3p_c | Glycerol 3-phosphate |
| | M_dttp_c | DTTP |
| | M_acald_c | Acetaldehyde |
| Module 32 | M_udpgalfur_c | UDP-galactofuranose |
| | M_k_c | Potassium |
| | M_clpn_c | cardiolipin |
| | M_cdp_c | CDP |

*continued on next page*

Table S3 – *continued from previous page*

| **Module** | **Metabolite ID** | **Metabolite Name** |
|---|---|---|
| | M_prpp_c | 5-Phospho-alpha-D-ribose 1-diphosphate |
| | M_leu__L_c | L-Leucine |
| | M_2pg_c | D-Glycerate 2-phosphate |
| | M_nadp_c | NADP+ |
| | M_gln__L_c | L-Glutamine |
| | M_s7p_c | Sedoheptulose 7-phosphate |
| | M_datp_c | DATP |
| | M_sprm_c | spermine |
| | M_fdp_c | D-Fructose 1,6-bisphosphate |
| | M_13dpg_c | 3-Phospho-D-glyceroyl phosphate |
| | M_mg2_c | Magnesium |
| | M_3pg_c | 3-Phospho-D-glycerate |
| | M_coa_c | CoA |
| | M_adp_c | ADP |
| | M_pap_c | Adenosine 3',5'-bisphosphate |
| | M_dcdp_c | DCDP |
| Module 33 | M_ura_c | Uracil |
| | M_r1p_c | D-ribose 1-phosphate |
| | M_nh3_c | NH3 |
| | M_pa_c | phosphatidic acid |
| | M_uri_c | Uridine |
| | M_dttp_c | DTTP |
| | M_datp_c | DATP |
| | M_ile__L_c | L-Isoleucine |
| | M_ade_c | Adenine |
| | M_udpgalfur_c | UDP-galactofuranose |

*continued on next page*

Table S3 – *continued from previous page*

| Module | Metabolite ID | Metabolite Name |
|---|---|---|
| | M_fdp_c | D-Fructose 1,6-bisphosphate |
| | M_gdp_c | GDP |
| | M_12dgr_c | 1,2-diacylglycerol |
| | M_trp__L_c | L-Tryptophan |
| | M_utp_c | UTP |
| | M_coa_c | CoA |
| | M_glu__L_c | L-Glutamate |
| | M_dctp_c | DCTP |
| | M_nad_c | NAD+ |
| | M_cdpdag_c | CDP-diacylglycerol |
| Module 34 | M_accoa_c | Acetyl-CoA |
| | M_ura_c | Uracil |
| | M_coa_c | CoA |
| | M_amp_c | AMP |
| | M_2dr5p_c | 2-Deoxy-D-ribose 5-phosphate |
| | M_r5p_c | D-ribose 5-phosphate |
| | M_pa_c | phosphatidic acid |
| | M_10fthfglu3_c | 10-formyltetrahydrofolate tri-L-glutamate |
| | M_pep_c | Phosphoenolpyruvate |
| | M_dgtp_c | DGTP |
| | M_udp_c | UDP |
| | M_pg_c | phosphatidylglycerol |
| | M_ppi_c | Diphosphate |
| | M_g1p_c | D-Glucose 1-phosphate |
| | M_ctp_c | CTP |
| | M_uri_c | Uridine |

*continued on next page*

Table S3 – *continued from previous page*

| Module | Metabolite ID | Metabolite Name |
|---|---|---|
| | M_gdp_c | GDP |
| | M_prpp_c | 5-Phospho-alpha-D-ribose 1-diphosphate |
| | M_ser__L_c | L-Serine |
| | M_mg2_c | Magnesium |
| Module 35 | M_g1p_c | D-Glucose 1-phosphate |
| | M_coa_c | CoA |
| | M_leu__L_c | L-Leucine |
| | M_dtmp_c | DTMP |
| | M_ade_c | Adenine |
| | M_pap_c | Adenosine 3',5'-bisphosphate |
| | M_ser__L_c | L-Serine |
| | M_r1p_c | D-ribose 1-phosphate |
| | M_asp__L_c | L-Aspartate |
| | M_uri_c | Uridine |
| | M_ctp_c | CTP |
| | M_pep_c | Phosphoenolpyruvate |
| | M_val__L_c | L-Valine |
| | M_phe__L_c | L-Phenylalanine |
| | M_fmn_c | FMN |
| | M_gln__L_c | L-Glutamine |
| | M_ppi_c | Diphosphate |
| | M_amp_c | AMP |
| | M_gdp_c | GDP |
| | M_ala__L_c | L-Alanine |
| Module 36 | M_gly_c | Glycine |
| | M_ala__L_c | L-Alanine |

*continued on next page*

Table S3 – *continued from previous page*

| **Module** | **Metabolite ID** | **Metabolite Name** |
|---|---|---|
| | M_dcdp_c | DCDP |
| | M_met__L_c | L-Methionine |
| | M_prpp_c | 5-Phospho-alpha-D-ribose 1-diphosphate |
| | M_ile__L_c | L-Isoleucine |
| | M_clpn_c | cardiolipin |
| | M_k_c | Potassium |
| | M_glyc3p_c | Glycerol 3-phosphate |
| | M_udpg_c | UDP-glucose |
| | M_acald_c | Acetaldehyde |
| | M_s7p_c | Sedoheptulose 7-phosphate |
| | M_pg_c | phosphatidylglycerol |
| | M_tag_c | triacylglycerol |
| | M_xu5p__D_c | D-Xylulose 5-phosphate |
| | M_dgtp_c | DGTP |
| | M_dctp_c | DCTP |
| | M_mg2_c | Magnesium |
| | M_ade_c | Adenine |
| | M_nadp_c | NADP+ |
| Module 37 | M_nad_c | NAD+ |
| | M_cdpdag_c | CDP-diacylglycerol |
| | M_gtp_c | GTP |
| | M_mg2_c | Magnesium |
| | M_udpg_c | UDP-glucose |
| | M_r5p_c | D-ribose 5-phosphate |
| | M_pyr_c | Pyruvate |
| | M_ura_c | Uracil |

*continued on next page*

Table S3 – *continued from previous page*

| Module | Metabolite ID | Metabolite Name |
|---|---|---|
| | M_pg_c | phosphatidylglycerol |
| | M_ade_c | Adenine |
| | M_glyc3p_c | Glycerol 3-phosphate |
| | M_g3p_c | Glyceraldehyde 3-phosphate |
| | M_12dgr_c | 1,2-diacylglycerol |
| | M_sprm_c | spermine |
| | M_dctp_c | DCTP |
| | M_tag_c | triacylglycerol |
| | M_pap_c | Adenosine 3',5'-bisphosphate |
| | M_galfur12dgr_c | Gal-DAG |
| | M_r1p_c | D-ribose 1-phosphate |
| | M_fa_c | fatty acid |
| Module 38 | M_fmn_c | FMN |
| | M_r5p_c | D-ribose 5-phosphate |
| | M_coa_c | CoA |
| | M_ade_c | Adenine |
| | M_ump_c | UMP |
| | M_1ag3p_c | 1-acyl-glycerol 3-phosphate |
| | M_cdp_c | CDP |
| | M_datp_c | DATP |
| | M_adp_c | ADP |
| | M_nh3_c | NH3 |
| | M_pg_c | phosphatidylglycerol |
| | M_10fthfglu3_c | 10-formyltetrahydrofolate tri-L-glutamate |
| | M_sprm_c | spermine |
| | M_adn_c | Adenosine |

*continued on next page*

Table S3 – *continued from previous page*

| Module | Metabolite ID | Metabolite Name |
|---|---|---|
| | M_ura_c | Uracil |
| | M_uri_c | Uridine |
| | M_acald_c | Acetaldehyde |
| | M_cmp_c | CMP |
| | M_trnalys_c | TRNA(Lys) |
| | M_nac_c | Nicotinate |
| Module 39 | M_alatrna_c | L-Alanyl-tRNA(Ala) |
| | M_dadp_c | DADP |
| | M_pc_c | Phosphatidylcholine |
| | M_tyrtrna_c | L-Tyrosyl-tRNA(Tyr) |
| | M_cys__L_c | L-Cysteine |
| | M_trnacys_c | TRNA(Cys) |
| | M_dcyt_c | Deoxycytidine |
| | M_thmpp_c | thiamin diphosphate |
| | M_dcmp_c | DCMP |
| | M_dtdp_c | DTDP |
| | M_nadph_c | NADPH |
| | M_e4p_c | D-Erythrose 4-phosphate |
| | M_glytrna_c | Glycyl-tRNA(Gly) |
| | M_dctp_c | DCTP |
| | M_pro__L_c | L-Proline |
| | M_ile__L_c | L-Isoleucine |
| | M_val__L_c | L-Valine |
| | M_leu__L_c | L-Leucine |
| | M_ala__L_c | L-Alanine |
| | M_trnalys_c | TRNA(Lys) |

*continued on next page*

Table S3 – *continued from previous page*

| **Module** | **Metabolite ID** | **Metabolite Name** |
|---|---|---|
| Module 40 | M_asn__L_c | L-Asparagine |
| | M_gsn_c | Guanosine |
| | M_coa_c | CoA |
| | M_pc_c | Phosphatidylcholine |
| | M_dcyt_c | Deoxycytidine |
| | M_thmpp_c | thiamin diphosphate |
| | M_glyc3p_c | Glycerol 3-phosphate |
| | M_1ag3p_c | 1-acyl-glycerol 3-phosphate |
| | M_s7p_c | Sedoheptulose 7-phosphate |
| | M_ppi_c | Diphosphate |
| | M_lac__L_c | L-Lactate |
| | M_pap_c | Adenosine 3',5'-bisphosphate |
| | M_nac_c | Nicotinate |
| | M_udp_c | UDP |
| | M_adp_c | ADP |
| | M_atp_c | ATP |
| | M_ca2_c | Calcium |
| | M_ile__L_c | L-Isoleucine |
| | M_dgtp_c | DGTP |
| | M_tag_c | triacylglycerol |
| Module 41 | M_pep_c | Phosphoenolpyruvate |
| | M_gsn_c | Guanosine |
| | M_1ag3p_c | 1-acyl-glycerol 3-phosphate |
| | M_12dgr_c | 1,2-diacylglycerol |
| | M_g6p_c | D-Glucose 6-phosphate |
| | M_gtp_c | GTP |

*continued on next page*

Table S3 – *continued from previous page*

| **Module** | **Metabolite ID** | **Metabolite Name** |
|---|---|---|
| | M_ura_c | Uracil |
| | M_coa_c | CoA |
| | M_dctp_c | DCTP |
| | M_ile__L_c | L-Isoleucine |
| | M_nac_c | Nicotinate |
| | M_f6p_c | D-Fructose 6-phosphate |
| | M_thr__L_c | L-Threonine |
| | M_acald_c | Acetaldehyde |
| | M_nh3_c | NH3 |
| | M_xu5p__D_c | D-Xylulose 5-phosphate |
| | M_amp_c | AMP |
| | M_lac__L_c | L-Lactate |
| | M_ca2_c | Calcium |
| | M_nad_c | NAD+ |
| Module 42 | M_his__L_c | L-Histidine |
| | M_gly_c | Glycine |
| | M_pc_c | Phosphatidylcholine |
| | M_lystrna_c | L-lysyl-tRNA(Lys) |
| | M_f6p_c | D-Fructose 6-phosphate |
| | M_cys__L_c | L-Cysteine |
| | M_nadp_c | NADP+ |
| | M_arg__L_c | L-Arginine |
| | M_udpg_c | UDP-glucose |
| | M_e4p_c | D-Erythrose 4-phosphate |
| | M_nh3_c | NH3 |
| | M_k_c | Potassium |

*continued on next page*

Table S3 – *continued from previous page*

| **Module** | **Metabolite ID** | **Metabolite Name** |
|---|---|---|
| | M_s7p_c | Sedoheptulose 7-phosphate |
| | M_nac_c | Nicotinate |
| | M_glyc3p_c | Glycerol 3-phosphate |
| | M_mg2_c | Magnesium |
| | M_dtmp_c | DTMP |
| | M_prpp_c | 5-Phospho-alpha-D-ribose 1-diphosphate |
| | M_ppi_c | Diphosphate |
| | M_pep_c | Phosphoenolpyruvate |
| Module 43 | M_his__L_c | L-Histidine |
| | M_pc_c | Phosphatidylcholine |
| | M_thmpp_c | thiamin diphosphate |
| | M_gly_c | Glycine |
| | M_dcmp_c | DCMP |
| | M_arg__L_c | L-Arginine |
| | M_clpn_c | cardiolipin |
| | M_trnagln_c | TRNA(Gln) |
| | M_prpp_c | 5-Phospho-alpha-D-ribose 1-diphosphate |
| | M_trnalys_c | TRNA(Lys) |
| | M_nad_c | NAD+ |
| | M_gtp_c | GTP |
| | M_mettrna_c | L-methionyl-tRNA(Met) |
| | M_ca2_c | Calcium |
| | M_lac__L_c | L-Lactate |
| | M_sprm_c | spermine |
| | M_ile__L_c | L-Isoleucine |
| | M_udpgalfur_c | UDP-galactofuranose |

*continued on next page*

Table S3 – *continued from previous page*

| **Module** | **Metabolite ID** | **Metabolite Name** |
|---|---|---|
| | M_atp_c | ATP |
| | M_3pg_c | 3-Phospho-D-glycerate |
| Module 44 | M_asp__L_c | L-Aspartate |
| | M_pyr_c | Pyruvate |
| | M_nad_c | NAD+ |
| | M_ura_c | Uracil |
| | M_phe__L_c | L-Phenylalanine |
| | M_dgtp_c | DGTP |
| | M_cdpdag_c | CDP-diacylglycerol |
| | M_mg2_c | Magnesium |
| | M_accoa_c | Acetyl-CoA |
| | M_nadp_c | NADP+ |
| | M_r5p_c | D-ribose 5-phosphate |
| | M_adn_c | Adenosine |
| | M_pi_c | Phosphate |
| | M_glu__L_c | L-Glutamate |
| | M_gtp_c | GTP |
| | M_sprm_c | spermine |
| | M_acald_c | Acetaldehyde |
| | M_pg_c | phosphatidylglycerol |
| | M_tag_c | triacylglycerol |
| | M_g3p_c | Glyceraldehyde 3-phosphate |
| Module 45 | M_his__L_c | L-Histidine |
| | M_ca2_c | Calcium |
| | M_gly_c | Glycine |
| | M_cys__L_c | L-Cysteine |

*continued on next page*

Table S3 – *continued from previous page*

| **Module** | **Metabolite ID** | **Metabolite Name** |
|---|---|---|
| | M_gmp_c | GMP |
| | M_dctp_c | DCTP |
| | M_cmp_c | CMP |
| | M_ppi_c | Diphosphate |
| | M_10fthfglu3_c | 10-formyltetrahydrofolate tri-L-glutamate |
| | M_gsn_c | Guanosine |
| | M_k_c | Potassium |
| | M_xu5p__D_c | D-Xylulose 5-phosphate |
| | M_gtp_c | GTP |
| | M_coa_c | CoA |
| | M_dgtp_c | DGTP |
| | M_s7p_c | Sedoheptulose 7-phosphate |
| | M_nac_c | Nicotinate |
| | M_nh3_c | NH3 |
| | M_udpgalfur_c | UDP-galactofuranose |
| | M_dump_c | DUMP |
| Module 46 | M_ala__L_c | L-Alanine |
| | M_nadp_c | NADP+ |
| | M_s7p_c | Sedoheptulose 7-phosphate |
| | M_adn_c | Adenosine |
| | M_lac__L_c | L-Lactate |
| | M_ser__L_c | L-Serine |
| | M_dump_c | DUMP |
| | M_ctp_c | CTP |
| | M_pi_c | Phosphate |
| | M_cmp_c | CMP |

*continued on next page*

Table S3 – *continued from previous page*

| Module | Metabolite ID | Metabolite Name |
|---|---|---|
| | M_k_c | Potassium |
| | M_ca2_c | Calcium |
| | M_ru5p__D_c | D-Ribulose 5-phosphate |
| | M_fdp_c | D-Fructose 1,6-bisphosphate |
| | M_glu__L_c | L-Glutamate |
| | M_g3p_c | Glyceraldehyde 3-phosphate |
| | M_dcdp_c | DCDP |
| | M_ump_c | UMP |
| | M_r1p_c | D-ribose 1-phosphate |
| | M_udpg_c | UDP-glucose |
| Module 47 | M_dump_c | DUMP |
| | M_pap_c | Adenosine 3',5'-bisphosphate |
| | M_2pg_c | D-Glycerate 2-phosphate |
| | M_pi_c | Phosphate |
| | M_mg2_c | Magnesium |
| | M_udp_c | UDP |
| | M_nadph_c | NADPH |
| | M_pyr_c | Pyruvate |
| | M_asp__L_c | L-Aspartate |
| | M_r1p_c | D-ribose 1-phosphate |
| | M_accoa_c | Acetyl-CoA |
| | M_pg_c | phosphatidylglycerol |
| | M_13dpg_c | 3-Phospho-D-glyceroyl phosphate |
| | M_s7p_c | Sedoheptulose 7-phosphate |
| | M_ura_c | Uracil |
| | M_nadp_c | NADP+ |

*continued on next page*

Table S3 – *continued from previous page*

| **Module** | **Metabolite ID** | **Metabolite Name** |
|---|---|---|
| | M_thr__L_c | L-Threonine |
| | M_gdp_c | GDP |
| | M_ca2_c | Calcium |
| | M_cys__L_c | L-Cysteine |
| Module 48 | M_ru5p__D_c | D-Ribulose 5-phosphate |
| | M_dtdp_c | DTDP |
| | M_dump_c | DUMP |
| | M_sm_c | Sphingomyelin |
| | M_clpn_c | cardiolipin |
| | M_gsn_c | Guanosine |
| | M_thr__L_c | L-Threonine |
| | M_cdp_c | CDP |
| | M_1ag3p_c | 1-acyl-glycerol 3-phosphate |
| | M_arg__L_c | L-Arginine |
| | M_gmp_c | GMP |
| | M_ser__L_c | L-Serine |
| | M_prpp_c | 5-Phospho-alpha-D-ribose 1-diphosphate |
| | M_gtp_c | GTP |
| | M_chsterol_c | cholesterol |
| | M_coa_c | CoA |
| | M_dnad_c | Deamino-NAD+ |
| | M_udpgalfur_c | UDP-galactofuranose |
| | M_gly_c | Glycine |
| | M_trnalys_c | TRNA(Lys) |
| Module 49 | M_chsterol_c | cholesterol |
| | M_g6p_c | D-Glucose 6-phosphate |

*continued on next page*

Table S3 – *continued from previous page*

| **Module** | **Metabolite ID** | **Metabolite Name** |
|---|---|---|
| | M_gly_c | Glycine |
| | M_xu5p__D_c | D-Xylulose 5-phosphate |
| | M_clpn_c | cardiolipin |
| | M_lystrna_c | L-lysyl-tRNA(Lys) |
| | M_tyr__L_c | L-Tyrosine |
| | M_ca2_c | Calcium |
| | M_glyc3p_c | Glycerol 3-phosphate |
| | M_gsn_c | Guanosine |
| | M_pc_c | Phosphatidylcholine |
| | M_g3p_c | Glyceraldehyde 3-phosphate |
| | M_atp_c | ATP |
| | M_pyr_c | Pyruvate |
| | M_datp_c | DATP |
| | M_val__L_c | L-Valine |
| | M_trnalys_c | TRNA(Lys) |
| | M_glu__L_c | L-Glutamate |
| | M_k_c | Potassium |
| | M_pg_c | phosphatidylglycerol |
| Module 50 | M_nadph_c | NADPH |
| | M_sm_c | Sphingomyelin |
| | M_cdp_c | CDP |
| | M_13dpg_c | 3-Phospho-D-glyceroyl phosphate |
| | M_lys__L_c | L-Lysine |
| | M_pc_c | Phosphatidylcholine |
| | M_udpgalfur_c | UDP-galactofuranose |
| | M_arg__L_c | L-Arginine |

*continued on next page*

Table S3 – *continued from previous page*

| Module | Metabolite ID | Metabolite Name |
|---|---|---|
| | M_dcdp_c | DCDP |
| | M_xu5p__D_c | D-Xylulose 5-phosphate |
| | M_gmp_c | GMP |
| | M_1ag3p_c | 1-acyl-glycerol 3-phosphate |
| | M_gln__L_c | L-Glutamine |
| | M_gtp_c | GTP |
| | M_glyc3p_c | Glycerol 3-phosphate |
| | M_dtdp_c | DTDP |
| | M_pi_c | Phosphate |
| | M_leu__L_c | L-Leucine |
| | M_r1p_c | D-ribose 1-phosphate |
| | M_gsn_c | Guanosine |

Table S4: Gene functional modules.

| Module | Gene ID | Gene Product |
|---|---|---|
| Module 1 | G_0518 | Lipoprotein signal peptidase |
| | G_0332 | Uncharacterized protein |
| | G_0593 | Uncharacterized protein |
| | G_0080 | Uncharacterized protein |
| | G_0610 | DNA-formamidopyrimidine glycosylase |
| | G_0389 | Uncharacterized protein |
| | G_0548 | tRNA (cytidine(34)-2'-O)-methyltransferase |
| | G_0549 | Non-canonical purine NTP pyrophosphatase |
| | G_0517 | Uncharacterized RNA pseudouridine synthase |
| | G_0609 | Chromosome replication initiation protein |
| | G_0693 | Uncharacterized protease |
| | G_0329 | Uncharacterized pseudouridine synthase |
| | G_0612 | DNA polymerase III subunit alpha |
| | G_0076 | Asparagine–tRNA ligase |
| | G_0710 | Low specificity hydrolase |
| | G_0106 | Exodeoxyribonuclease VII large subunit |
| | G_0519 | Isoleucine–tRNA ligase |
| | G_0379 | Uncharacterized protein |
| | G_0600 | Ribonuclease J |
| | G_0611 | DNA polymerase I |
| Module 2 | G_0910 | 50S ribosomal protein L34 |
| | G_0363 | 16S rRNA processing protein |
| | G_0503 | Uncharacterized protein |
| | G_0877 | Riboflavin ECF transporter S component |
| | G_0010 | Nucleoside ABC transporter ATP-binding protein |

*continued on next page*

Table S4 – *continued from previous page*

| Module | Gene ID | Gene Product |
|---|---|---|
| | G_0006 | DNA gyrase subunit B |
| | G_0886 | Proton-glutamate symporter |
| | G_0494 | N-acetylmannosamine-6-phosphate 2-epimerase |
| | G_0005 | Uncharacterized protein |
| | G_0500 | Maturation protease for ribosomal protein L27 |
| | G_0011 | Nucleoside ABC transporter substrate-binding protein |
| | G_0094 | Uncharacterized protein |
| | G_0544 | Heat-inducible transcription repressor |
| | G_0909 | Ribonuclease P protein component |
| | G_0081 | tRNA uridine(34) 5-carboxymethylaminomethyl synthesis GTPase |
| | G_0009 | Nucleoside ABC transporter permease |
| | G_0606 | Phosphoglycerate kinase |
| | G_0604 | Uncharacterized protein |
| | G_0600 | Ribonuclease J |
| | G_0366 | L16-binding dependent 50S subunit-maturation GTPase |
| Module 3 | G_0831 | Phosphoribosylpyrophosphate synthetase |
| | G_0610 | DNA-formamidopyrimidine glycosylase |
| | G_0332 | Uncharacterized protein |
| | G_0612 | DNA polymerase III subunit alpha |
| | G_0106 | Exodeoxyribonuclease VII large subunit |
| | G_0830 | Uncharacterized protein |
| | G_0438 | Uncharacterized protein |

*continued on next page*

Table S4 – *continued from previous page*

| **Module** | **Gene ID** | **Gene Product** |
|---|---|---|
| | G_0329 | Uncharacterized pseudouridine synthase |
| | G_0080 | Uncharacterized protein |
| | G_0328 | Chromosome segregation protein B |
| | G_0389 | Uncharacterized protein |
| | G_0437 | Putative 3'-5' exoribonuclease |
| | G_0379 | Uncharacterized protein |
| | G_0338 | Uncharacterized lipoprotein |
| | G_0611 | DNA polymerase I |
| | G_0832 | Aminoacyl-tRNA hydrolase |
| | G_0387 | tRNA 2-thiouridine(34) synthase |
| | G_0443 | 5-Formyltetrahydrofolate cyclo-ligase |
| | G_0451 | Glyceraldehyde-3-phosphate dehydrogenase |
| | G_0609 | Chromosome replication initiation protein |
| Module 4 | G_0779 | PTS sugar transporter |
| | G_0908 | Membrane protein insertase |
| | G_0415 | Chromosome segregation protein |
| | G_0775 | Ribonuclease R |
| | G_0361 | 23S rRNA (pseudouridine(1915)-N3)-methyltransferase |
| | G_0517 | Uncharacterized RNA pseudouridine synthase |
| | G_0352 | Uncharacterized protein |
| | G_0826 | DNA polymerase III subunit delta |
| | G_0438 | Uncharacterized protein |
| | G_0388 | Uncharacterized protein |
| | G_0819 | Thioredoxin-disulfide reductase |
| | G_0433 | Uncharacterized protein |

*continued on next page*

Table S4 – *continued from previous page*

| **Module** | **Gene ID** | **Gene Product** |
|---|---|---|
| | G_0437 | Putative 3'-5' exoribonuclease |
| | G_0824 | Excinuclease ABC subunit A |
| | G_0907 | Low specificity hydrolase |
| | G_0416 | Uncharacterized protein |
| | G_0601 | Uncharacterized protein |
| | G_0821 | HPr(Ser) kinase/phosphatase |
| | G_0418 | Ribonuclease III |
| | G_0047 | DNA polymerase III subunit gamma and tau |
| Module 5 | G_0779 | PTS sugar transporter |
| | G_0415 | Chromosome segregation protein |
| | G_0388 | Uncharacterized protein |
| | G_0775 | Ribonuclease R |
| | G_0776 | SsrA-binding protein |
| | G_0789 | F0F1 ATP synthase subunit delta/epsilon |
| | G_0517 | Uncharacterized RNA pseudouridine synthase |
| | G_0361 | 23S rRNA (pseudouridine(1915)-N3)-methyltransferase |
| | G_0908 | Membrane protein insertase |
| | G_0416 | Uncharacterized protein |
| | G_0389 | Uncharacterized protein |
| | G_0413 | Adenine phosphoribosyltransferase |
| | G_0414 | Guanosine-3',5'-bis(diphosphate) 3'-pyrophosphohydrolase |
| | G_0418 | Ribonuclease III |
| | G_0438 | Uncharacterized protein |
| | G_0379 | Uncharacterized protein |

*continued on next page*

Table S4 – *continued from previous page*

| **Module** | **Gene ID** | **Gene Product** |
|---|---|---|
| | G_0778 | Uncharacterized protein |
| | G_0433 | Uncharacterized protein |
| | G_0819 | Thioredoxin-disulfide reductase |
| | G_0791 | F0F1 ATP synthase subunit gamma |
| Module 6 | G_0285 | Elongation factor 4 |
| | G_0196 | Spermidine/putrescine ABC transporter permease |
| | G_0910 | 50S ribosomal protein L34 |
| | G_0819 | Thioredoxin-disulfide reductase |
| Module 7 | G_0887 | Coenzyme A Disulfide Reductase |
| | G_0419 | Phosphate acyltransferase |
| | G_0137 | 50S ribosomal protein L31 |
| | G_0402 | rRNA maturation RNase |
| | G_0005 | Uncharacterized protein |
| | G_0710 | Low specificity hydrolase |
| | G_0167 | Oligopeptide ABC transporter ATP-binding protein |
| | G_0434 | 23S rRNA (uridine(1939)-m5)-methyltransferase |
| | G_0010 | Nucleoside ABC transporter ATP-binding protein |
| | G_0430 | Uncharacterized DNA-binding protein |
| | G_0379 | Uncharacterized protein |
| | G_0910 | 50S ribosomal protein L34 |
| | G_0429 | Signal recognition particle-docking protein |
| | G_0164 | Uncharacterized protein |

*continued on next page*

Table S4 – *continued from previous page*

| **Module** | **Gene ID** | **Gene Product** |
|---|---|---|
| Module 8 | G_0415 | Chromosome segregation protein |
| | G_0416 | Uncharacterized protein |
| | G_0419 | Phosphate acyltransferase |
| | G_0418 | Ribonuclease III |
| | G_0285 | Elongation factor 4 |
| | G_0798 | Uracil phosphoribosyltransferase |
| | G_0290 | tRNA pseudouridine(55) synthase |
| | G_0414 | Guanosine-3',5'-bis(diphosphate) 3'-pyrophosphohydrolase |
| | G_0796 | F0F1 ATP synthase subunit A |
| | G_0604 | Uncharacterized protein |
| | G_0291 | FAD synthetase |
| | G_0644 | 50S ribosomal protein L17 |
| | G_0046 | Recombination protein |
| | G_0648 | 50S ribosomal protein L36 |
| | G_0910 | 50S ribosomal protein L34 |
| | G_0642 | ECF transporter ATPase |
| | G_0378 | NAD(+) synthase |
| | G_0646 | 30S ribosomal protein S11 |
| | G_0289 | Ribosome-binding factor A |
| | G_0264 | Uncharacterized serine/threonine protein kinase |
| Module 9 | G_0338 | Uncharacterized lipoprotein |
| | G_0419 | Phosphate acyltransferase |
| | G_0330 | Deoxyguanosine kinase |

*continued on next page*

Table S4 – *continued from previous page*

| **Module** | **Gene ID** | **Gene Product** |
|---|---|---|
| | G_0034 | Uncharacterized efflux ABC transporter permease |
| | G_0412 | Uncharacterized protein |
| | G_0416 | Uncharacterized protein |
| | G_0413 | Adenine phosphoribosyltransferase |
| | G_0106 | Exodeoxyribonuclease VII large subunit |
| | G_0415 | Chromosome segregation protein |
| | G_0418 | Ribonuclease III |
| Module 10 | G_0010 | Nucleoside ABC transporter ATP-binding protein |
| | G_0910 | 50S ribosomal protein L34 |
